# Minute-scale oscillatory sequences in medial entorhinal cortex

**DOI:** 10.1101/2022.05.02.490273

**Authors:** Soledad Gonzalo Cogno, Horst A. Obenhaus, R. Irene Jacobsen, Flavio Donato, May-Britt Moser, Edvard I. Moser

**Author notes:** Equal contribution (Flavio Donato, Edvard I. Moser). Corresponding authors: Soledad Gonzalo Cogno,; Flavio Donato,; May-Britt Moser,; Edvard I. Moser,; Editorial correspondence.

## Abstract

The medial entorhinal cortex (MEC) hosts many of the brain’s circuit elements for spatial navigation and episodic memory, operations that require neural activity to be organized across long durations of experience^1^. While location is known to be encoded by a plethora of spatially tuned cell types in this brain region^2–6^, little is known about how the activity of entorhinal cells is tied together over time. Among the brain’s most powerful mechanisms for neural coordination are network oscillations, which dynamically synchronize neural activity across circuit elements^7–10^. In MEC, theta and gamma oscillations provide temporal structure to the neural population activity at subsecond time scales^1,11–13^. It remains an open question, however, whether similarly powerful coordination occurs in MEC at behavioural time scales, in the second-to-minute regime. Here we show that MEC activity can be organized into a minute-scale oscillation that entrains nearly the entire cell population, with periods ranging from 10 to 100 seconds. Throughout this ultraslow oscillation, neural activity progresses in periodic and stereotyped sequences. This activity was elicited while mice ran at free pace on a rotating wheel in darkness, with no change in its location or running direction and no scheduled rewards. The oscillation sometimes advanced uninterruptedly for tens of minutes, transcending epochs of locomotion and immobility. Similar oscillatory sequences were not observed in neighboring parasubiculum or in visual cortex. The ultraslow oscillation of activity sequences in MEC may have the potential to couple its neurons and circuits across extended time scales and to serve as a scaffold for processes that unfold at behavioural time scales, such as navigation and episodic memory formation.

## Main

Brain function emerges from the dynamic coordination of interconnected neurons^9,10, 14–16^. At subsecond time scales, cells are coordinated within and across dispersed brain regions by way of neuronal oscillations^7,9,13, 17–20^. Studies have reported oscillations also at slower time scales, with periods lasting from seconds to minutes, in individual neurons^21–26^ and in local field potentials^27–29^. It remains unknown, however, how pervasive these ultraslow oscillations are, whether they serve a role in neuronal coordination, and if they do, how the activity of participating neurons is organized in space and time across the neural circuit.

We directed our search for ultraslow oscillations to the medial entorhinal cortex (MEC), a brain circuit that by containing many of the elements for navigational behavior^2–6^ and episodic memory formation^1,30,31^, may possess mechanisms to organize neural activity at time scales of seconds to minutes. Ultraslow oscillations, if they exist in MEC, might structure neural activity over long time scales and interact with faster oscillations, such as theta and gamma rhythms, which are predominant in this brain area^11,12,32^. To maximize the detectability of ultraslow MEC oscillations and to rule out variations in external stimuli as sources of modulation, we monitored activity in hundreds of MEC cells with two-photon calcium imaging while head-fixed mice ran on a rotating wheel for 30 or 60 minutes^33–35^, in darkness and with no scheduled rewards^36,37^ (Fig. 1a).

**Figure 1.**
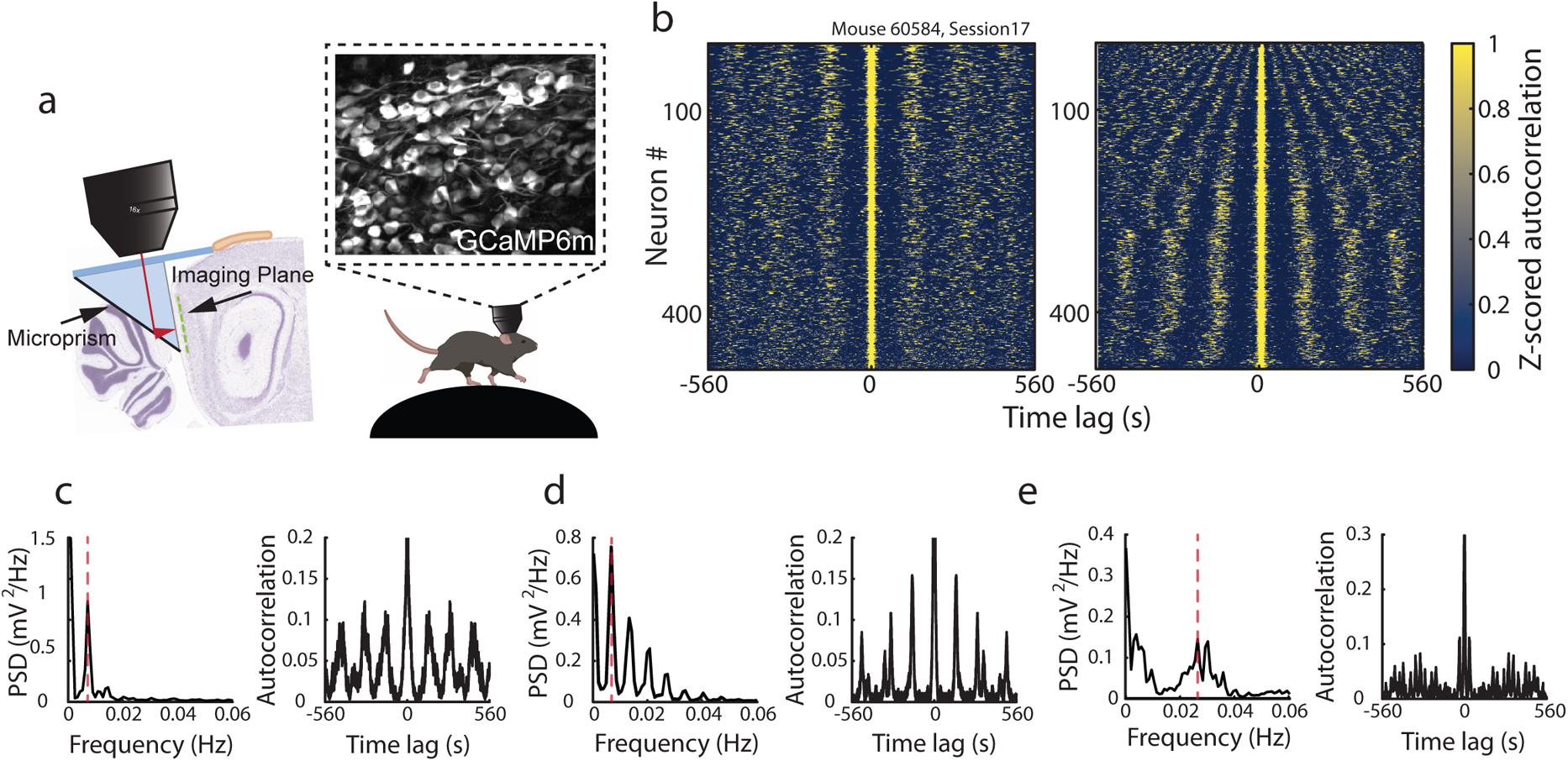
Ultraslow oscillations in calcium activity of MEC neurons. **a.** Schematic representation of the experimental set-up. Neural activity is monitored through a prism from GCaMP6m-expressing neurons of the medial entorhinal cortex (MEC) in head-fixed mice running in darkness on a non-motorized running wheel. Mice alternate freely between running and rest. **b.** Stacked autocorrelations of single-cell calcium activity for one example session (3600 s, or 1 h, of continuous recording, 484 neurons; session 17 from animal #60584.). Each row is the autocorrelation of one cell’s deconvolved and binarized calcium activity (subsequently referred to as the cell’s “calcium activity”), plotted as a function of time lag. Z-scored autocorrelations are color-coded. Left: Neurons are sorted according to the maximum power of the power spectral density (PSD) calculated on each autocorrelation separately, in a descending order. The vertical bands suggest that single cell calcium activity is periodic. Right: The same neurons sorted according to peak frequency in the PSD. The curved nature of the bands illustrates that while most cells exhibited slow oscillation, the frequency of the oscillation showed some variation across cells. **c.** PSD (left) calculated on the autocorrelation (right) of one example cell’s calcium activity. The dashed red line indicates the primary frequency at which the PSD peaks. The sole narrow peak at 0.0066 Hz is mirrored by the well-defined oscillatory pattern in the autocorrelation. **d.** As in (c) but for another example cell. The PSD peaks at 0.0066 Hz and has harmonics at 0.0132, 0.0207 and 0.0273 Hz. **e.** As in (c) but for another example cell in the same recording. The PSD peaks at 0.0038 Hz and 0.0264 Hz. Both peaks are much wider than in (c), corresponding to a weaker oscillatory pattern in the autocorrelation.

### Activity of MEC neurons undergoes ultraslow oscillations

Behavior on the rotating wheel was characterized by bouts of running, at variable speed and acceleration, interleaved with periods of rest (Extended data Fig. 2a). To determine if neural activity in MEC exhibits ultraslow oscillations in this task, for each recorded cell we deconvolved the calcium signals^38,39^ and binarized the obtained signals, using a bin size of 129 ms. This process yielded a deconvolved and binary calcium activity that had in each time bin a value of 1 in presence of calcium events, and 0 otherwise (“calcium activity” for the rest of the paper). For each cell, we calculated the autocorrelation of the calcium activity and the corresponding power spectral density (PSD). When the autocorrelation diagrams of all cells from one session were stacked into a matrix with cells as rows and time lags as columns, we observed vertical bands (Fig. 1b, left), suggesting that the calcium activity of individual cells was oscillatory and that many cells shared a similar oscillation frequency. When sorting the autocorrelations in the matrix according to the frequency at which the PSDs peaked (“primary frequency”), the spread of oscillatory frequencies became clearer (Fig. 1b, right). Some cells had only one prominent peak in their PSD (Fig 1c), suggesting that they were active at equidistant intervals throughout the session. Other cells had several peaks, often with the higher frequencies appearing as harmonics of a fundamental frequency, or they had wider peaks, indicating more variable activity intervals (Fig. 1d and e). In the example session in Fig. 1b, most cells (72%, 348 of 484) had a primary frequency lower than 0.01 Hz (44% of the cells had a primary frequency within the range 0.006-0.008 Hz), and there were no cells whose PSD peaked at frequencies higher than 0.1 Hz. In the complete data set (15 sessions over 5 animals), there was some variation in frequencies across sessions and animals but the primary frequency was always below 0.1 Hz (all 6231 cells; range of maximum frequencies across 15 sessions: 0.036-0.057 Hz). Taken together, these findings demonstrate ultraslow oscillations of single cell calcium activity in MEC, at periods in the order of tens of seconds to minutes.

### MEC population activity is organized into oscillatory sequences

To determine whether the ultraslow oscillation of calcium activity of different cells is coordinated and temporally structured at the neural population level, we introduced a two-step procedure. First, we stacked the calcium activity of all cells to produce a matrix that had as many rows as recorded cells, and as many columns as time bins (bin size 129 ms). Second, using time bins as data points, we calculated instantaneous correlations between the calcium activity of all pairs of cells, and used these values to sort the cells such that those that are nearby in the sorting are highly synchronized. The cell pair with the highest correlation value was identified and one of the two cells was defined as the “lead” cell. The remaining cells were sorted, in a descending manner, based on their correlation value with the lead cell.

When cells were sorted by correlations and their activity plotted across session time, we observed periodic sequences of neuronal activation (Fig. 2a). The sequences unfolded successively at a steady rate, with no interruption for tens of minutes. The periodic sequences indicate the presence of an oscillation at the neural population level that coordinates the order of activity among the neurons. Each cell was followed first by cells that were highly synchronous, and then successively by less synchronous cells until synchronization caught up again (Extended data Fig. 3a).

**Figure 2.**
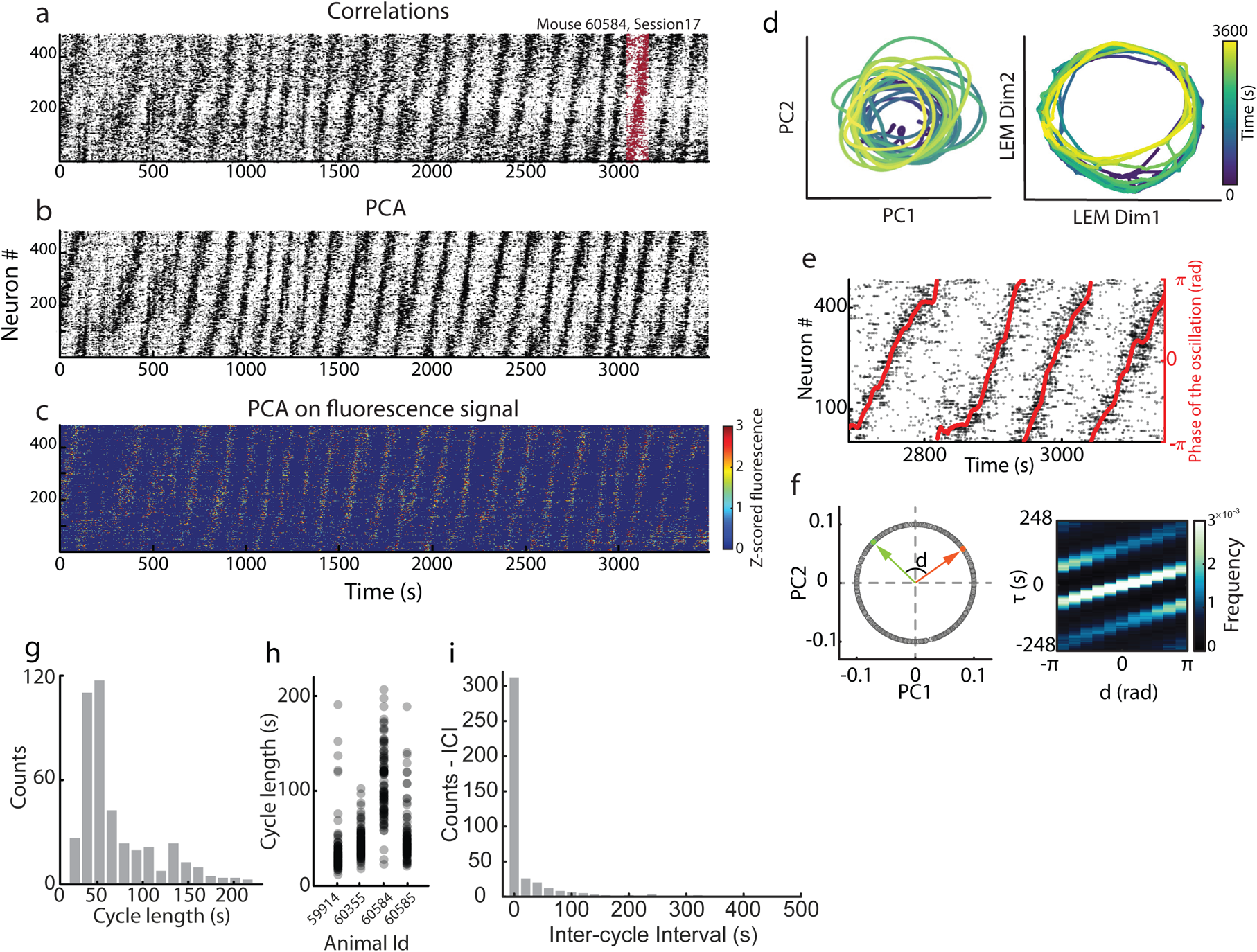
Ultraslow oscillations in MEC consist of neuronal sequences. **a.** Raster plot representation of the matrix of calcium activity obtained after stacking the calcium activity of all cells recorded in one experimental session (same as in Fig. 1b). Each row of the raster plot shows the calcium activity of one neuron plotted as a function of time (in seconds, bin size 129 ms). Time bins with calcium events are indicated with black dots. Time bins with no calcium events are white. Neurons are sorted according to the correlation between the calcium activity. The sorting revealed sequences of neuronal activity. One example sequence is indicated in red. Notice the slow temporal scale of the sequences (121 s for the highlighted sequence). **b.** As in (a) but now with neurons sorted according to the PCA method, where we calculated for each cell the arctangent of the ratio between the cell’s loading on principal component 2 (PC2) and PC1, and then sorted the cells according to those values in a descending manner. **c.** As in (b) but showing the fluorescence calcium signals instead of the deconvolved calcium activity. Z-scored calcium signals are color-coded. Neurons are sorted according to the PCA method. **d.** Projection of neural activity of the session presented in (a-c) onto a low-dimensional embedding defined by the first two principal components of PCA (left), and by the first two dimensions of a LEM analysis (right). Time is color-coded. Neural trajectories are circular, with population activity propagating along a ring-shaped manifold. One full rotation of the population activity along the ring-shaped manifold is defined as a “cycle” of the population oscillation. **e.** Raster plot as in (b), with the phase of the oscillation overlaid in red (right y axis: phase of the oscillation in radians). **f.** Left: Distance *d* between two neurons in the PCA sorting is calculated as the difference between the angles of the vectors defined by the loadings of each neuron on PC1 and PC2 with respect to PC1. The schematic shows the distance between two neurons, one in orange and the other in green. The length of the vectors is disregarded in this quantification. Right: Joint distribution of the time lag *τ* that maximizes the cross-correlation between the calcium activity of any given pair of neurons and their distance *d* in the PCA sorting. Color code: normalized frequency, each count is a cell pair. The increasing relationship between *τ* and *d* indicates sequential organization of neural activity. **g.** Distribution of cycle lengths across 15 oscillatory sessions over 5 animals (one animal did not have detectable oscillations, 421 cycles in total). Each count is an individual cycle. **h.** Cycle lengths shown separately for each animal with oscillations (421 cycles in total). For each animal all oscillatory sessions were pooled. Cycle length was heterogenous across sessions and animals. **i.** Distribution of inter-cycle intervals (ICI; 406 ICIs in total across 15 oscillatory sessions). Each count is an ICI. During uninterrupted oscillations the ICI is 0.

While sorting cells according to their correlation values unveiled recurring sequences of activity, computing correlations from the binarized calcium activity (or the raw calcium signals) can be inherently noisy due to variability in the frequency of deconvolved calcium events and dependence on fine tuning of hyperparameters such as the size of the kernel used to smooth the calcium activity. Thus, we sought a sorting approach that did not rely on hyperparameters. We leveraged the fact that sequences of activity constitute low-dimensional dynamics with intrinsic dimensionality equal to 1, and adopted an unsupervised approach based on dimensionality reduction^40^ to sort the cells. For each recording session we applied principal component analysis (PCA) to the full matrix of calcium activity, including all epochs of movement and immobility. We kept the first two principal components (PCs), which is the minimum number of components needed to embed non-linear 1-dimensional dynamics. Expecting that the ordering in cell activation would be expressed in the relationship between the cells’ loadings on the two PCs, we measured for each cell the angle *8* ∈ [*−π*, *π)* of the vector defined by the pair of loadings on PC1 and PC2, and then sorted the neurons based on these angles in a descending manner (Extended data Fig. 3b). Sorting the cells in this way (“PCA method”) revealed the same stereotyped sequences of neuronal activation that we previously found through correlations among cell pairs (Fig. 2b); however, the sequential organization was now more salient. Sequences proceeded uninterruptedly in a periodic manner and seemed to involve the majority of the recorded MEC cells. We will refer to these oscillatory sequences as the “population oscillation” to distinguish them from oscillations in single cell calcium activity. The population oscillation was not present if cells were not sorted or if the PCA method was applied to matrices of calcium activity in which the calcium events were temporally shuffled (Extended data Fig. 3c, first and second row on the left). The population oscillation was similarly observed when neurons were sorted according to unsupervised methods relying on a variety of non-linear dimensionality reduction techniques (Extended data Fig. 3c, third row on the left, and second column), and also when the neurons’ calcium activity was visualized by the unprocessed calcium signals (Fig. 2c), suggesting that it is not an artifact of the spike deconvolution.

Since sequences of neural activity constitute low-dimensional dynamics, we next asked what is the topology of the underlying manifold. First, we visualized the manifold by projecting the population activity onto the first two PCs. The manifold resembled a ring, along which the population activity propagated repeatedly with periodic boundary conditions (Fig. 2d left, Extended data Fig. 3d). However, because the ring-shaped manifold could be lying on a curved surface, in which case the linear embedding of PCA might result in distortions when the manifold is visualized, we next adopted a non-linear dimensionality reduction method. A Laplacian Eigenmap (LEM) approach was chosen because it has a lower number of hyperparameters as compared to the other non-linear techniques (Extended data Fig. 3c). We applied LEM to the matrix of calcium activity and then projected the population activity onto the 2-dimensional embedding spanned by the first two LEM dimensions. The manifold still had the shape of a ring, as previously suggested by the PCA projection, although now the circular pattern was more salient (Fig. 2d right). The progression of population activity along the manifold was tracked through a parameter that we call the “phase of the oscillation” and that we calculate as the arctangent of the ratio between the population activity modes given by PC2 and PC1 (Fig. 2e, red trace). During one full rotation of the population activity along the ring-shaped manifold, which we refer to as one “cycle”, the phase of the oscillation traversed [*−π*, *π*) rad.

While striking population oscillations were observed across multiple sessions and animals, the population activity exhibited considerable variability, ranging from non-patterned activity to highly stereotypic and periodic sequences (Extended data Fig. 4a). This variability was also observed when examining the joint distribution of time lags (*τ*) that maximized the correlation between cells’ calcium activity and the angular distances *d* in the PCA sorting (Fig. 2f left). In sessions with clear population oscillations, the time lag *τ* increased with the distance *d*, which indicated sequential activation of neural activity. This dependence was observed a discrete number of times in each session, which indicated that cells were active periodically and at a fixed frequency or at an integer multiple of it (Fig. 2f right, built on the example session shown in Fig. 2a, and Extended data Fig. 4b top for another example with a different time scale). In sessions without detectable population oscillations such structure was not observed (Extended data Fig. 4b bottom). This variability in population dynamics prompted us to quantify, for each session, the extent to which the population activity was oscillatory through an oscillation score that ranged from 0 (no oscillation) to 1 (strong oscillations). The score was calculated by first binning the angular distance between cells in the PCA sorting, using 11 bins, and then counting the fraction of bins in which the cross correlations between the calcium activity of cell pairs peaked at regular intervals (Online Methods). The distribution of oscillation scores over the entire MEC dataset (27 recording sessions in 5 mice) was bimodal (Extended data Fig. 4c), with 12/27 sessions exhibiting scores between 0 and 0.2 (no oscillations), and 15/27 sessions scoring between 0.72 and 1. The latter sessions were classified as oscillatory (15 oscillatory sessions over 5 mice; one of the mice did not have any oscillatory sessions; Extended data Fig. 4a).

**Figure 4.**
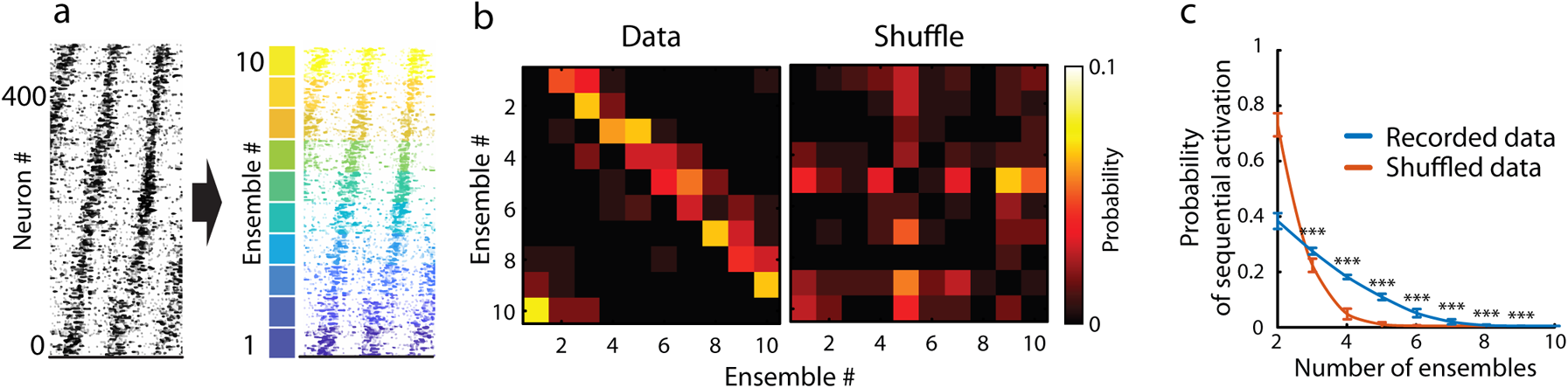
The population oscillation consists of unidirectional periodic activity sequences. **a.** Schematic of the process for splitting neurons into ensembles of co-active cells. Neurons sorted according to the PCA method are allocated to 10 equally sized ensembles (color-coded). **b.** Left: Matrix of transition probabilities between pairs of ensembles at consecutive time points. Data are from the example session in Fig. 2a (bin size = 15.12 s). Right: Same as left panel but for one shuffle realization. Transition probabilities are color coded. In the left diagram, note the higher probability of transitions between consecutive ensembles (increased probabilities near the diagonal), the directionality of transitions (increased probabilities above diagonal) and the periodic boundary conditions in ensemble activation (presence of transitions from ensemble 10 to ensemble 1). **c.** Probability of sequential ensemble activation as a function of the number of ensembles that are sequentially activated (mean ± S.D.; For 3-9 ensembles: *n* = 15 oscillatory sessions, 7500 shuffle realizations, *p ≤ 5*.4 × 10^−11^, range of *Z* values: 6.45 to 59.18, one-tailed Wilcoxon rank-sum test). Blue, recorded data; orange, shuffled data. For each session, the probability of sequential ensemble activation was calculated over 500 shuffled realizations, and shuffled realizations were pooled across sessions.

For each oscillatory session, we investigated how the population activity varied across individual cycles and whether the length of individual cycles varied within and between sessions. We identified individual cycles by extracting the subset of adjacent time bins where the phase of the oscillation increased smoothly within the range [-π,π) (bin size 129 ms, Extended data Fig. 5a,b). We divided each cycle into 10 segments, and for each segment we calculated the mean rate of calcium events over recorded neurons (total number of events across cells divided by cycle segment duration and number of cells). Across sessions we found that the percentage rate change from the segment with the minimum event rate to the segment with the maximum rate was no more than 18% (Extended data Fig. 5c). There was no significant difference in median event rate between pairs of segments within cycles (Extended data Fig. 5c). Next we quantified, within and across sessions and animals, the variability in the length of individual cycles, or the time for population activity to traverse the ring once (Extended data Fig. 5a). Cycle lengths ranged from tens of seconds to minutes (Fig. 2g) and showed high variability across sessions and animals (Fig. 2h, Extended data Fig. 5d) but there was little variability within individual sessions (Extended data Fig. 5a,e). Cycle length was independent of the number of recorded cells (Extended data Fig. 5f).

**Figure 3.**
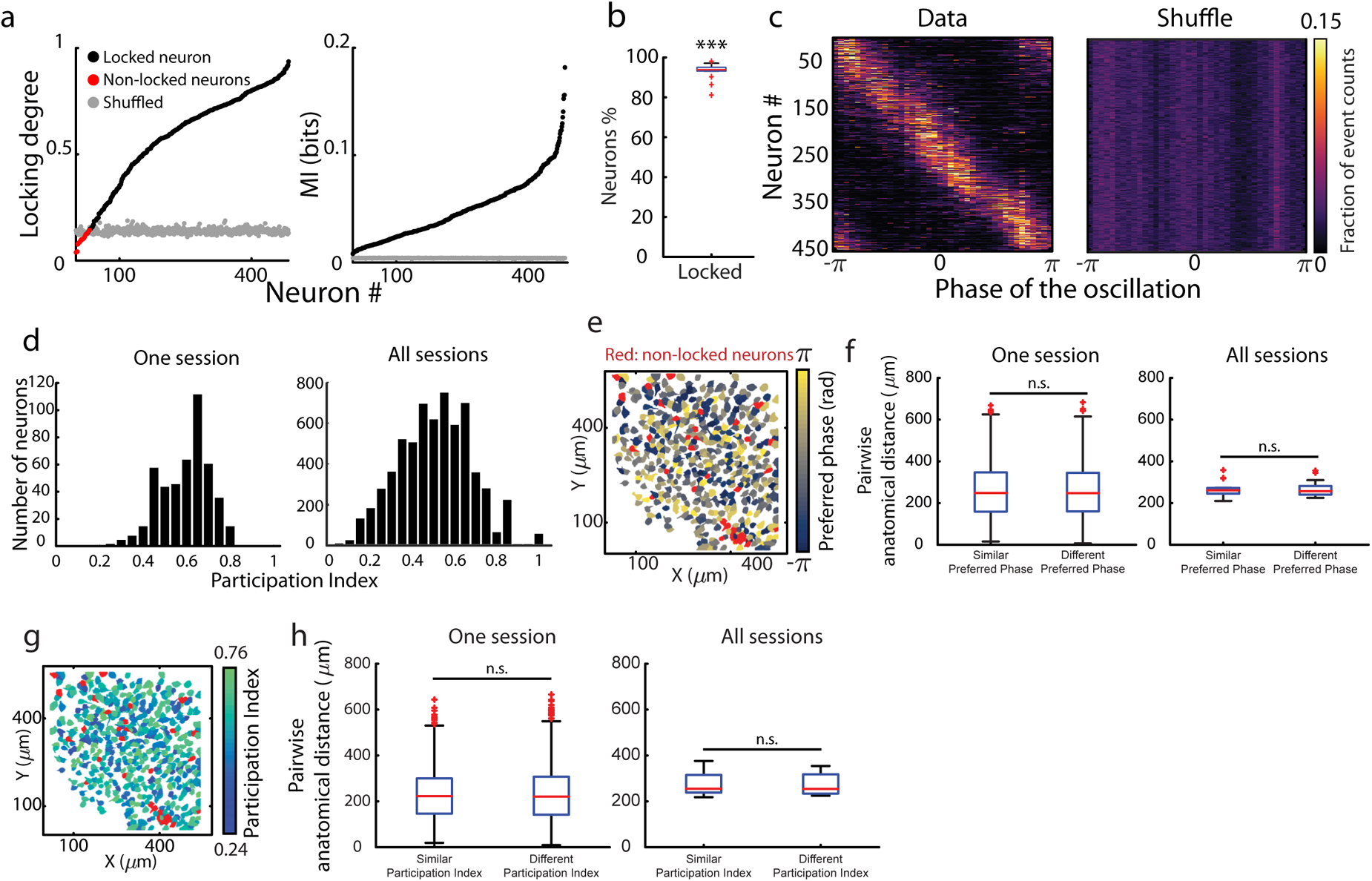
Nearly all MEC neurons are locked to the population oscillation. **a.** Left: Distribution of locking degrees for all imaged neurons in the example session in Fig. 2a. The locking degree, computed as the length of the mean vector over the distribution of phases at which calcium events occurred, takes values between 0 (absence of locking) and 1 (perfect locking). Black dots indicate locked neurons, red dots non-locked neurons, grey dots the 99^th^ percentile of the null distribution used to assess locking. For locked cells the locking degree is larger than the 99^th^ percentile of the null distribution (458 of 484 cells were locked to the phase of the oscillation). Neurons are sorted according to their looking degree in an ascending manner. Bin size = 129 ms. Right: Distribution of values of mutual information (MI, in bits) between the phase of the oscillation and the counts of calcium events (“event counts”) for all imaged neurons in the example session in Fig. 2a. Black dots indicate the values of MI and grey dots the estimated bias in the MI. For all cells the MI is larger than the bias. Neurons are sorted according to their MI value in an ascending manner. Bin size = 0.52 s. **b.** Box plot showing percentage of locked neurons over all sessions (median = 94%; one sample t-test for a null hypothesis of 50% locked and non-locked cells, *n*=15 oscillatory sessions, *p* = 1 × 10^−15^, *t* =38.6). Red line indicates median across sessions, bottom and top lines in blue indicate lower and upper quartiles, respectively. The length of the whiskers indicates 1.5 times the interquartile range. Red crosses show outliers exceeding 1.5 times the interquartile range. *** *p* < 0.001, ** *p* < 0.01, * *p* < 0.05, n.s. *p* > 0.05. **c.** Tuning of single cell calcium activity to the phase of the oscillation. Left: Each row indicates the tuning curve of one locked neuron of the example session in Fig. 2a (n = 458 locked cells). Right: Same as left, but now for the tuning curves obtained in one shuffle realization of the data in which the calcium events were temporally shuffled. Tuning curves were calculated by determining the fraction of event counts across phase bins of the oscillation (bin size~0.16 rad, 40 bins in total). Tuning curves are color coded. **d.** Left: Distribution of participation indexes across neurons in the example session shown in Fig. 2a (n = 484 cells). The participation index (PI) quantifies the extent to which a cell’s calcium activity is distributed across all cycles of the population oscillation, or rather concentrated in a few cycles, regardless of its locking degree. PI was calculated for each cell separately as the number of cycles needed to account for 90% of the total number of event counts. Right: Distribution of participation indexes across all 15 oscillatory sessions (n = 6231 cells). Each count in each of the plots is a neuron. **e.** Anatomical distribution of neurons in the field of view (FOV) of the example session in Fig. 2a. The preferred phase of each neuron, calculated as the mean phase at which the calcium events occurred, is color-coded. Neurons in red are not significantly locked to the phase of the oscillation. The preferred phases are anatomically intermingled. Dorsal MEC on top, medial on the right, as in Extended data Fig. 1. **f.** Left: Box plot of pairwise anatomical distances between cells with similar preferred phase (each cell in the pair has a preferred phase ~ 0 rad) or different preferred phase (one cell in the pair has a preferred phase ~ 0, and the other one a preferred phase ~ π rad). Data are for the example session in Fig. 2a (*n* = 990 distances in the similar group, 2025 distances in the different group, *p* = 0.65, *Z* = 0.46, Wilcoxon rank-sum test). Right: Similar to the left panel but for all 15 oscillatory sessions, including the example session in the left panel (*p* = 0.80, *Z* = 0.25, Wilcoxon rank-sum test). A fraction of 10% of the total number of locked cells was used to define the groups with preferred phase ~ 0 rad or ~π rad. Symbols as in Fig. 3b. **g.** Same as (e) but for the participation index. Note that also the PIs are anatomically intermingled. **h.** Similar to (f) but for the participation index. Left: Box plot of pairwise anatomical distances between cells with similar or different participation indexes for the example session in Fig. 2a (*n* = 990 distances in the similar group, 2025 distances in the different group, *p* = 0.62, *Z* = 0.5, Wilcoxon rank-sum test). Right: Similar to the left panel but for all 15 oscillatory sessions, including the example in the left panel (*n* = 15 sessions, *p* = 0.87, *Z* = 0.17, Wilcoxon rank-sum test). A fraction of 10% of the total number of locked cells was used to define the groups with small and large participation indexes. Symbols as in Fig. 3b.

Finally, we quantified the duration of epochs with uninterrupted population oscillation. We calculated the inter-cycle interval (ICI) for all cycles in one session as the amount of time that elapsed between the moment the phase of the oscillation reaches *π* after completing one turn along the ring, and the moment it is equal to −*π* prior to initiating the next turn along the ring. ICIs were then pooled across sessions. ICIs were present at different lengths, ranging from 0 s when cycles are consecutive (69% of ICIs, 279 of 406 ICIs) to a maximum of 452 s (Fig. 2i). The fraction of session time with population oscillation varied within and across animals (Extended data Fig. 5g), yet - when present - the oscillation could progress uninterruptedly for minutes in each of the animals and span up to 23 consecutive cycles (Extended data Fig. 5h).

Taken together, these results suggest that when animals are engaged in a self-paced locomotor task under minimal sensory stimulation and in the absence of rewards, population activity in the superficial layers of the MEC is organized into a minute-scale population oscillation consisting of periodic sequences of neural activity.

### The majority of MEC neurons are locked to the population oscillation

To determine the degree to which calcium activity in individual MEC neurons was tuned to the population oscillation, we computed for each neuron the locking of its calcium activity to the phase of the oscillation. For each cell, the locking degree was calculated as the length of the mean vector obtained from the distribution of oscillation phases at which calcium events occurred, with a range from 0 (uniform distribution of oscillation phases at which calcium events occur) to 1 (all calcium events occur at the same oscillation phase). Significant locking degrees were observed for the vast majority of the recorded cells (Fig. 3a left, calculated on data from the example session in Fig. 2a; 458 significantly locked neurons over 484 total neurons recorded, or 95%). For these cells, the locking degree was higher than the 99th percentile of a null distribution obtained by temporally shuffling the cell’s calcium events. The observed values of locking degree were consistent with another measure of locking that does not make any assumptions about unimodal distribution of the data: the mutual information between the counts of calcium events and the phase of the oscillation (Fig. 3a right, Extended data Fig. 6a, bin size 0.52 s). The predominance of phase-locked neurons was observed in all 15 oscillatory sessions (Fig. 3b, 5841 locked neurons out of 6231, ~94%).

The locking degree was highest for cells with an oscillatory frequency similar to the frequency of the population oscillation, which constituted the large majority of the cells (Extended data Fig. 6b,c). However, neurons with weak phase locking were still engaged in the population oscillation, since the oscillatory sequences were maintained when neurons with high locking degree were excluded (Extended data Fig. 6d). Yet, the more cells were excluded, the harder it was to observe the population oscillation, indicating that the oscillatory dynamics is a property of neural populations (Extended data Fig. 6d). Because the population oscillation involves the vast majority of MEC neurons in the recording region (~95%), the sequences most likely include a mixture of functional cell types such as grid, head-direction, and object-vector cells, given that (i) no cell type accounts for more than 15-25% of the cells in layer II of dorsal MEC^41,42^, and (ii) functional cell types in this area are spatially intermixed within field of views of the microscope that are smaller than the one we used^42^.

Each locked neuron exhibited a preference for activity within a narrow range of phases of the oscillation (Fig. 3c calculated on the example session in Fig. 2a, Extended data Fig. 6e). Across the entire recorded population, all phases of the oscillation were equally present when the cells’ preferred phase was calculated as the mean oscillation phase at which the cell’s calcium events occurred. A uniform distribution was observed both within individual experimental sessions (Fig. 3c) and across all sessions with oscillations (Extended data Fig. 6e,f).

Not all neurons participated in each individual oscillation cycle, however. We quantified the degree to which cells skipped cycles through a participation index (PI), calculated as the fraction of cycles needed to account for 90% of the total amount of calcium events of a neuron in one session. For neurons that were active only in a few cycles the participation index was small (participation index *~* 0), and for neurons that were reliably active during most of the cycles the participation index was high (participation index *~* 1, Extended data Fig. 6g shows three example neurons of the session in Fig. 2a). Participation index variability was observed both within individual experimental sessions (Fig. 3d left), and across all oscillatory sessions (Fig. 3d right). The participation index did not correlate with the degree to which the single cell oscillatory frequency matched the population oscillation frequency (Extended data Fig. 6h).

We next asked how differences between cells’ preferred phase or participation mapped onto the MEC surface (Extended data Fig. 1c), keeping in mind that the population oscillation in MEC could have features of travelling waves, where the population activity moves progressively across anatomical space^43–48^. We calculated pairwise anatomical distances in the microscope’s field of view (i) between cells with similar preferred phases and (ii) between cells with different preferred phases (Extended data Fig. 6i). If topographical organization were present, we would expect smaller pairwise anatomical distances for the cells with the most similar preferred phases. The results showed, however, that cells with similar and dissimilar preferred phases were anatomically intermingled (Movie 1 and Fig. 3e,f). Changing the fraction of cells used to define the groups with similar and different preferred phases had no impact on the degree of intermixing (Extended data Fig. 6j). There was also no topography in the neurons’ participation indexes (Fig. 3g,h; Extended data Fig. 6k,l).

Taken together, these findings suggest that even though the majority of MEC neurons were locked to the population oscillation, both their locking degree and the participation in individual cycles varied across the population.

### The population oscillation consists of periodic and unidirectional activity sequences

We next sought to quantify the sequential activation and periodicity of neural activity during the population oscillation. In order to average out the variability observed at the single cell level in terms of oscillation frequency, locking degree and participation index (Fig. 1,3), we decided to study the neural population dynamics by means of ensembles of co-active cells (Extended data Fig. 7a). As expected, the participation index increased when activity was considered for merged groups of cells, or neuronal ensembles, instead of single neurons (Extended data Fig. 7b). Because the participation index plateaued after 5 merging iterations, consisting of approximately 10 ensembles depending on the session, we chose to assign neurons to a total of 10 ensembles, based on their proximity in the sorting obtained through the PCA method (Fig. 4a, note that each cell belongs to only one ensemble). Ensembles were representative of the activity of their assigned neurons (Extended data Fig. 7c-f), their activity oscillated at the same frequency as the population oscillation (Extended data Fig. 7g,h), and the ensembles were anatomically intermingled (Extended data Fig. 7i-k).

To quantify the temporal dynamics of the ensemble activity, we calculated the probability by which activity transitioned between ensembles across adjacent time bins, and expressed the resulting probabilities as a transition matrix (Fig. 4b). For each time bin of the recording session (bin size shown in Extended data Fig. 7l), we only kept the ensemble with the highest activity (red rectangle in Extended data Fig. 7m; Extended data Fig. 7n). The analysis revealed (i) that transitions between adjacent ensembles were more frequent than transitions between ensembles that were farther apart, and (ii) that transitions occurred with a preferred directionality (Fig 4b, left). Transitions from ensemble 10 to ensemble 1 were equally frequent as transitions between consecutive ensembles (Extended data Fig. 7o), as expected from the periodic nature of the population oscillation. No such structure was seen in transition matrices obtained after shuffling the calcium activity of all cells (Fig 4b, right). The findings were upheld when the transition matrix was used as an adjacency matrix to build a directed weighted graph (Extended data Fig. 7p).

We then asked whether preferences in nearby ensemble transitions gave rise to stereotyped activity sequences. We calculated the probability of sequential ensemble activation by counting the number of times that, within one session, a given number of ensembles was activated in a strictly ascending manner. The procedure was applied on both recorded and shuffled data (Fig 4c). In the oscillatory sessions the sequential activation of three of more ensembles was 2.3 times more likely in the recorded data than in the shuffled data (probability of sequential activation of ≥ 3 ensembles in recorded data = 0.62; probability of sequential activation of ≥ 3 ensembles in shuffled data = 0.27). These findings motivated us to compute a sequence score, which quantifies how sequential the ensemble activity is within a session. The sequence score was calculated as the probability of observing three or more ensembles sequentially activated. As expected, the score was larger for sessions that were classified as oscillatory according to the oscillation score (which quantifies the presence of single cell periodic activity, Extended data Fig. 7q). Statistical significance was assessed by temporally shuffling the matrix of calcium activity. While sequence scores were significant in 100% of the oscillatory sessions (15 of 15), significant sequential activity was demonstrated also in 41% of the non-oscillatory sessions (5 of 12, Extended data Fig. 7r). Taken together, our results suggest that the population oscillation is composed of periodic, sequential and unidirectional activation of ensembles of highly-correlated neurons.

### The population oscillation is present during both running and immobility

Fast oscillations and single cell firing in the entorhinal-hippocampal system can be modulated by a number of movement-associated parameters, such as running state, position, travel distance, running speed and acceleration^2,3,49,50^. These relationships prompted us to investigate whether similar dependencies are present for population oscillations that occur at the seconds-to-minutes time scale (Fig. 5a).

**Figure 5.**
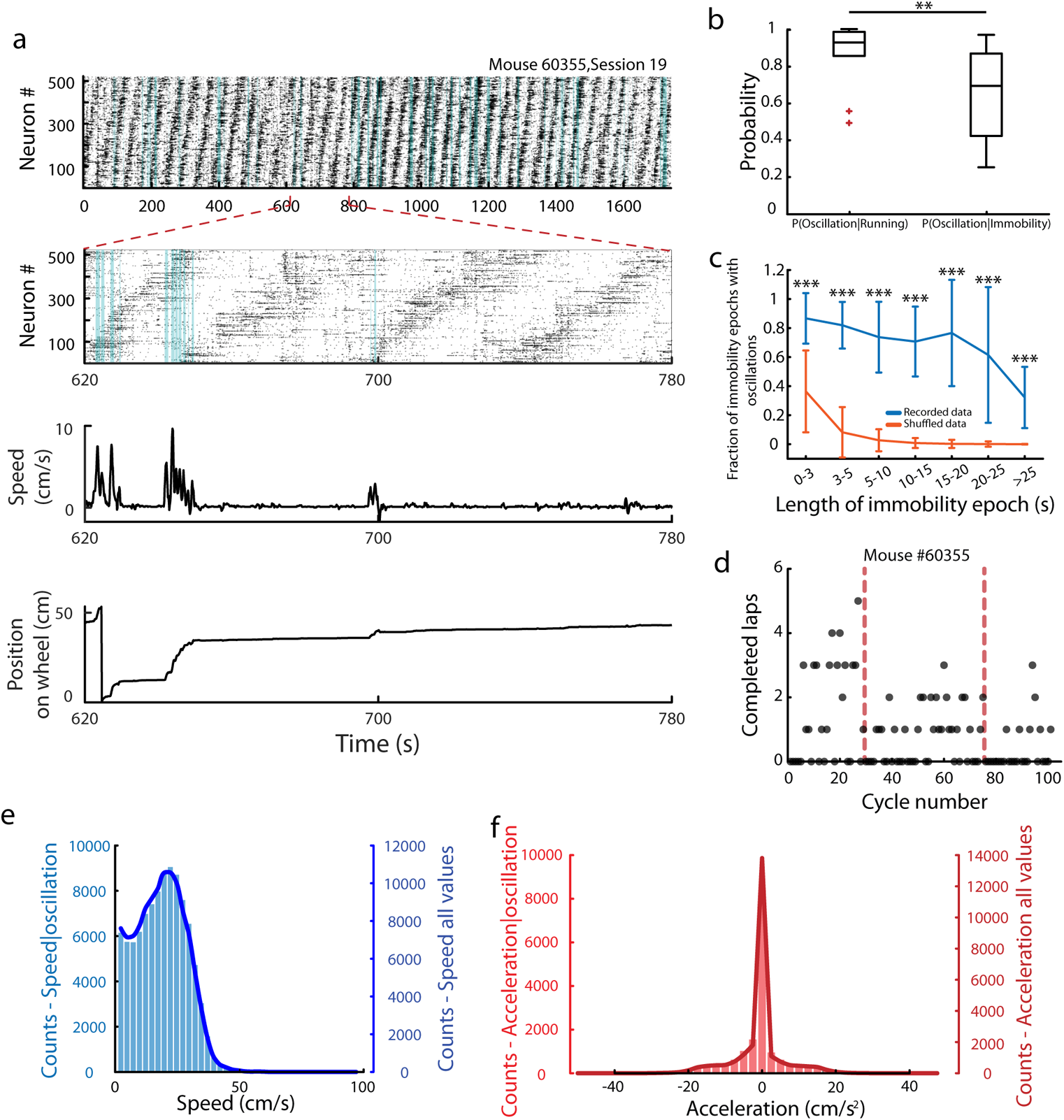
The MEC population oscillation is independent of movement. **a.** Top: raster plot of one recorded session (30 min, 520 neurons). Time bins colored in blue indicate that the animal ran faster than 2 cm/s. Inset indicates 160 s of neural activity. Middle: Instantaneous speed of the animal. Bottom: Position of the animal on the wheel, expressed relative to an arbitrary point on the wheel. **b.** Box plot showing probability of observing the population oscillation given that the animal was either running or immobile (median probability of oscillations during running = 0.93; median probability of oscillations during immobility = 0.69; two sample Wilcoxon signed-rank test on the probability of oscillation for running vs. immobility, *n* = 10 oscillatory sessions over the 3 animals that had the tracking synchronized to imaging, *p* = 0.002, *W* = 55). Box-plot symbols as in Fig. 3b. **c.** Fraction of immobility epochs with population oscillation as a function of length of the immobility epoch (mean ± S.D.). For each length bin, the fraction of immobility epochs with population oscillation was averaged across sessions (n = 10 oscillatory sessions over 3 animals). Note the continued presence of oscillations during extended immobility intervals. Blue: recorded data (n = 10 per length bin); Orange: shuffled data (n = 5000 per length bin, 500 shuffled realizations per session were pooled). Recorded vs shuffled data: *p ≤*2.62× 10^−6^, 4.7*≤ Z ≤* 47.5, Wilcoxon rank-sum test. **d.** Number of completed laps as a function of cycle number. Each dot indicates one individual cycle. Three sessions recorded in one animal are pooled. Dashed line indicates separation between sessions. **e.** Distribution of speed values during the fraction of the session with population oscillation (blue bars; n = 167389 time bins across cycles of 10 oscillatory sessions, bin size = 129 ms) and for the entire session (blue solid line, with and without oscillation; n = 238505 time bins across 10 oscillatory sessions over 3 animals, bin size = 129 ms). Note the almost identical shape of the distributions, suggesting there is no specific range of speed values associated with the population oscillation. **f.** As in (e) but for the distribution of acceleration values. There is no difference in the range of acceleration values during the fraction of the session with population oscillation.

We first sought to determine whether the population oscillation is associated with specific behavioural states such as running (animal moves along the rotating wheel) or immobility (position on the wheel remains unchanged, regardless of body movement). The amount of running vs. immobility varied substantially across sessions (the fraction of time spent running ranged from 0.04 to 0.87, median = 0.53, Extended data Fig. 2a). Analyses of the relationship between movement and population oscillations were restricted to 10 oscillatory sessions in 3 animals, for which the behavioural tracking was successfully synchronized to imaging (Online Methods). By following a two-step approach, we estimated for these sessions the probability of observing the population oscillation given that the animal was either running or immobile. First, for each session we identified the time bins that belonged to individual cycles of the oscillation (Extended data Fig. 5a), and we labeled those bins as “oscillation bins”. The fraction of bins labeled as oscillation bins was 0.73 ± 0.07 (mean ± S.E.M., n = 10 sessions). Next, to compute the conditional probabilities we assigned a second label to each bin depending on whether it occurred during running or immobility (speed *≥* or < 2cm/s; fraction of bins labeled as running = 0.43 ± 0.09, mean ± SEM, n = 10 sessions). We found that the population oscillation was predominant during running bouts (Fig. 5b, left), but it was also observed during immobility (Fig. 5b, right), suggesting that the oscillation is not exclusive to epochs in which the animal is engaged in locomotion. Population oscillation cycles were continuous for immobility durations spanning from 1 s to more than 25 s (Fig. 5c, Extended data Fig. 2b). The continued presence of the population oscillation during long epochs of immobility suggests that behavioural state and running distance have a limited role in driving the progression of the population oscillation in MEC, and stands in contrast to previous observations in CA1 of the hippocampus, where stereotypic sequences of neural activity subsided within 2 seconds after motion was terminated^36^.

We then asked whether the population oscillation is modulated by the animal’s position, running speed or acceleration on the wheel, which all varied substantially during the course of a session. Across the 10 sessions, the range of speed values was 0-75.4 cm/s, with a median of 0 cm/s and a median during running behaviour of 7.8 cm/s, whereas acceleration values ranged from −86.3 to 108.9 cm/s^2^, with a median of 0 cm/s^2^ for all the data as well as the running epochs specifically. The number of laps the animals completed on the wheel during one oscillation cycle was also highly heterogeneous, ranging from 0 to 86 laps per cycle across all animals (lap length~54 cm; Fig. 5d, Extended data Fig. 2c), suggesting that the progression of oscillatory activity did not map the animal’s position on the wheel. To determine the impact of running speed on the population oscillation, we extracted all oscillation bins and calculated the distribution of observed speed values during those bins. This distribution was almost identical to the distribution of speed values observed during the full length of the sessions, which also included epochs without the population oscillation (Fig. 5e, 314 cycles pooled across 10 oscillatory sessions). Oscillation cycles took place during a wide range of speed values, spanning from 0 to more than 50 cm/s (Fig. 5e). Oscillation bins were similarly independent of acceleration (Fig. 5f).

Finally, since the population oscillation was observed more often during running bouts (Fig. 5b), we investigated whether changes in speed were associated with the initiation of oscillation cycles. We found no difference in speed 10 s before and after cycle onset (Extended data Fig. 2d, left; see Extended data Fig. 2e-h for individual recordings). This result also held for cycles that were 10 s or more apart, i.e. for cycles that belonged to different epochs with uninterrupted population oscillation (Extended data Fig. 2d, right). Altogether, our results show that the MEC population oscillation is not modulated by the animal’s position on the wheel and is only mildly modulated by the animal’s running behavior, consistent with the idea that the entorhinal network can generate such oscillations using intrinsic mechanisms.

### Population oscillations were not observed in other brain regions

The fact that ultraslow oscillations have been reported in widely different brain areas^21–29^ prompted us to investigate whether a population oscillation composed of oscillatory sequences of neural activity of the kind we found in the MEC could be observed in other regions too. We recorded the activity of hundreds of cells in the superficial layers of the parasubiculum (PaS), a high-end parahippocampal region abundant with grid, head-direction and border cells but with a different circuit structure and a weaker theta rhythmicity than MEC^42,51^ (25 sessions over 4 animals, Extended data Fig. 8a,b). In addition, we investigated the superficial layers of visual cortex^52^ (VIS, 19 sessions over 3 animals, Extended data Fig. 8c), which differs from MEC^53,54^ in its network architecture and in the high dimensionality of its neural population activity. The mice performed the same minimalistic self-paced running task as in the MEC recordings (range of speed values in PaS/VIS animals across sessions=0-58.6/0-60.3 cm/s; median number of completed laps on rotating wheel in PaS/VIS animals across sessions=145/104; maximum number of completed laps on rotating wheel in PaS/VIS animals across sessions=502/1743).

To determine whether single-cell calcium activity in PaS and VIS was periodic and oscillated at ultraslow frequencies we followed the same procedure as for the MEC sessions. We found that the calcium activity of a fraction of cells in both brain areas was indeed periodic (Fig. 6a,b). However, in neither brain region were these oscillations organized into sequences of neural activity. The population oscillation was observed neither with the PCA method nor with pairwise correlations or any of the non-linear dimensionality reduction techniques that we had used (Fig. 6c-f; Extended data Fig. 9a). When projected onto a linear low-dimensional embedding, the population activity did not display a ring-shaped topology (Extended data Fig. 9b,c). For all sessions in PaS and VIS, the oscillation scores were lower than the threshold defined from the MEC data to classify sessions as oscillatory (Extended data Fig. 9d; threshold = 0.72, see Extended data Fig. 4c), suggesting that a population oscillation was weak or absent (Fig. 6g). Taken together, these results suggest that MEC has network mechanisms for sequential coordination of single-cell oscillations that are not present in PaS or VIS.

**Figure 6.**
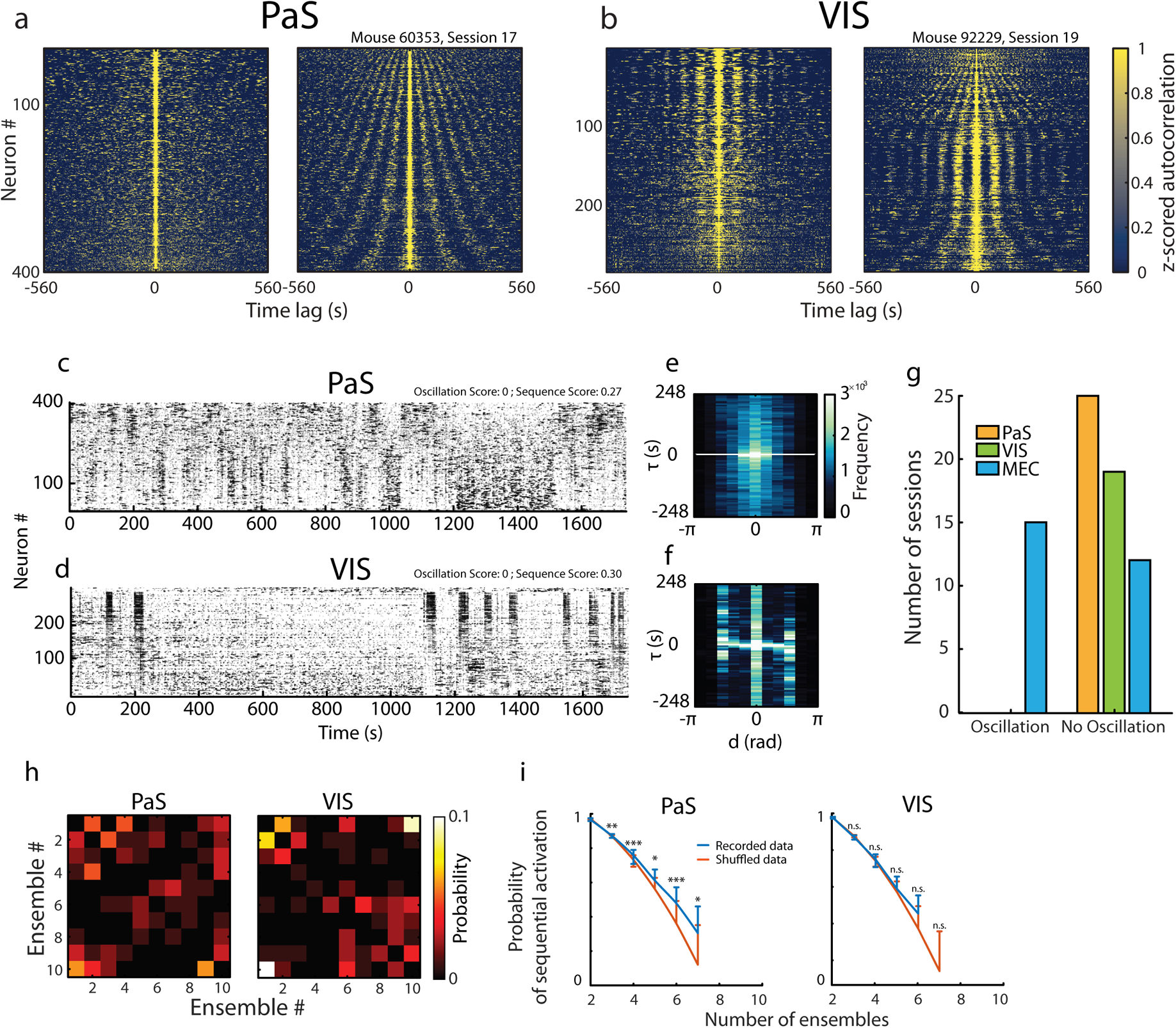
The population oscillation is not observed in parasubiculum or visual cortex. **a,b.** Stacked autocorrelations for two example sessions recorded in parasubiculum (a, PaS; 1800 s, 402 simultaneously recorded neurons) and visual cortex (b, VIS; 1800 s. 289 simultaneously recorded neurons). Each row is the autocorrelation of one cell’s calcium activity, plotted as a function of time lag. Z-scored autocorrelations are color-coded. Cells are sorted according to maximum power (left of each panel) or peak frequency (right of each panel) of the PSD, as in Fig. 1b. **c,d.** PCA-sorted raster plots (as in Fig. 2b) for two example sessions recorded in PaS (Fig. 6a) and VIS (Fig. 6b). Notice lack of stereotyped sequences of activity. Oscillation score and sequence score are indicated at the top. **e,f.** Joint distributions of time lag *τ* that maximizes the cross-correlation between any given pair of neurons and their distance *d* in the PCA sorting (as in Fig. 2f), applied to the recordings in Fig. 6a (PaS) and 6b (VIS). Normalized frequency is color-coded. Notice lack of linear relationship between *d* and *τ*, in contrast to Fig. 2f. **g.** Number of sessions with and without population oscillation in MEC (blue, 27 sessions in total), VIS (green, 19 sessions) and PaS (yellow, 25 sessions) based on oscillation scores and threshold defined from the MEC dataset (see Extended data Fig. 4c). **h.** Transition probabilities between ensembles across consecutive time bins (bin size ~ 8.5 s) for the PaS example session in Fig. 6a (left) and the VIS example session in Fig. 6b (right). Symbols as in Fig. 4b. **i.** Probability of sequential ensemble activation as a function of the number of ensembles that are sequentially activated in PaS (left) and VIS (right) (mean ± S.D.). Blue, recorded data (25 PaS sessions; 19 VIS sessions); orange, shuffled data. For each session, the probability of sequential ensemble activation was calculated over 500 shuffled realizations, and shuffled realizations were pooled across sessions for each brain area separately. Probability is shown on a log-scale. In PaS the probability of long sequences was significantly larger in experimental data than in shuffled data (For 3-7 ensembles: *n* = 25 PaS sessions, 12500 shuffled realizations, range of *p* values: *5*.7 × 10^−4^ to 0.036, range of *Z* values: 1.80 to 3.25, one-tailed Wilcoxon rank-sum test). This was not the case in VIS (For 3-6 ensembles: *n* = 19 VIS sessions, 9500 shuffled realizations, range of *p* values: 0.09 to 0.99, range of *Z* values: −3.34 to 1.36, one-tailed Wilcoxon rank-sum test).

While ensemble analyses showed no preference for transitions between co-active cells that were nearby in the PCA sorting (Fig 6h; Extended data Fig. 10a-d), the probability of sequential activation in PaS was significantly larger in experimental data than in shuffled data (Fig. 6i left, n = 25 sessions). No significant difference was found in VIS (Fig. 6i right, n = 19 sessions). In line with this result, the percentage of sessions with significant sequence scores (defined as the probability of observing the sequential activation of 3 or more ensembles) was highest for oscillatory sessions in MEC (15 out of 15, 100%), intermediate for PaS (7 out of 25, 28%) and lowest for VIS (1 out of 19, 0.05%) (Extended data Fig. 10e). Features of the animal’s behaviour were not different between sessions with significant and non-significant sequence scores (Extended data Fig. 10f-i; all regions).

Finally, the presence of sequential ensemble activation in many PaS recordings, but not in VIS, motivated us to investigate whether this difference could reflect a stronger tendency for VIS neurons to cluster into a few synchronized states. We quantified synchronization through the absolute value of the correlation between the calcium activity of all cell pairs, as well as co-activity, calculated as the fraction of cells that had simultaneous calcium events in bins of 129 ms. Data from VIS had both higher correlation values (Extended data Fig. 10j) and higher co-activity (Extended data Fig. 10k), compared to PaS. The strong synchronization of calcium activity in VIS is consistent with previous observations of recurring co-activity among subsets of neurons (ensembles) in this brain region^37,55^.

## Discussion

Our experiments identify an ultraslow neural population oscillation that organizes neural activity in the majority of neurons recorded in the MEC of awake head-fixed mice during self-paced running as well as intermittent segments of rest. Across recording sessions, the length of individual oscillation cycles can range from tens of seconds to minutes, but the time scale is generally fixed within an individual recording session. This oscillation is expressed as unidirectional sequences of activity that can repeat uninterruptedly for tens of minutes. The oscillation entrains periodic activity in individual neurons despite some variability in the frequency of single cell activity. Individual cells are activated at specific phases of the population oscillation, with phase preferences distributed uniformly across the population. Unlike faster oscillations, which typically have the greater part of the neural activity centered within a narrow segment of the oscillation cycle^56–58^, the population oscillation in MEC maintains a relatively constant activity rate throughout the cycle, reflecting the steady progression of a sequence of neural activity.

Oscillations at time scales of tens of seconds to minutes have been reported in individual cells of multiple brain areas and in a variety of brain states including anaesthesia, sleep and alert immobility^21–26^. EEG and fMRI recordings from awake humans^59–62^, as well as LFP from anaesthetized and awake animals^27–29^ have similarly demonstrated oscillatory changes at periods of 10 s or longer. However, we do not know from those observations how the activity of individual neurons is organized with respect to each other during the oscillation. The present data demonstrate an ultraslow population oscillation at a time scale ranging from tens of seconds to minutes that, in a controlled behavioral setting and under minimal sensory stimulation, engages sequentially the vast majority of neurons in the recorded area of MEC. The population oscillation is only mildly modulated by the animal’s running behavior and is therefore more likely to emerge from intrinsic network mechanisms.

The ultraslow oscillatory sequences of the MEC stand out from instances of slow sequential neural activity that have not been described in terms of oscillations. In the hippocampus, neural activity in CA1 cells is organized into stereotypic sequences when rats or mice run on a rotating wheel in a cue-rich environment^63^ or in a sensory-restricted environment similar to the one used here^36,64^. Unlike what we observed in MEC, these sequences are more strictly coupled to ongoing behavioral activity and running distance^36^, and they have not been found to exhibit temporal periodic organization at a well-defined frequency. Moreover, while nearly 95% of all MEC neurons in the present study were locked to the population oscillation and most of them participated in at least half of the cycles, hippocampal sequences involve only a small fraction of the network (5% in ref. 36). Such a difference in participation would be in agreement with the view that the MEC supports a low-dimensional population code where the cells’ responses covary across environments and behavioural states^53,54,65^, whereas the hippocampus supports a more high-dimensional population code that may orthogonalize distinct experiences^66–69^. The MEC population oscillation also differs from retinal waves and cortical waves in the developing nervous system^43–48^, as well as travelling waves in the adult hippocampus^70,71^, which all move progressively through anatomical space, in a topographic manner not observed in the present data.

The presence of oscillatory activity in individual cells of all three regions – visual cortex, parasubiculum and MEC – together with the absence of population oscillation in the former two, points to MEC as having network mechanisms for sequential coordination of single cell oscillations that are not present in parasubisulum or visual cortex. Such mechanisms might share similarities with prewired sequences in the hippocampus^72^, or they may be supported by plasticity rules operating on slow time scales^73^. The population oscillation of the MEC is consistent with dynamics expected in a one-dimensional continuous attractor network^74–76^ where cells are conceptualized as lying on a functional ring with positions determined by the cells’ loadings on the first two principal components. However, it has not been determined whether such ring-like connectivity exists among the high percentage of MEC neurons entrained by the population oscillation, and, if there is such connectivity, which signal is responsible for moving the bump of activity along the ring. Sequential activity could also be generated by other types of structured connectivity, for example in recurrently connected networks^77,78^ and in feedforward networks in which sequences may arise through synfire chains or rate propagation^79–83^. But while structured connectivity might allow for slow transitions in the ensemble activity^84,85^, none of the mechanisms proposed so far generate minute-scale repeating sequences with the variability at the single-cell level that we here observed in the MEC population oscillation.

We propose two related functions for the MEC population oscillation. First, the reported oscillations might have a role in large-scale coordination of entorhinal circuit elements^7,10,86,87^, either by synchronizing faster oscillatory activity^25,29, 60–62^, such as theta and gamma^11,12,32^, or by organizing neural activity across functionally dissociable cell classes, such as grid cells, head direction cells, border cells, and object-vector cells^2–6,88^. Given that each of these functional cell classes accounts for less than 15-25% of the local cell population in dorsal MEC^41,42^, whereas nearly all recorded neurons in the present study were locked to and participated in the population oscillation, the population oscillation is likely to consist of a mixture of functional cell types. Coordination by the population oscillation may prevent functional cell classes from drifting apart and help the circuit maintain correlated firing over the entire duration of an experience^89,90^. Second, the MEC oscillatory sequences may act as a scaffold to support computations that must take place on the fly with little time for extensive circuit plasticity, such as fast storage of memories associated with one-time experiences^69,91^, or the assembly of representations for complex sensory stimuli that evolve over time^37,92^. The ultraslow population oscillation may also enable the circuit to keep track of time during extended behavioral experiences^93^. The temporally organized firing of time cells in MEC^94,95^ and the more extended temporal evolution of neural population activity in LEC^96^ may be facilitated by an underlying sequence template. Whether the population oscillations serve such coordination and scaffolding functions across a broader spectrum of behaviors than in the minimalistic task used here remains to be determined. If similarly slow periodic sequences are expressed across a wider span of behaviors, including sleep and free exploration, they must interface with dynamics of MEC cells on a number of manifolds, such as in ensembles of head direction cells and grid cells^54,97,98^.

## Acknowledgments

We thank W. Zong, C. Battistin, I. Davidovich, Y. Roudi and E. Kropff for discussion, Y. Burak for discussion and comments on the manuscript, D.W. Tank for sharing hardware, software and advice during the earliest stages of the study, and A. Tsao, G.B. Keller and T. Bonhoeffer for help in setting up initial procedures for benchtop 2-photon imaging for mice running on a rotating wheel. We thank A.M. Amundsgård, K. Haugen, K. Jenssen, E. Kråkvik, I. Ulsaker-Janke, and H. Waade for technical assistance. The work was supported by a Synergy Grant to E.I.M. and Yoram Burak from the European Research Council (‘KILONEURONS’, Grant Agreement N° 951319), an RCN FRIPRO grant to E.I.M. (grant number 286225), a Centre of Excellence scheme grant to M.-B.M. and E.I.M. and a National Infrastructure grant to E.I.M. and M.-B.M. from the Research Council of Norway (Centre of Neural Computation, grant number 223262; NORBRAIN, grant number 295721), the Kavli Foundation (M.-B.M. and E.I.M.), a direct contribution to M.-B.M. and E.I.M. from the Ministry of Education and Research of Norway, and a European Research Council Starting Grant (ERC-ST2019 850769) and an Eccellenza Grant from the Swiss National Science Foundation (PCEGP3_194220) to F.D.

## Author Contributions

F.D., H.O., M.-B.M. and E.I.M. planned and designed the initial experiments, with later input from S.C.G.; F.D. and R.I.J. performed the experiments; H.O. developed hardware and imaging software, preprocessed the data and performed initial analysis; S.G.C., M.-B.M. and E.I.M. conceptualized and proposed analyses, with input from F.D.; S.G.C. performed analyses of neural activity; F.D. performed histological analyses; S.G.C., F.D., M.-B.M. and E.I.M. interpreted data; S.G.C. and F.D. visualized data; S.G.C. and E.I.M. wrote the paper, with initial contributions from F.D. and with periodic input from all authors; M.-B.M. and E.I.M. supervised and funded the project.

## Supplementary Information

is available for this paper.

## Author Contact Information

Correspondence should be addressed to S.G.C., F.D., M.-B.M. or E.I.M. Requests for materials should be directed to E.I.M.

## Reprints and Permissions

information is available at www.nature.com/reprints

## Legends

**Extended data Figure 1.**
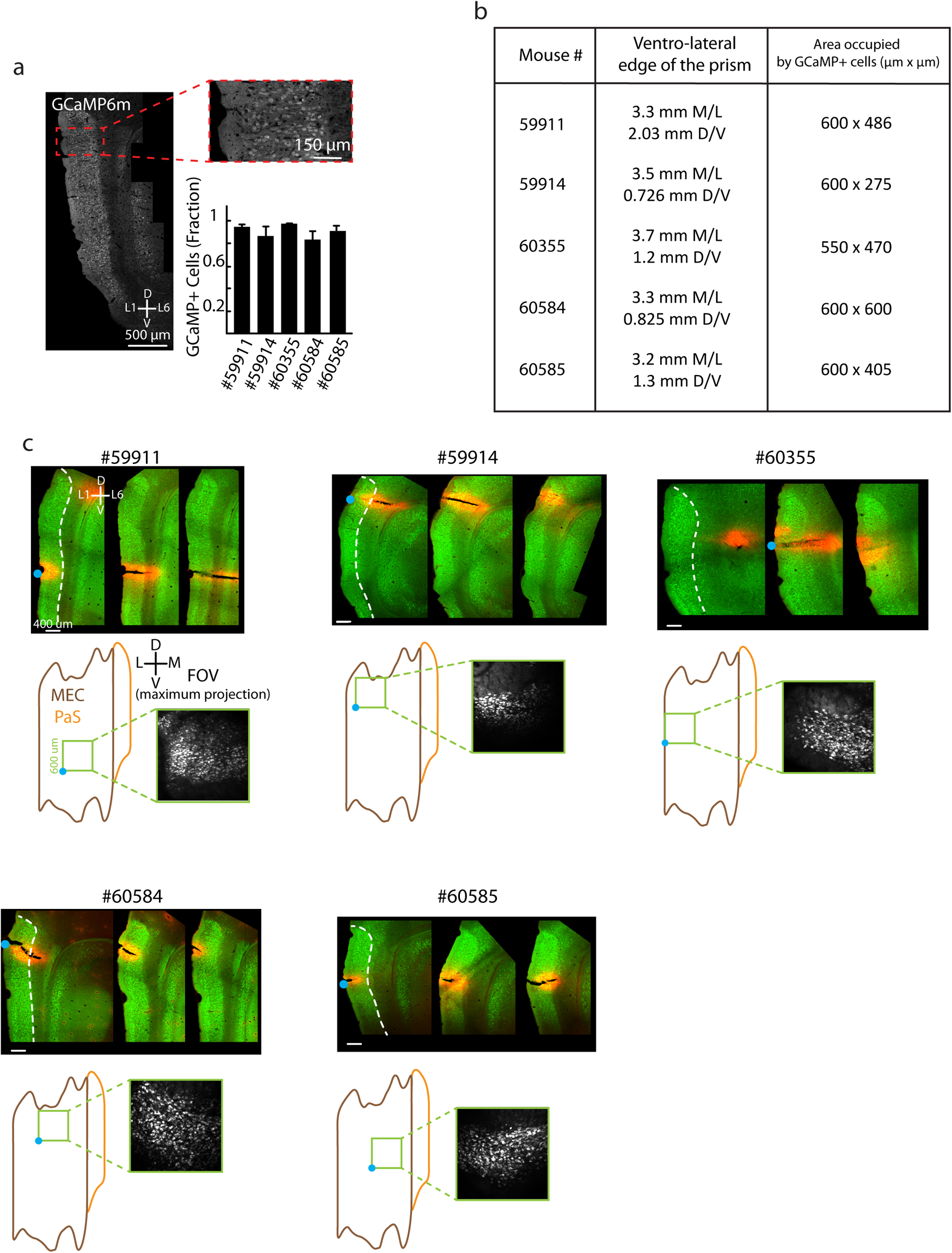
Histology showing imaging locations for each animal in the MEC group. **a.** Left: Representative sagittal image indicating GCaMP6m expression in the superficial layers of the MEC upon local viral injection at postnatal day P1 (sagittal section). Images were acquired with a 20× objective mounted on a confocal laser scanning microscope LSM 880 (Zeiss). Scale bar 500 μm. Red inset and top right: 60× magnification of the most dorsal portion of the MEC. Scale bar 150 μm. Bottom right: Fraction of neurons in the image that express GCaMP6m; data are shown for all 5 animals with MEC imaging. Error bar indicates the S.D. calculated across multiple adjacent slices. **b.** Location of the ventro-lateral edge of the prism in stereotactic coordinates, and area of the FOV occupied by cells expressing GCaMP6m. Data are shown for each MEC-imaged animal. Mouse #59911 had no oscillations. **c.** Prism location in mice that underwent calcium imaging in MEC. Top: Maximum intensity projections of 50 μm thick sagittal brain sections. For each of the 5 mice in (b), 3 sections, shown from lateral (left) to medial (right), were acquired with an LSM 880, 20×. A DiL-coated piano wire pin was inserted at the ventrolateral corner of FOV to enable identification of the FOV on histology sections. Green is GCaMP6m signal, red is DiL signal. Scale bar is 400 μm. The white stippled line encapsulates the superficial layers of MEC. The blue dot adjacent to the leftmost image of the series marks the location of the ventro-lateral corner of the prism. Bottom: estimated location of the FOV for two-photon imaging, projected onto a flat map encompassing MEC (brown outline) and parasubiculum (PaS, yellow outline). The blue dot marks the location of the pin used to demarcate the most lateral-ventral border of the prism, while the green square inset is the microscope’s FOV. Inset images show the mean (left) and maximum (right) intensity projections of the FOV. Anteroposterior (AP) and dorsoventral (DV) axes are indicated in panels a and c.

**Extended data Figure 2.**
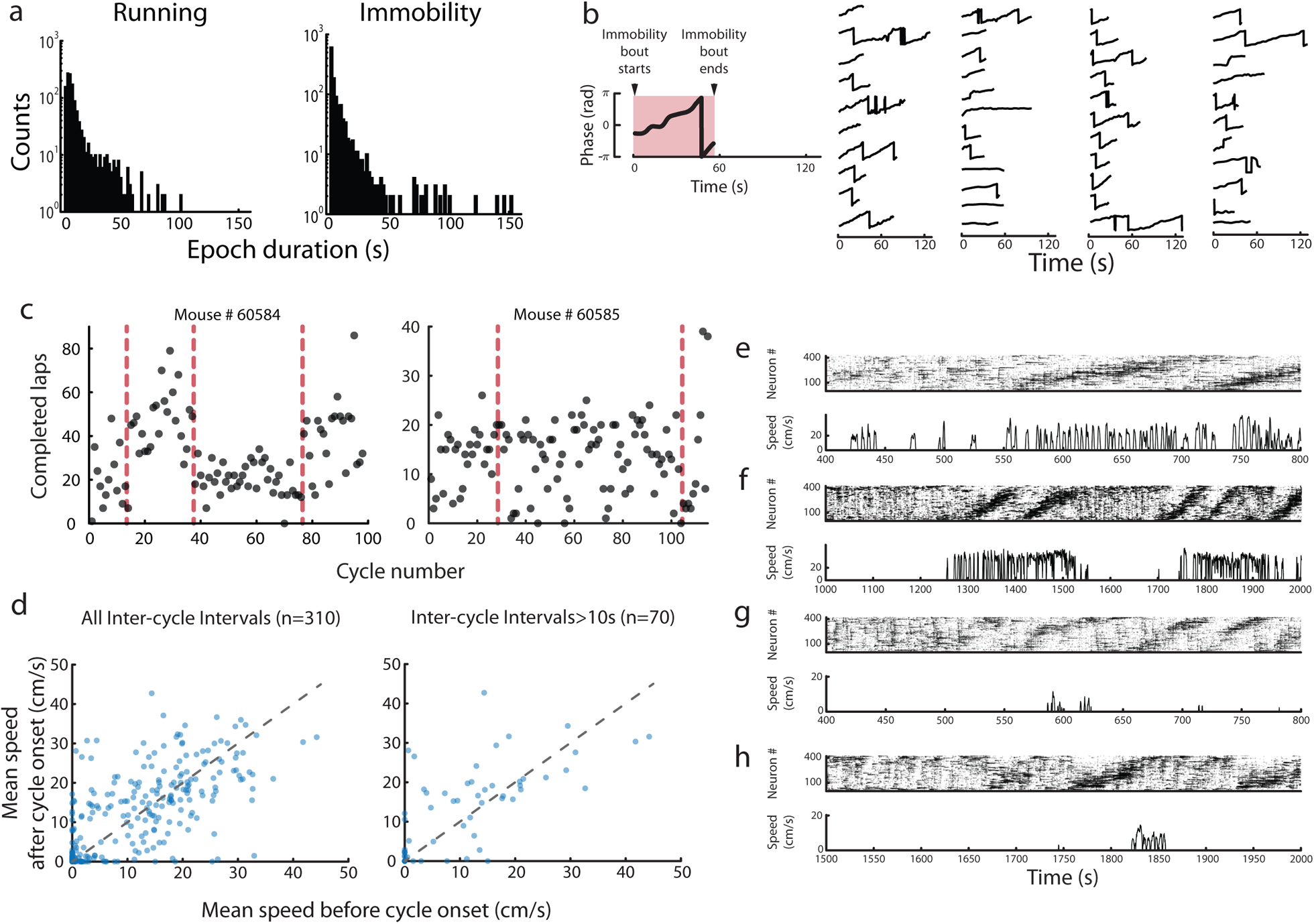
Relationship between the population oscillation and behavior. **a.** Quantification of the animals’ behavior during head-fixation on the wheel. Distribution of duration of running (speed ≥ 2 cm/s, left) and immobility (speed < 2 cm/s, right) epochs for 10 oscillatory sessions over the 3 animals with synchronized behavioral tracking and imaging (1289 running bouts and 1286 immobility bouts in total). Each count is an epoch. **b.** Left: Schematic of the change in phase of the oscillation during immobility epochs that were longer than 25 s and that occurred during the population oscillation. Right: 44 of these epochs from the same 3 mice as in (a). As in the schematic on the left, each line represents the progression of the phase of the oscillation (*y* axis, from –π to π rad) as a function of time (*x* axis, in seconds). The start of each immobility epoch is aligned at t=0, and the epoch lasts for as long as the line continues. Different epochs have different lengths, covering a range from 25 s to 258 s. For visualization purposes only the first 120 s are displayed (3 of the epochs were truncated; these had durations of 127.9, 258.2, 136.1 s). Sudden transitions from π to –π rad reflect the periodic nature of the oscillation. **c.** Number of completed laps on the wheel per cycle of the population oscillation as a function of the cycle number after pooling sessions (range of completed laps on rotating wheel across 10 sessions = 10-1164, median = 624). Sessions are pooled for each animal separately (mouse #60584, 4 sessions; mouse #60585, 3 sessions; the third animal is shown in Fig. 5d). Each dot indicates one individual cycle. The dashed line indicates separation between sessions. **d.** Left: To determine whether the population oscillation is modulated by onset of running we calculated the mean running speed during time intervals of 10 s right before and right after the cycle onset (one sample Wilcoxon signed-rank test on the difference between speed before and after cycle onset, *n* = 310 cycle onsets over 10 sessions from 3 animals, *p* = 0.82, *W* = 25). Right: Same as left but only for cycles that were 10 s or more apart, i.e. for cycles belonging to different oscillatory epochs (one sample Wilcoxon signed-rank test on the difference between speed before and after cycle onset, *n* = 70 cycle onsets over 10 sessions from 3 animals, *p* = 0.12, *W* = 857). Note that there is no systematic change in speed after onset of cycles. **e-h.** Examples of fractions of sessions with increased speed after cycle onset (exceptions from the general pattern shown in d). Top of each panel: Raster plots, symbols as in Fig. 2a (bin size = 129 ms). Bottom of each panel: Instantaneous speed of the animal during the recording in the top panel. Length of the displayed fraction of the session was 400, 1000, 400 and 500 s, respectively, for (e-h). Notice that while speed is higher after onset of the cycle in these examples, the increase of speed does not always occur right after cycle onset, but sometimes before (e,f), and sometimes tens of seconds after (g,h).

**Extended data Figure 3.**
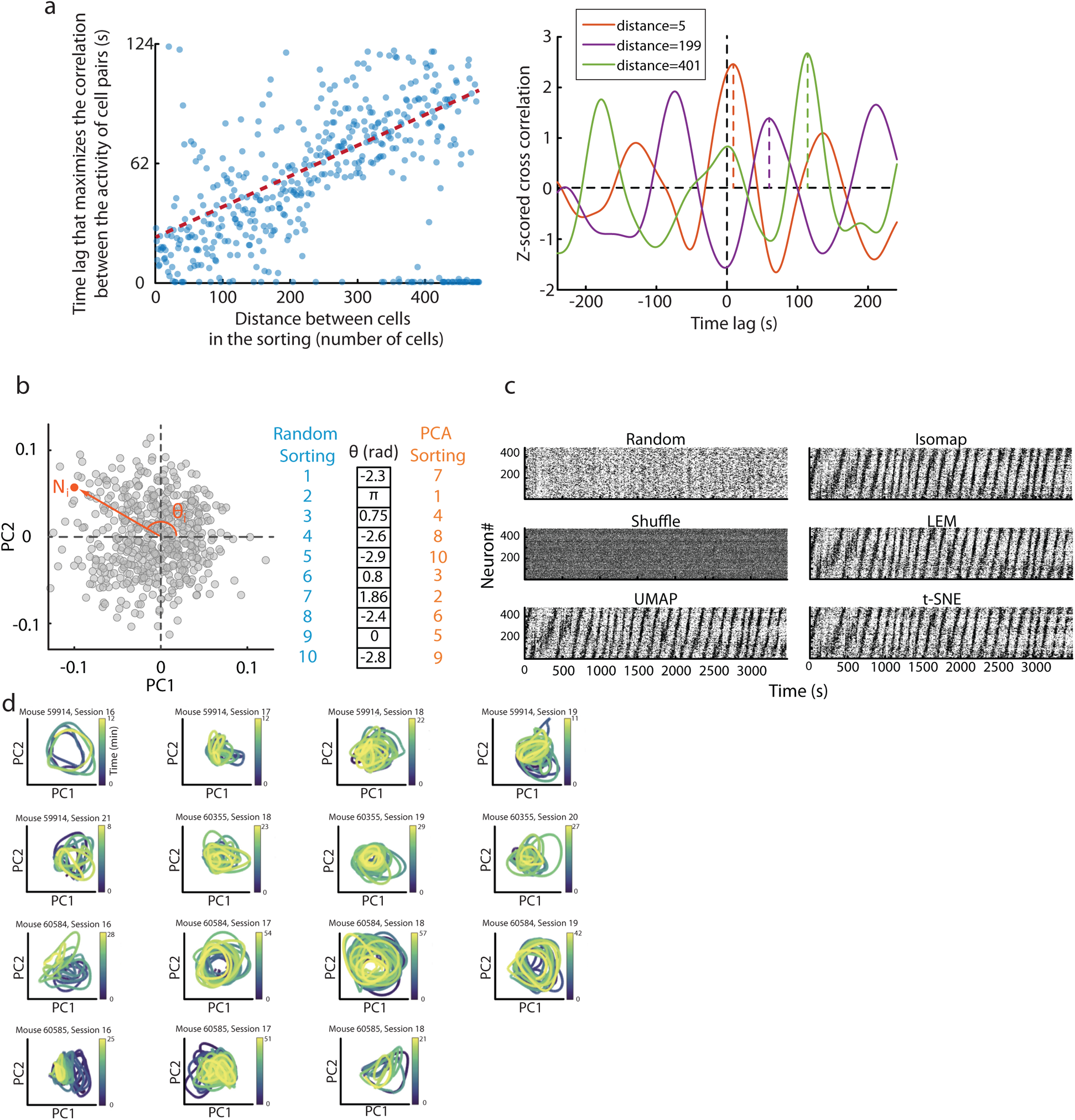
Oscillatory sequences shown by cell sorting based on correlation or dimensionality reduction. **a.** Left: Because neural activity progresses sequentially, the time lag that maximizes the correlation between the calcium activity of pairs of cells increases with their distance in the correlation sorting. Sorting is performed as in Fig. 2a. Time lag is expressed in seconds, distance is expressed as the number of cells between the two cells in the sorting. Notice that for large distances (e.g. > 300 cells), the time lag to peak correlation is either larger than 60 s or close to zero. This bimodality is due to the periodicity of the MEC population oscillation. The dashed line indicates a linear regression (*n* = 301 cell pairs, *R^2^* = 0.17, *p* = 2 × 10^−14^, the line was fitted between the intermediate samples to avoid the effect of the periodic boundary conditions). Right: The cross correlation between the calcium activity of pairs of cells is oscillatory and temporally shifted. Examples are shown for 3 cell pairs with different distances in the sorting based on correlation values. Orange: cells are 5 cells apart; purple: cells are 199 cells apart; green: cells are 401 cells apart. The dotted line indicates the time lag at which the cross correlation peaks within the first peak. Note that the larger the distance between the cells in the sorting, the larger the time lag that maximizes the cross correlation. **b.** Schematic representation of the “PCA method”. Principal component analysis (PCA) was applied to the binarized matrix of deconvolved calcium activity (“matrix of calcium activity”) of individual sessions by considering every neuron as a variable, and every population vector as an observation. The first two principal components (PC1, PC2) were identified. In the plane defined by PC1 and PC2 (left), the loading of each neuron defines a vector, which has an associated angle θ ∈ [*−π*, *π*) with respect to the axis of PC1 (in the schematic, neuron N_i_ (orange) is characterized by an angle θ_i_). Neurons were sorted according to their angles θ in a descending order (right). Cyan: neuron sorting before application of the PCA method. Orange: neuron sorting after the application of the PCA method. **c.** Population oscillations consisting of oscillatory sequences are not revealed with a random sorting of the cells (top left) or when the PCA sorting method is applied to temporally shuffled data (middle left). A population oscillation similar to that of Fig. 2a,b (with correlation sorting or PCA method) is recovered when neurons are sorted according to non-linear dimensionality reduction techniques (UMAP, Isomap, LEM, t-SNE). Each row of each raster plot is a neuron, whose calcium activity is plotted as a function of time (as in Fig. 2a). Every black dot represents a time bin where a neuron was active (bin size = 129 ms). **d.** Projection of neural activity during the population oscillation onto a low-dimensional embedding generated by the first two principal components obtained by applying PCA to the matrix of calcium activity of each session. Each plot shows one session; all 15 oscillatory sessions are presented. Time is color-coded and shown in minutes, and the temporal range corresponds to all concatenated epochs with population oscillation in the session. Neural trajectories are circular, with population activity propagating along a ring-shaped manifold.

**Extended data Figure 4.**
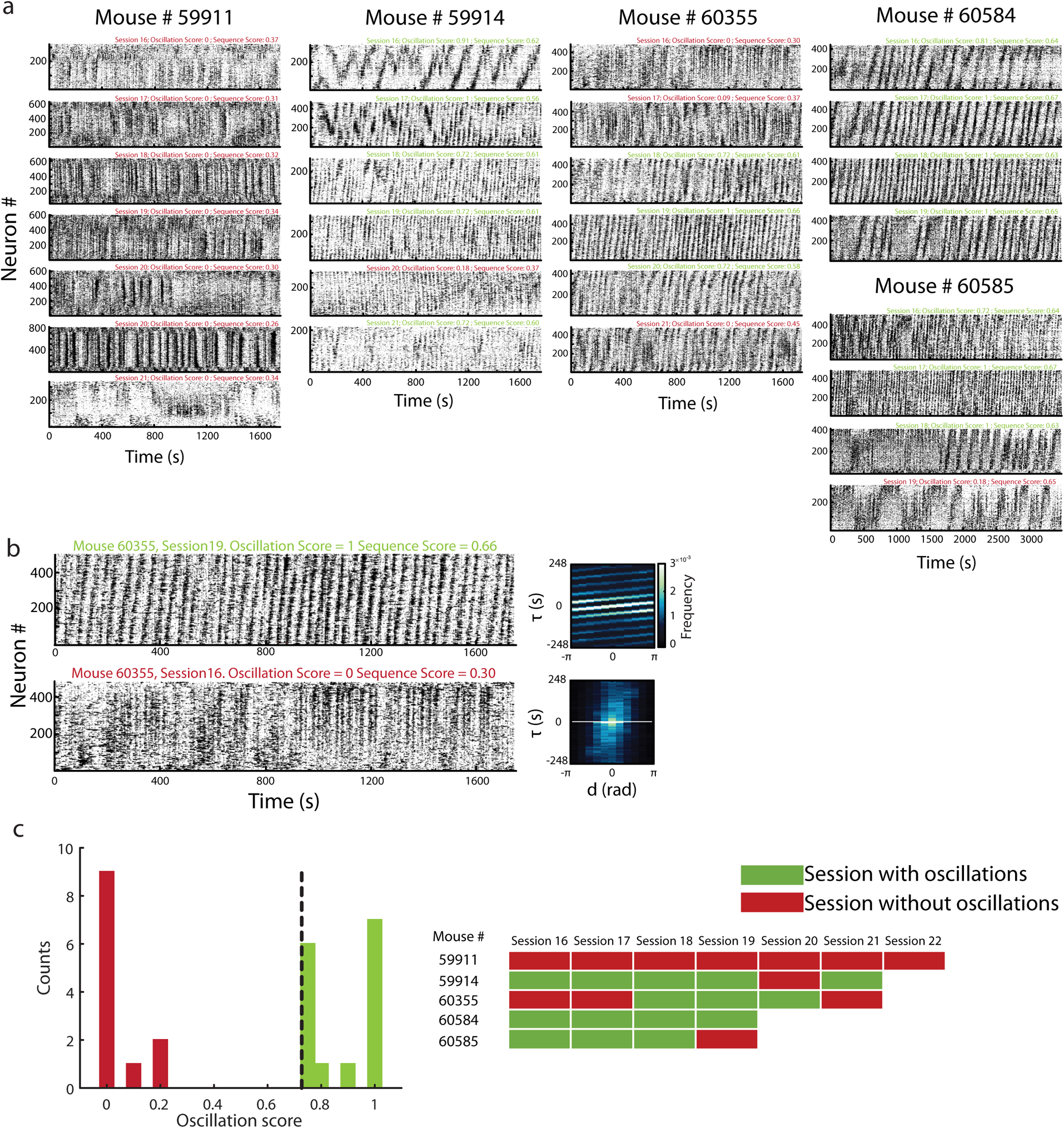
Sorted raster plots for the complete MEC dataset. **a:** PCA-sorted raster plots (as in Fig. 2b) for all analysed sessions across the 5 animals in which MEC population activity was recorded, sorted by animals and day of recording. Session numbering starts the first day of habituation on the wheel, with 15 habituation sessions. One session was recorded per day, and recordings were conducted on consecutive days. Note that sessions had lengths of approximately 1800 s or 3600 s. Oscillation score and sequence score were calculated for each session separately and are indicated at the top right corner of every calcium matrix. The scores colored in green correspond to sessions with population oscillation (see panel c), scores colored in red to sessions without population oscillation. **b:** Example sessions with (top) and without (bottom) population oscillation. These sessions were recorded in the same area of the MEC in the same animal, but on different days (Mouse #60355 in panel a). Left: Raster plots of the matrices of calcium activity. Right: Joint distributions of the time lag *τ* that maximizes the correlation between the calcium activity of any given pair of neurons and their distance *d* in the PCA sorting (as in Fig. 2f). Color code: normalized frequency, each count is a cell pair. Notice the lack of linear pattern in the session without population oscillation. **c.** Left: Distribution of oscillation scores for sessions recorded in MEC (27 sessions in total over 5 animals). Each count is a session. The oscillation score quantifies the extent to which single cell calcium activity is periodic, and ranges from 0 (no oscillations) to 1 (oscillations). Dashed line: Threshold used for classifying sessions as oscillatory (oscillation score ≥ 0.72) or non-oscillatory sessions (oscillation score < 0.72). The threshold was chosen based on the bimodal nature of the distribution. Right: List of sessions sorted by animal and number of sessions the animals experienced on the wheel. Session numbering as in panel a. Red, sessions classified as not oscillatory; green, session classified as oscillatory.

**Extended data Figure 5.**
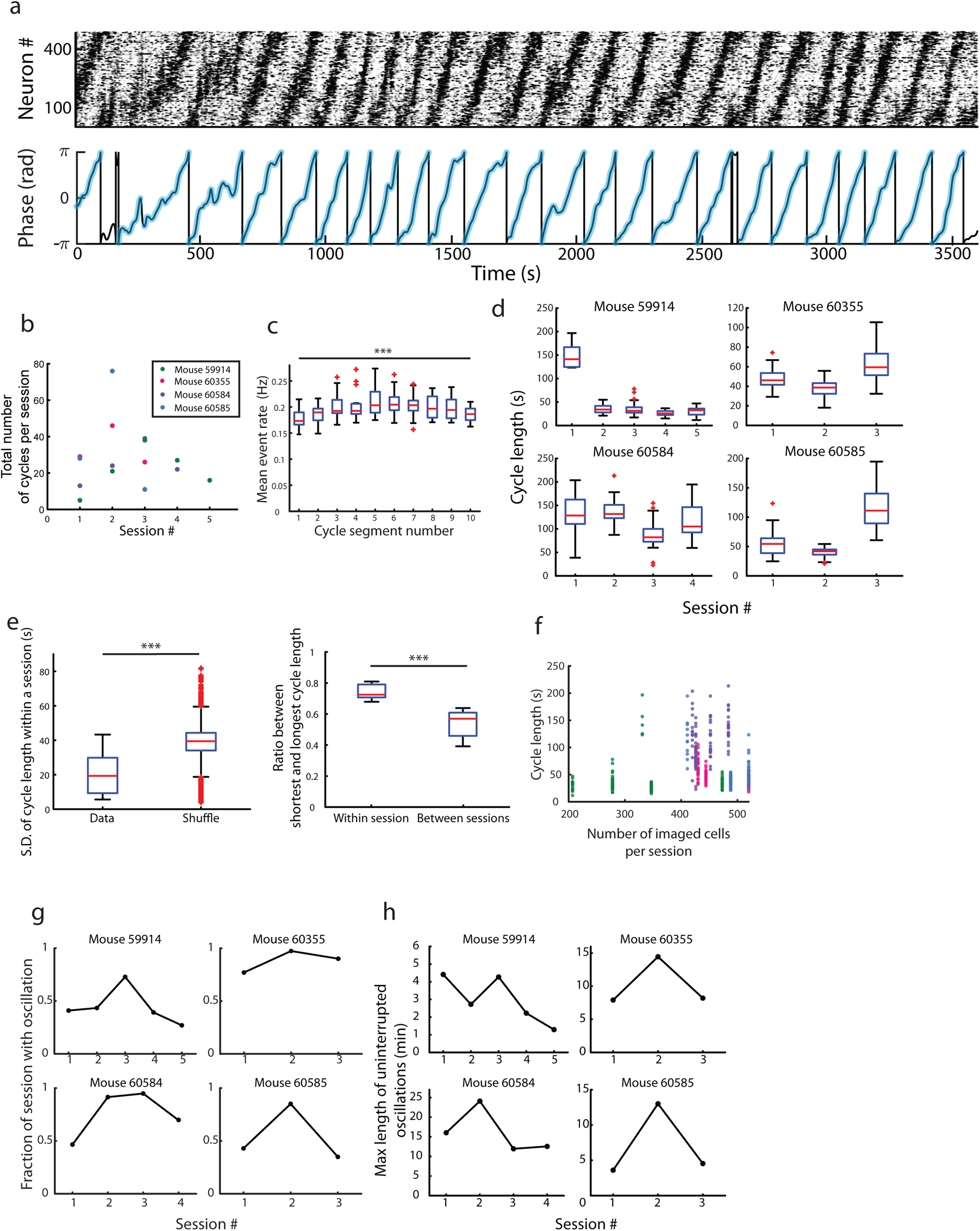
Identification of individual cycles and population oscillation characterization. **a.** Top: Raster plot of the PCA-sorted matrix of calcium activity of the example session in Fig. 2a. Bottom: Phase of the oscillation calculated on the session shown in the top panel is shown in black, and phase of individual cycles is colored in cyan. During one cycle of the population oscillation the phase of the oscillation traversed [*−π*, *π*) rad. To identify individual cycles, first the phase of the oscillation was calculated across the entire session, second discontinuities in the succession of such phases were identified and used to extract putative cycles and third, putative cycles were classified as cycles if the phase of the oscillation progressed smoothly and in an ascending manner, allowing for the exception of small fluctuations (lower than 10% of 2*π*, e.g. as in the sequence at 500 s). Points of sustained activity were ignored. Fractions of cycles in which the phase of the oscillation traversed 50% or more of the range [*−π*, *π*) rad were also analysed (for example at the beginning of the session). **b.** Total number of individual cycles per session, across 15 oscillatory sessions. Animal number is color-coded. **c.** Box plot showing mean event rate as a function of cycle segment for all 15 oscillatory sessions. Each cycle was divided into 10 segments of equal length, and for each cycle segment the mean event rate was calculated as the total number of calcium events across cells divided by the length of the segment and the number of recorded cells. Red lines indicate median across sessions, the bottom and top lines in blue indicate lower and upper quartiles, respectively. The length of the whiskers indicates 1.5 times the interquartile range. Red crosses show outliers that lie more than 1.5 times outside the interquartile range. The mean event rate remained approximately constant across the length of the cycle. While a non-parametric analysis revealed an overall difference (*n* = 15 oscillatory sessions per segment, *p*=0.0052, *χ*2=23.49, Friedman test), the rate change from the segment with minimum to maximum event rate was no more than 18% and there were no significant differences in the event rate between pairs of segments (Wilcoxon rank-sum test with Bonferroni correction, p>0.05 for all pairs). *** *p* < 0.001, ** *p* < 0.01, * *p* < 0.05, n.s. *p* > 0.05. **d.** Box plot of cycle length for each cycle of the oscillation, for the 15 oscillatory sessions. Note the relatively fixed length of cycles in individual sessions. Symbols as in (c). **e.** Left: Box plot of the standard deviation of cycle length within a session, in experimental and shuffled data. The standard deviation of cycle length is smaller in the experimental data (*n* = 15 oscillatory sessions, 7500 shuffle realizations, *p* = 1.8 × 10^−7^, *Z* = 5.08, one-tailed Wilcoxon rank-sum test). Right: Box plot of the ratio between the shortest cycle length and the longest cycle length for all pairs of cycles within and between sessions. This fraction is larger for cycle pairs in the within-session group (*n* = 15 oscillatory sessions, the mean fraction per session and group was calculated separately, *p* = 1.7 × 10^−6^, *Z* = 4.64, one-tailed Wilcoxon rank-sum test). Notice that for each cycle pair, the larger this ratio, the more similar the length of the cycles are. **f.** The cycle length is not correlated with the number of recorded cells in the session (*n* = 421 cycles across 15 oscillatory sessions, *p* = 0.02, *p* = 0.64, Spearman correlation). Each dot is a cycle. Animal number is color-coded as in (b). **g.** Fraction of the session in which the MEC population engaged in the oscillation. Session length was 30 min for mice 59914 and 60355, and 60 min for mice 60584 and 60585. **h.** Duration of the longest epoch with uninterrupted population oscillation. Only epochs that met the strict criterion of no separation between cycles were considered.

**Extended data Figure 6.**
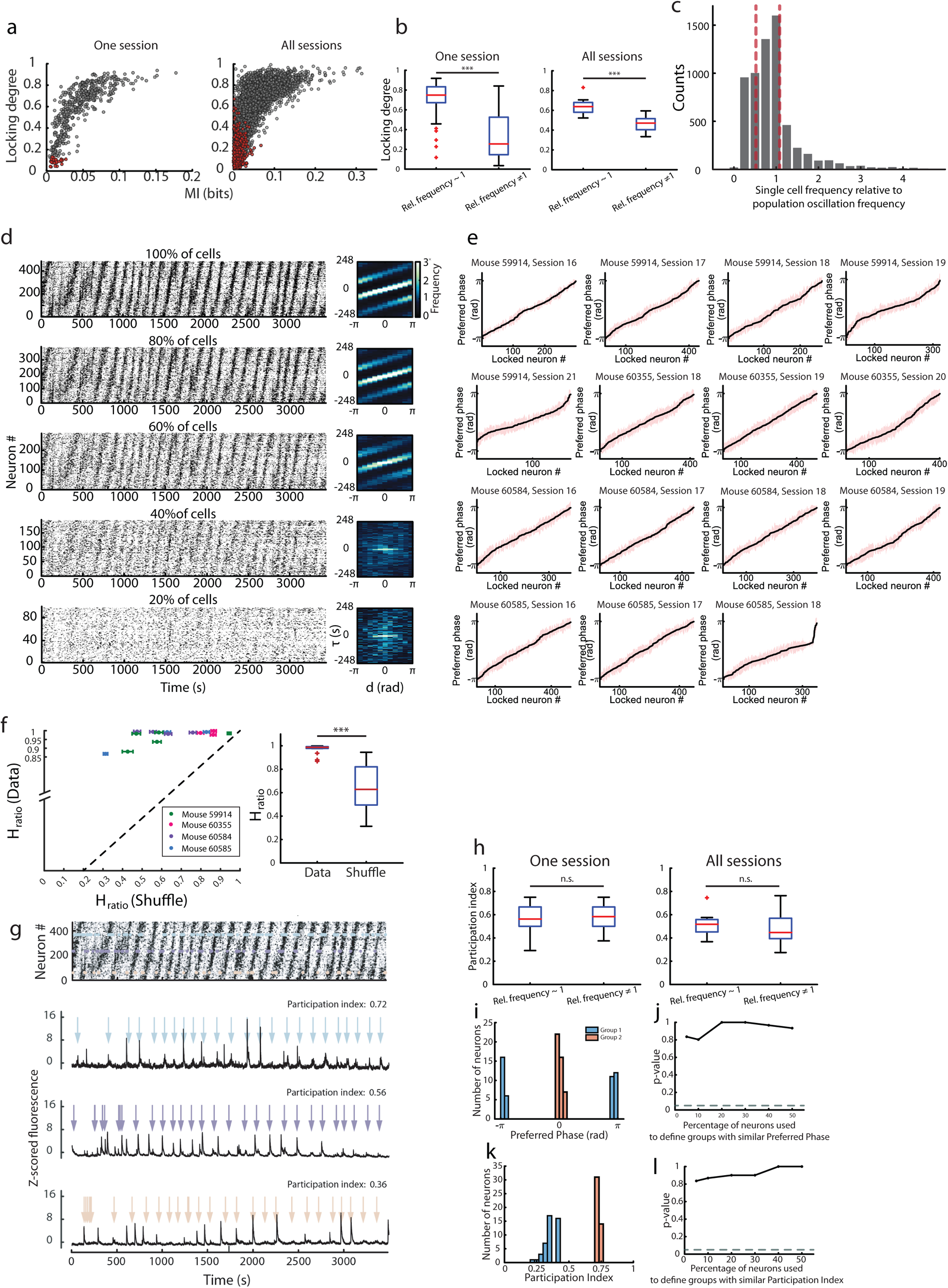
Characterization of locking degree and participation index. **a.** Consistency between two measures of phase locking for individual neurons. The locking degree was calculated for each cell as the length of the mean vector over the distribution of oscillation phases ([-π,π) rad) at which the calcium events occurred (bin size = 129 ms). The locking degree was consistent with the mutual information between the calcium event counts and the phase of the oscillation (bin size = 0.52 s). Scatter plots show the relation between the two measures, with each dot representing one neuron. Left: Data from the example session in Fig. 2a (n = 484 cells). Right: All neurons from all 15 oscillatory sessions are pooled (n = 6231 cells). Red dots indicate neurons that did not meet criteria for locking. The consistency between the two measures strengthens the conclusion that the vast majority of the neurons in MEC are locked to the population oscillation. **b.** Left: Box plot comparing locking degree for cells with an oscillatory frequency that was similar (relative frequency ~ 1) or different (relative frequency *≠* 1) from the frequency of the population oscillation in the example session in Fig. 2a (*n* = 48 cells in each group, *p* = 3.4× 10^−11^, *Z* = 6.63, Wilcoxon rank-sum test). Right: As left panel but for the locking degree across all 15 oscillatory sessions, including the example in the left panel (*n* = 15 sessions, *p* = 2.8× 10^−5^, *Z* = 4.19, Wilcoxon rank-sum test). Ten per cent of the total number of cells was used to define each of the groups with similar (relative frequency ~ 1) and different (relative frequency *≠* 1) oscillatory frequency as compared to the population oscillation frequency. Relative frequency was calculated for each cell as the oscillatory frequency of the cell’s calcium activity divided by the oscillatory frequency of the population oscillation in the session. Symbols as in Fig. 3b. Note that cells with relative frequency similar to 1 are more locked to the phase of the oscillation. For all percentages considered to define similar and different groups (5, 10, 20, 30, 40, and 50%) the p-values were significant. **c.** Histogram showing the distribution of single-cell oscillatory frequency divided by the population oscillation frequency of the session (n = 6231 cells pooled across 15 oscillatory sessions). A value of 1.0 indicates that single-cell and population frequency coincide. The left and right dashed lines indicate 25^th^ (0.52) and 75^th^ (1.08) percentiles respectively. Note that for approximately half of the data the oscillatory frequency is very similar at single-cell and population level. **d.** The population oscillation remains visible after excluding increasing fractions of neurons and keeping only those with the lowest locking degree. Each row shows a PCA-sorted raster plot (left, symbols as in Fig. 2b) and the corresponding joint distributions of the time lag *τ* that maximizes the correlation between the calcium activity of neuron pairs and their distance *d* in the PCA sorting (right, symbols as in Fig. 2f). The fraction of included neurons is indicated on top of the raster plot. For building the raster plots neurons were sorted according to their locking degree value and neurons with the highest locking degrees were removed. **e.** Distribution of preferred phases (the mean phase at which the calcium events occurred) in the population of locked neurons for all 15 oscillatory sessions. Black line indicates the preferred phases; red intervals indicate one standard deviation (calculated over the oscillation phases at which the calcium events of an individual cell occurred). Neurons are sorted according to their preferred phase in an ascending manner. The preferred phases cover the entire range of phases from −π to π. **f.** Phase preferences are distributed evenly across the MEC cell population. Left: The nearly-flat nature of the phase distribution is illustrated by comparing the entropy of the distribution of preferred phases in recorded (*y* axis) and shuffled data (*x* axis). H_ratio_ is the entropy of the distribution of preferred phases (calculated as in e) estimated from the data and divided by the entropy of a flat distribution (H_ratio_ = 1 if the distribution of preferred phases is perfectly flat, H_ratio_ = 0 if all neurons have the same preferred phase). Each point in the scatterplot indicates one session (15 sessions). Horizontal error bars indicate one S.D across shuffled realizations. The black dashed line indicates identical values for recorded and shuffled data. Animal number if color-coded. Notice the discontinuity in the *y* axis between 0 and 0.85. H_ratio_ is substantially larger for recorded data than for shuffled data. Right: Box plot of H_ratio_ for recorded and shuffled data. For each session the 1000 shuffled realizations were averaged (*n* = 15 oscillatory sessions, *p* = 6 × 10^−6^, *Z* = 4.52, Wilcoxon rank-sum test). Symbols as in Fig. 3b. **g.** Three example neurons from the example session in Fig. 2a. Top: Raster plot of the calcium matrix shown in Fig. 2a. Calcium events from the neuron with high participation index (PI, 0.72) are highlighted in light blue; from the neuron with intermediate PI (0.56) are highlighted in purple; from the neuron with low PI (0.36) are highlighted in orange. Bottom three panels: Z-scored fluorescence calcium signals as a function of time from the above neurons with high (top), intermediate (middle), and low (bottom) PIs. Colored arrows represent the time points at which the population oscillation is at the neuron’s preferred phase. Notice how the neuron with high PI tends to exhibit a peak in the calcium signal for most of the cycles. Neurons with intermediate and low PIs demonstrate the same but to a lesser extent, with the calcium signal not peaking in each cycle. **h.** Similar to (b), but for the participation index. Left: Data from the example session shown in Fig. 2a (*n* = 48 cells in each group, *p* = 0.51, *Z* = 0.66, Wilcoxon rank-sum test). Right: As left panel but for data pooled across 15 oscillatory sessions. The mean participation index was calculated for each group (“relative frequency ~ 1” and “relative frequency *≠* 1”) and each session separately and the data was then pooled across sessions (*n* = 15 sessions, *p* = 0.56, *Z* = 0.58, Wilcoxon rank-sum test). For all percentages considered to define the similar and different groups (5, 10, 20, 30, 40, and 50%) the p-values were non-significant. **i.** Histogram of preferred phases for the two groups of cells used to quantify the anatomical distribution of preferred phases for the example session in Fig. 2a. Cells in group one (two) had preferred phase ~ *π* rad (~ 0 rad). Each group had 45 locked cells, which is approximately ten per cent of the total number of locked cells in that session (454). Group one: blue; group two: orange. Distances between cells in group one (similar preferred phase), or between one cell in group one and one cell in group two (different preferred phase) were calculated. **j.** p-value for the difference in anatomical distance between the groups of cell pairs with similar preferred phase or different preferred phase (defined as in panel i), as a function of the percentage of cells used to build the groups of cells. The p-value was obtained through a Wilcoxon rank-sum test ran on the anatomical distances between cells with similar preferred phase (cells in group one) and cells with different preferred phase (distance between one cell in group 1 and one cell in group 2, for all pairs of cells). For all percentages considered (5, 10, 20, 30, 40, and 50%), the mean distances for the similar and the different classes were computed for each session. The means were then pooled across sessions (n = 15 oscillatory sessions). The dashed line indicates a level of significance of 0.05. Note that all p-values are much larger than the level of significance. **k.** Similar to (i) but for participation indexes. Cells in group one (two) had small (large) participation index. **l.** Similar to (j) but for the participation indexes. Symbols as in (j).

**Extended data Figure 7.**
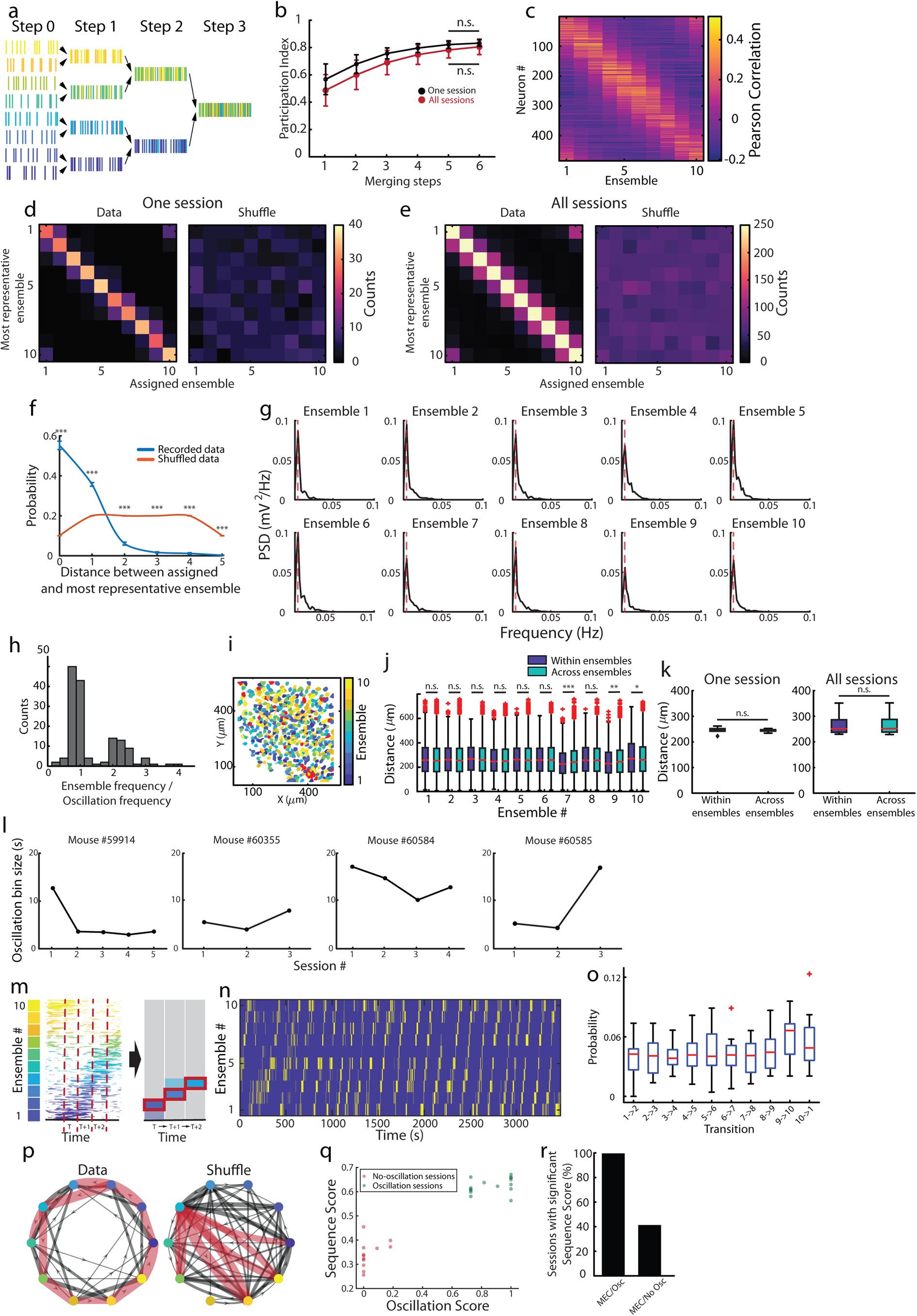
The population oscillation consists of periodic sequences of ensemble activation. **a.** Schematic of calcium activity merging steps. We began by sorting the neurons according to the PCA method. Next, in successive iterations, or merging steps, we added up the calcium activity of pairs of consecutive neurons (merging step = 1) or consecutive ensembles (merging step > 1). **b.** Participation index (PI) as a function of merging step (mean ± S.D.). Black trace, example session in Fig. 2a; red trace, all 15 oscillatory sessions. The more neurons per ensemble, the higher the participation index of the ensemble. Note that the participation index plateaus after 5 merging steps, which corresponds to approximately 10 ensembles (Wilcoxon rank-sum test to compare the participation indexes in merging steps 5 and 6; Black trace: *n* =30 PIs in merging step 5, *n* =15 PIs in merging step 6, *p* = 0.23, *Z* = 1.20; Red trace: *n* =15 PIs in merging step 5 and 6, PIs of each merging step were averaged for each session separately, *p* = 0.14, *Z* = 1.49). **c.** Tuning of single cell calcium activity to ensemble activity calculated as the Pearson correlation between the calcium activity of each neuron and the activity of each ensemble for the example session in Fig. 2a. Ensemble activity was calculated as the mean calcium activity across neurons in the ensemble. Each row is the tuning curve of one neuron, and neurons are sorted according to the PCA method. For each neuron, the calcium activity was positively correlated with a small subset of consecutive ensembles, and negatively correlated with the others. Pearson correlation is color-coded. **d.** The relationship between the calcium activity of each neuron and the activity of each ensemble was expressed by a Pearson correlation, as in (c). By repeating this calculation for all neurons across all ensembles, we could identify, for each neuron, the most representative ensemble (the one with maximal Pearson correlation). Left: 2D histogram of the most representative ensemble of each neuron and the ensemble it was assigned to based on the PCA sorting. Data are for the example session in Fig. 2a. Each count is a neuron; counts are color-coded (484 cells). Right: The same 2D histogram calculated on one shuffled realization of the data for the example session in Fig. 2a (484 cells). In the left diagram, note that the method for assigning cells into ensembles based on the PCA sorting correctly recovers the dependency between cells’ calcium activity and ensemble activity (higher number of counts along the diagonal). **e.** Same as (d), but for all neurons across all 15 oscillatory sessions (left, n = 6231), or one shuffled realization of the data (right, n = 6231). **f.** Probability distribution showing, for recorded data and shuffled data, the distance, in numbers of ensembles, between the assigned ensemble based on the PCA sorting and the most representative ensemble (as in d). The probability was calculated as the number of times that one given distance was observed in one session divided by the total number of recorded cells. Each count was one neuron. Note that the distance between the most representative ensemble and the assigned ensemble based on the PCA sorting reflects the periodic boundary conditions in ensemble activation and ranges from 0 to 5 (*x* axis). 500 shuffled realizations per session were averaged and compared to the mean distance per session in the recorded data. The probability of finding small distances (lower than 2) was larger in the recorded data (*n* = 15 sessions, for distances of 0 to 5 ensemble: *p ≤* 3.4 × 10^−6^, range of *Z*: 4.64 to 4.67; Wilcoxon rank-sum test), suggesting that single cell calcium activity was maximally correlated with the activity of the ensemble it was assigned to. Blue, recorded data; orange, shuffled data. Error bars indicate S.E.M. **g.** Ensemble activity oscillated at the same frequency as the population oscillation. Ensemble activity was calculated as the mean calcium activity across neurons in the ensemble. Power spectral density was calculated on the activity of each of the ten ensembles from the example session in Fig. 2a. Ensemble frequency was calculated as the peak frequency of the PSD, population oscillation frequency was computed as the total number of cycles (24 in this session) normalized by the amount of time in which the network engaged in the oscillation (~3600 s). The dashed line indicates the frequency of the population oscillation. Note that the dashed lines coincide with the peak of the PSD. **h.** Histogram showing the ratio between ensemble oscillatory frequency and population oscillation frequency in the session (calculated as in panel g; n = 150 data points given by 10 ensembles in each of the 15 oscillatory sessions). Each count is one ensemble. Note the two peaks at 1 and 2, indicating that ensembles tend to oscillate at the frequency of the population oscillation, or at an integer multiple of it. **i.** Anatomical distribution of recorded neurons for the example session in Fig. 2a. The ensemble each neuron has been assigned to based on the PCA sorting is color-coded. Neurons indicated in red were not locked to the phase of the oscillation. Note that ensembles are anatomically intermingled. Dorsal MEC on top, medial on the right, as in Extended data Fig. 1. **j.** Box plot of pairwise anatomical distance between neurons within an ensemble and between those neurons and the rest of the imaged neurons, i.e. across ensembles. Data are shown for each ensemble of the session in (i) (Wilcoxon rank-sum test to compare the within and across group distances for each ensemble separately; *n* = 1125 pairwise distances in the within ensemble group, except for ensemble 10, in which *n* = 1326; *n* = 20928 pairwise distances in the across ensemble group, except for ensemble 10, in which *n* = 22464, 0.0005 *≤ p ≤* 0.9528, 0.06 *≤ Z ≤* 3.50). Symbols as in Fig. 3b. Purple, distances between cells within one ensemble; green, distances between cells in different ensembles. **k.** Box plots of pairwise anatomical distance between neurons within one ensemble and across ensembles for the example session in (j) (left, *n* = 10 ensembles, *p* = 0.57, *Z* = 0.57, Wilcoxon rank-sum test) and across 15 oscillatory sessions including the example session in (j) (right). For each session the means for each of the “within” and “across” groups were computed across ensembles (*n* = 15 oscillatory sessions, *p* = 0.93, *Z* = 0.08, Wilcoxon rank-sum test). Symbols as in (j). **l.** To quantify the temporal progression of the population activity at the time scale at which the population oscillation evolved, we calculated, for each session, an oscillation bin size. This bin size is proportional to the inverse of the peak frequency of the PSD calculated on the phase of the oscillation, and hence captures the time scale at which the oscillation progresses. The oscillation bin size is shown for each of the 15 oscillatory sessions. **m.** Schematic of the method for quantifying temporal dynamics of ensemble activity. For each session and each ensemble we calculated the mean ensemble activity at each time bin (oscillation bin size). Only the ensemble with the highest activity within each time bin (red rectangle) was considered. The number of transitions between ensembles in adjacent time bins divided by the total number of transitions was used to calculate the transition matrices in Fig. 4b. **n.** The ensemble with the highest activity in each time bin, indicated in yellow and calculated as in (m), plotted as a function of time for the example session in Fig. 2a. All other ensembles are indicated in purple. Notice that the transformation in (m) preserves the population oscillation. **o.** Box plot showing transition probabilities between consecutive ensembles for all 15 oscillatory sessions. The probabilities remain approximately constant across transitions between ensemble pairs (*n* = 15 oscillatory sessions per transition, *p* = 0.56, *χ*2 = 7.77, Friedman test), and there were no significant differences between pairs of transitions (Wilcoxon rank-sum test with Bonferroni correction, *p* > 0.05 for all transitions). Symbols as in Fig. 3b. **p.** We further visualized the structure of the transitions in Fig. 4b by using the transition matrix as an adjacency matrix to build a directed weighted graph. Nodes indicate ensembles (color-coded as in m). Edges (lines) between any two nodes represent the transition probabilities between any two ensembles. The thickness of the edge is proportional to the value of the transition probability, while the arrows on each edge indicate the directionality of the transition. Red edges indicate edges whose associated transition probability is significant. Edges with significant transition probability were only found between consecutive or nearby nodes as well as between the nodes corresponding to ensemble 1 and 10, once again mirroring the periodic boundary conditions in ensemble activation. In shuffled realizations of the data there were edges that corresponded to significant transition probabilities, but those were not between neighboring nodes. **q.** Scatter plot showing relation between oscillation score and sequence score. The oscillation score quantifies the extent to which the calcium activity of single cells is periodic and ranges from 0 (no oscillation) to 1 (oscillation). The sequence score quantifies the probability of observing sequential activation of 3 or more ensembles. Each dot corresponds to one session. The sequence score increases with the oscillation score, and is highest for oscillatory sessions. Note that non-oscillatory sessions display non-zero values of sequence score, indicating the presence of sequential ensemble activity also in sessions below criteria for oscillation. **r.** Percentage of sessions with significant sequence score in sessions classified as oscillatory vs non-oscillatory. In MEC sessions with oscillations, 100% (15 of 15) of the sessions showed significant sequence scores, while in MEC sessions without oscillations, 41% (5 of 12) of the sessions demonstrated significant sequence scores. For corresponding raster plots, see Extended data Fig. 4a.

**Extended data Figure 8.**
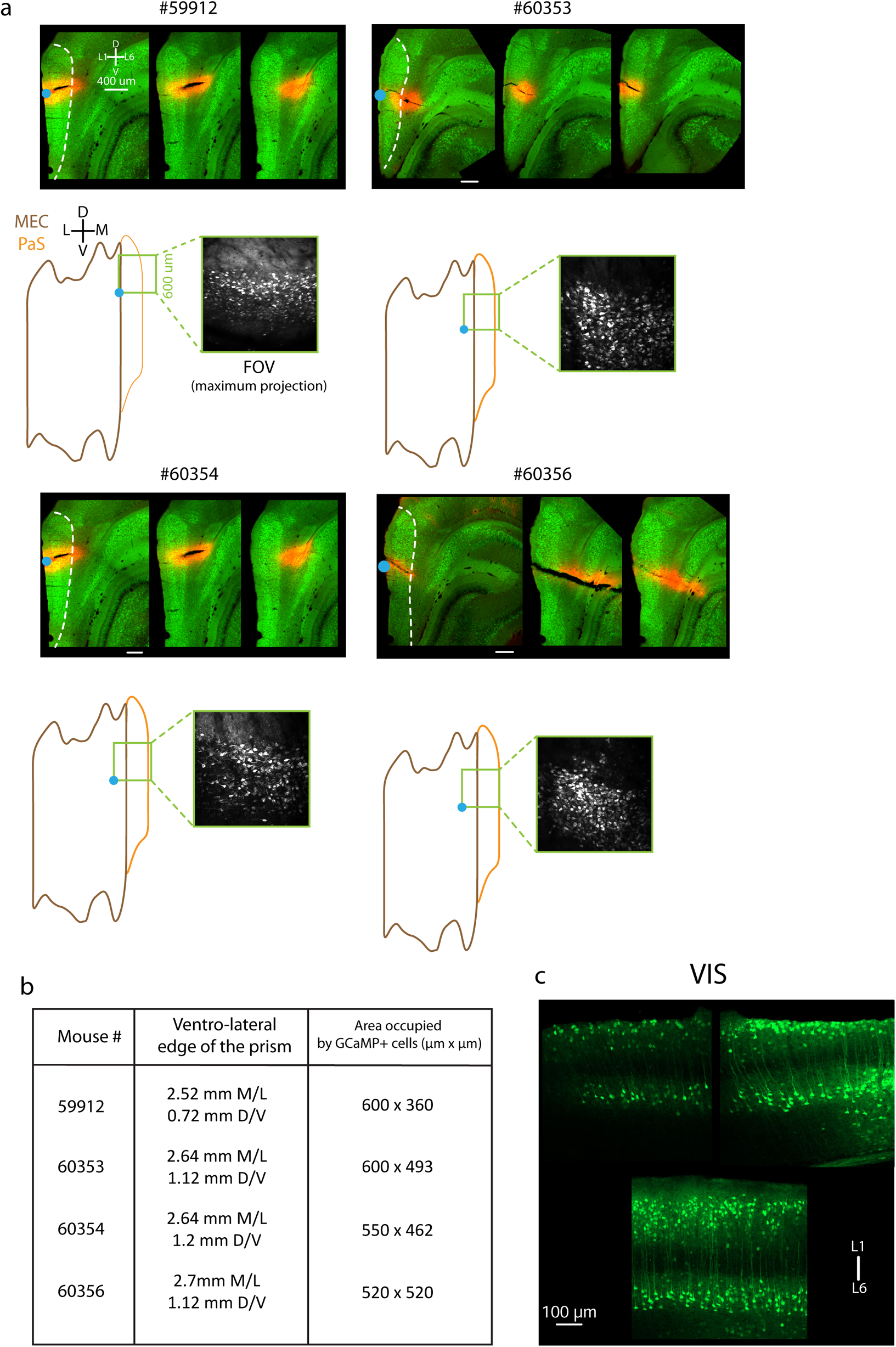
Histology showing imaging location in animals with FOVs in parasubiculum and visual cortex. **a.** Histological determination of prism location in parasubiculum-implanted mice. Top: Maximum intensity projection of 50 μm thick sagittal brain sections (sections acquired with an LSM 880, 20x). Three consecutive sections from the same mouse are shown, from the most lateral (left) to the most medial (right). Green is GCaMP6m signal, while red is Di L signal (used to demarcate ventrolateral corner of the prism, as in Extended data Fig. 1). Scale bar is 400 μm. The white stippled line encapsulates the superficial layers of the parasubiculum (PaS). Dorsal PaS on top, layer 1 on the left. Bottom: Estimated location of the field of view (FOV) on a flat map encompassing MEC (brown outline) and PaS (yellow outline). The blue dot marks the location of the pin used to demarcate the most lateral-ventral border of the prism, while the green square inset shows the microscope FOV. Inset images show mean (left) and maximum (right) intensity projections of the FOV. Dorsoventral (DV), and mediolateral (ML) axes are indicated. **b.** Location of the ventro-lateral edge of the prism in stereotactic coordinates, and area of the FOV occupied by cells expressing GCaMP6m for each PaS-imaged animal. **c.** Histological determination of imaging location in the visual cortex (VIS) in mice that underwent calcium imaging. Green is GCaMP6m signal. Images are taken from coronal slices, and zoomed in on visual cortex (Scale bar is 100 μm; L1 at the top, L6 at the bottom). Dorsal pole of the brain is on top. Maximum intensity projection, LSM 880, 20x.

**Extended data Figure 9.**
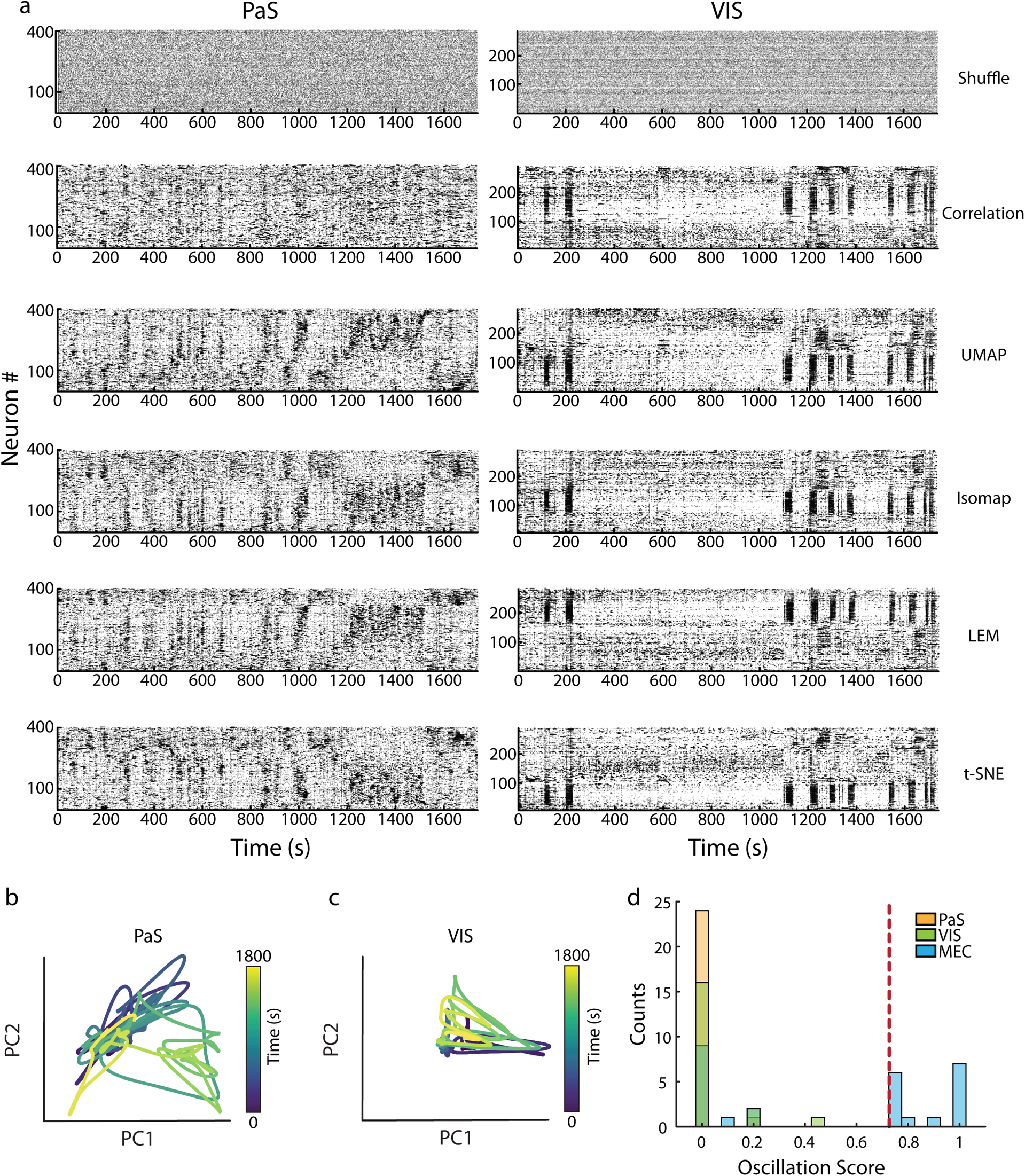
Lack of population oscillations in parasubiculum and visual cortex. **a:** Alternative sorting methods, as in Extended data Fig. 3c, but applied to sessions recorded in the PaS (left) or VIS (right). The PCA sorting method applied to temporally shuffled data did not unveil a population oscillation (first row). No population oscillation was recovered when neurons were sorted according to their correlation values (second row), or according to different dimensionality reduction techniques (UMAP, Isomap, LEM, t-SNE). Each row of each raster plot shows the calcium activity of a single neuron, with activity plotted as a function of time, as in Fig 2a. Every dot indicates that one neuron was active at one specific time bin (bin size = 129 ms). Sequence scores and oscillation scores are presented in Fig 6c,d. **b,c.** Projection of the neural activity onto the low-dimensional embedding defined by the first two principal components obtained from applying PCA to the matrix of calcium activity of the PaS session (b) and the VIS session (c) shown in Fig. 6a. Bin size = 8.5 s. Note lack of obvious ring topology. Time is color-coded. **d.** Distribution of oscillation scores for the entire data set, as in Extended data Fig. 4c (19 VIS sessions, 25 PaS sessions, 27 MEC sessions of which 15 were classified as oscillatory). Dashed line indicates threshold for classifying sessions as oscillatory with reference to the MEC data. Note that the bars for different brain regions sometimes overlap, and that bars are colored with transparency for visualization purposes (e.g. for sessions in PaS with oscillation score 0, the count is 24).

**Extended data Figure 10.**
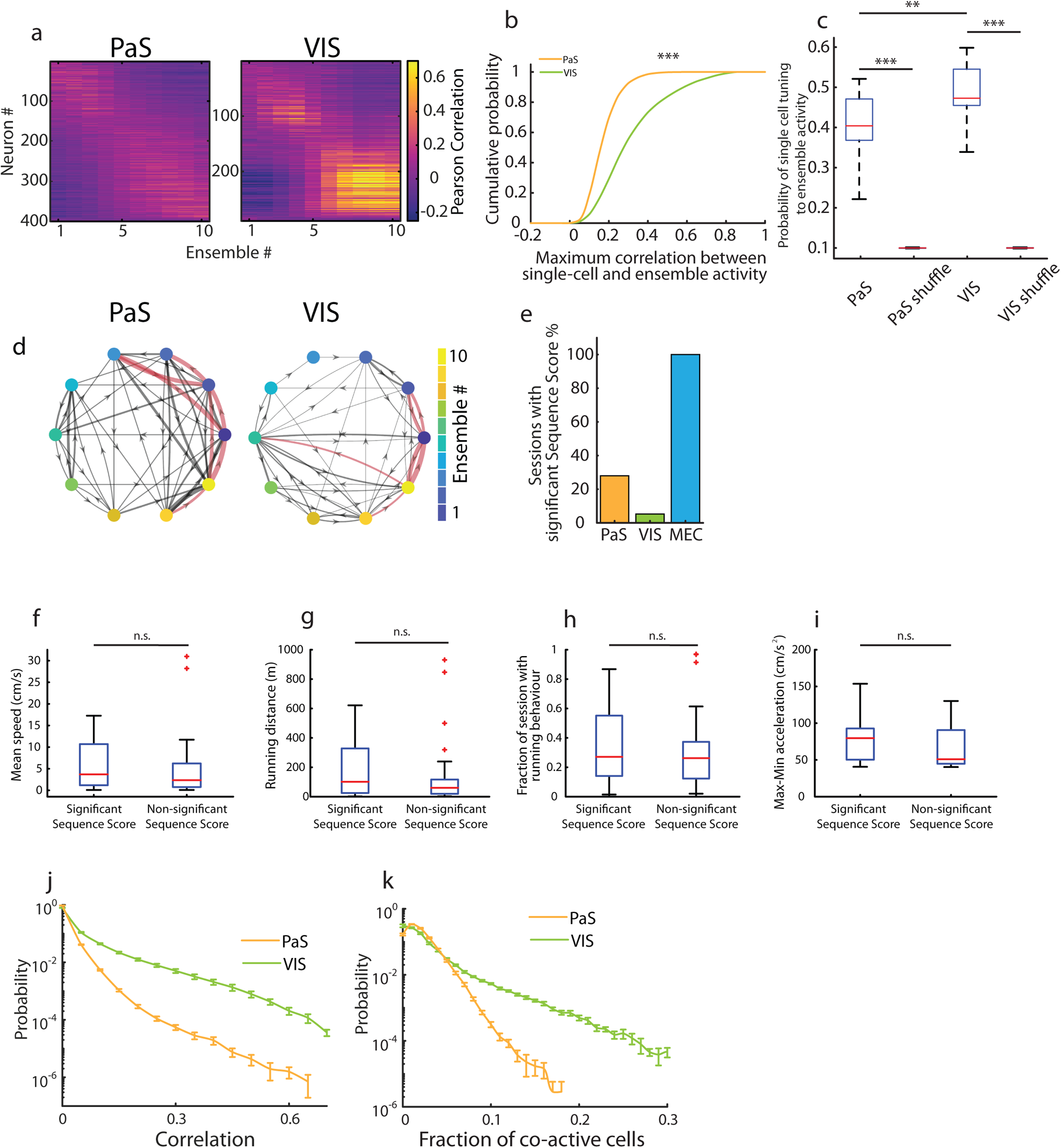
Population activity is less synchronized and more sequentially organized in PaS than VIS. **a.** Tuning of single cell calcium activity to ensemble activity expressed as the Pearson correlation between the calcium activity of each neuron and the activity of each ensemble, shown for the PaS (left, 402 cells) and the VIS (right, 289 cells) example sessions presented in Fig. 6a. Each row is the tuning curve of one neuron, and neurons are sorted according to the PCA method. Color indicates Pearson correlation. Note that the VIS session exhibits a cluster of high correlation values for ensembles 5-10, which might indicate the presence of high co-activity in cells allocated to those ensembles. **b.** Cumulative distribution of the maximum Pearson correlation value between each cell’s calcium activity and the ensemble activity. Data are for the same two example sessions as in (a). Note that VIS exhibits larger correlation between single-cell calcium activity and ensemble activity (*n* = 6037 VIS cells across 19 sessions, *n* = 10868 PaS cells across 25 sessions, *p* = 1, *D* = 0.4179, Kolmogorov Smirnov test). **c.** Probability that the ensemble a cell was assigned to based on the PCA sorting coincides with its most representative ensemble, calculated as the ensemble for which the Pearson correlation between the cell’s calcium activity and ensemble activity is maximal. The probability is calculated as the fraction of cells in an individual session for which the PCA-assigned ensemble and the most representative ensemble coincide. In the box plot; all PaS sessions (n=25) and all visual cortex sessions (n=19) were pooled. For each session in each brain area the matrix of calcium activity was shuffled 500 times, next the PCA-assigned and most representative ensemble were calculated for each cell in the session and the probability that these coincide was computed over all cells and averaged across shuffle realizations per session. Ensemble activity was representative of cells’ calcium activity in both brain areas (Wilcoxon rank-sum test comparing recorded and shuffled data for each brain area separately; PaS: *n* = 25 sessions, *p* = 1.42 × 10^−9^, *Z* = 6.05; VIS: *n* = 19 sessions, *p* = 1.48 × 10^−7^, *Z* = 5.25), although this effect was more pronounced for VIS than for PaS neurons (*n* = 25 PaS sessions, 19 VIS sessions; *p* = 0.0077, *Z* = 2.42, one-tailed Wilcoxon rank-sum test; median VIS = 0.47, median PaS = 0.40). Symbols as in Fig. 3b. **d.** Based on the transition matrices calculated in Fig. 6h, we built directed weighted graphs as in Extended data Fig. 7p. Red edges indicate edges whose associated transition probability is higher than the 95^th^ percentile of the transition probabilities obtained after temporally shuffling the data. **e.** Percentage of sessions with significant sequence score (MEC oscillatory sessions: 15 of 15, PaS: 7 of 25; VIS: 1 of 19). The sequence score quantifies the probability of observing sequential activation of 3 or more ensembles. **f.** Box plot of mean speed for sessions with and without significant sequence score. Mean speed was not different between these sessions (*n* = 21 sessions with significant sequence score and behavioural tracking synchronized to imaging: 13 MEC from which 10 were oscillatory + 7 PaS + 1 VIS; *n* = 30 sessions without significant sequence score and behavioural tracking synchronized to imaging: 1 MEC + 11 PaS + 18 VIS; *p* = 0.39, *Z* =0.85, Wilcoxon rank-sum test). Symbols as in Fig. 3b. **g.** Same as (f) but for total running distance (*p* = 0.42, *Z* =0.79, Wilcoxon rank-sum test). **h.** Same as (f) but for fraction of the session with running behaviour (*p* = 0.63, *Z* =0.47, Wilcoxon rank-sum test). **i.** Same as (f) but for the total amplitude of acceleration values, estimated as the maximum acceleration minus the minimum acceleration value observed in one session (*p* = 0.1, *Z* =1.62, Wilcoxon rank-sum test). **j.** Normalized distribution of the Pearson correlation values (absolute value) between the activity of cell pairs in VIS (green) and in PaS (yellow). Each dot indicates the mean across sessions (25 PaS sessions, 19 VIS sessions; all sessions in the data set were used, not only those with behavioural tracking synchronized to imaging), error bars indicate S.E.M. Probability is shown on a log-scale. **k.** Same as (j) but for the distribution of values of coactivity for all sessions recorded in PaS (yellow) and VIS (green). Coactivity was estimated for each session separately as the fraction of the recorded cells that was simultaneously active in 129 ms bins. Probability is shown on a log-scale.

**Movie 1**

Motion corrected video of one oscillatory session (session 17) from animal #60584. Time in seconds in top left, scale bar is 50 microns. The video was obtained by sampling every 10th frame of the motion-corrected Suite2p output, and using a 3 frame moving average (inter-frame time ~ 310 ms). The video shows 10 consecutive sequences.

## Methods

All experiments were performed in accordance with the Norwegian Animal Welfare Act and the European Convention for the Protection of Vertebrate Animals used for Experimental and Other Scientific Purposes, Permit numbers 6021, 6008, and 7163.

### Subjects

C57/Bl6 mice were housed in social groups of 2-6 individuals per cage, with access to nesting material and a planar running wheel. The mice were kept on a 12h light/12h darkness schedule in a temperature- and humidity-controlled vivarium. Food and water were provided ad libitum. The data were collected from a cohort of 12 animals (5 implanted in medial entorhinal cortex (MEC), 4 in parasubiculum (PaS), 3 in visual cortex (VIS)).

### Surgeries

Surgeries were performed according to a two-step protocol. During the first procedure, newborn pups or adult animals were injected in MEC/PaS or adult animals were injected in VIS with a virus carrying a construct for the expression of the calcium indicator GCaMP6m. The virus (for all injections: AAV1-Syn-GcaMP6m; titer: 3.43e13 GC/ml, Cat#AV-1-PV2823, UPenn Vector Core, University of Pennsylvania, USA) was diluted 1:1 in sterile DPBS (1X Dulbecco’s Phosphate Buffered Saline, Gibco, ThermoFisher). During the second procedure, two weeks later, a microprism was implanted to gain optical access to infected neurons located in MEC and PaS, or a glass window was inserted to obtain similar access in VIS.

For all surgeries, anesthesia was induced by placing the subjects in a plexiglass chamber filled with isoflurane vapor (5% isoflurane in medical air, flow of 1 l/min). Surgery was performed on a heated surgery table (38°C). Air flow was kept at 1 l/min with 1.5–3% isoflurane as determined from physiological monitoring of breathing and heartbeat. The mice were allowed to recover from surgery in a heated chamber (33°C) until they regained complete mobility and alertness.

#### Virus Injection and microprism implantation in MEC and PaS

In the first surgical procedure, newborn pups received injections of AAV1-Syn-GCaMP6m one day after birth^99^. Analgesics were provided immediately before the surgery (Rymadil, Pfizer, 5 mg/kg). Pre-heated ultrasound gel (39°C, Aquasonic 100, Parker) was generously applied on the pup’s head in order to create a large medium for the transmission of ultrasound waves. Real-time ultrasound imaging (Vevo 1100 System, Fujifilm Visualsonics) allowed for targeted delivery of the viral mixture to specific areas of the brain. During ultrasound imaging, the pup was immobilized through a custom-made mouth adapter. The ultrasound probe (MS-550S) was lowered to be in close contact with the gel and hence the pup’s head to allow visualization of the targeted structures. The probe was kept in place for the whole duration of the procedure via the VEVO injection mount (VEVO Imaging Station. Imaging in B-Mode, frequency: 40 MHz; power: 100%; gain: 29 dB; dynamic range: 60 dB). Target regions were identified by structural landmarks: the MEC or PaS were identified in the antero-posterior and medio-lateral axis by the appearance of the aqueduct of Sylvius and the lateral sinus. The target area for injection was comparable to a coronal section at ~-4.7 mm from bregma in the adult animal. The solution containing the virus (250 ± 50 nl per injection) was injected in the target regions via beveled glass micropipettes (Origio, custom made; outer tip opening: 200 μm; inner tip opening: 50 μm) using a pressure-pulse system (Visualsonics, 5 pulses, 50 nl per pulse). The pipette tip was pushed through the brain without any incision on the skin, or craniotomy through the skull, and, to reduce the duration of the procedure, retracted immediately after depositing the virus in the target area. The anatomical specificity of the infection was verified by imaging serial sections of the infected hemispheres after experiment completion (see “Histology and reconstruction of field of view location”).

Two weeks after the viral injection, we performed a second procedure, in which a microprism was implanted to gain optical access to the superficial layers of MEC and PaS^100^. The implanted microprism was a right-angle prism with 2 mm side length and reflective enhanced aluminum coating on the hypotenuse (Tower Optical). The prism was glued to a 4mm-diameter (CS-4R, thickness #1) round coverslip with UV curable adhesive (Norland). On the day of surgery, mice were anesthetized with isoflurane (IsoFlo, Zoetis, 5 % isoflurane vaporised in medical air delivered at 0.8-1 l/min) after which two analgesics were provided through intraperitoneal injection (Metacam, Boehringer Ingelheim, 5 mg/kg or Rimadyl, Pfizer, 5 mg/kg, and Temgesic, Indivior, 0.05-0.1 mg/kg) and one local analgesic was applied underneath the skin covering the skull (Marcain, Aspen, 1-3 mg/kg). Their scalp was removed with surgical scissors and the surface of the bone was dried before being generously covered with optibond (Kerr). To increase the thickness and stability of the skull and overall preparation, a thin layer of dental cement (Charisma, Kulzer) was applied on the exposed skull, except in the location above the implant, where a 4 mm-wide circular craniotomy was made. The craniotomy was positioned over the dorsal surface of the cortex and cerebellum, with the center positioned *~* 4 mm lateral from the center of the medial sinus, and above the transverse sinus just above the MEC and PaS. After the dura was removed above the cerebellum, the lower edge of the prism was slowly pushed in the empty space between the forebrain and the cerebellum, just posterior to the transverse sinus. The edges of the coverslip were secured to the surrounding skull with with UV-curable dental cement (Venus Diamond Flow, Kulzer). A custom-designed steel headbar was attached to the dorsal surface of the skull, centered upon and positioned parallel to the top face of the microprism. All exposed areas of the skull, including the headbar, were finally covered with dental cement (Paladur, Kulzer) and made opaque by adding carbon powder (Sigma Aldrich) until the dental cement powder became dark grey.

#### Virus injection and glass window implantation in VIS

In a different cohort of animals than those used for MEC/PaS imaging, we induced the expression of GCaMP6m in neurons of the adult VIS for subsequent imaging. We targeted the injection of the same AAV1-Syn-GCaMP6m viral solution used in the developing MEC and PaS to the primary visual cortex. On the day of surgery, 3-5 months old mice were anesthetized with isoflurane (IsoFlo, Zoetis, 5 % isoflurane vaporised in medical air delivered at 0.8-1 l/min) after which two analgesics were provided through intraperitoneal injection (Metacam, Boehringer Ingelheim, 5 mg/kg or Rimadyl, Pfizer, 5 mg/kg, and Temgesic, Indivior, 0.05-0.1 mg/kg) and one local anaesthetic was applied underneath the skin covering the skull (Marcain, Aspen, 1-3 mg/kg). The virus was injected at three locations in VIS, all of which were within the following anatomical ranges: 2.3-2.5 mm lateral from the midline, 0.9-1.3 mm anterior from lambda^101^. At each injection site, 50 nl of the virus was injected 0.5 mm below the dura and the pipette was left in place for 3-4 min to enable the virus to diffuse. The pipette was then brought to 0.3 mm below the dura and another 50 nl was injected. The pipette was then left in place for 5-10 min before retracting it completely. The speed of the injections was 5 nl/s.

Two weeks after the viral injection, a surgery to chronically implant a glass window on VIS was performed. The animals were handled as previously described for the prism surgery in MEC/PaS, including anesthesia, delivery of analgesics, and scalp removal. Optibond was applied to the exposed skull except in the location of the craniotomy. A 4 mm-wide craniotomy was made, centered on the virus injection coordinates, and a 4 mm glass window was placed underneath the skull edges of the craniotomy. The glass was slightly larger than the craniotomy, so after it was maneuvered in place, the upward pressure exerted by the brain secured it in place against the skull, thereby minimizing the presence of empty gaps that might favor tissue and bone regrowth. The edges of the window were secured with UV-curable dental cement and superglue before the positioning of the headbar as described for the MEC-PaS implantation. All exposed areas of the skull, including the headbar, were finally covered with dental cement (Paladur, Kulzer) that was made opaque by adding carbon powder (Sigma Aldrich) until the dental cement powder became dark grey.

### Self-paced running behavior under sensory-minimized conditions

Training of animals began 2 days after the prism implantation in MEC and PaS, and 12 days after the implantation of a cranial window in VIS. Mice were head-restrained by a headbar with their limbs resting on a freely rotating styrofoam wheel with a metal shaft fixed through the center. The radius of the wheel was *~*85 mm and the width 70 mm. Low friction ball bearings (HK 0608, Kulelager AS, Molde, NO) were affixed to the ends of the metal shaft and held in place on the optical table using a custom mount. This arrangement allowed the mice to self-regulate their movement. The position of the animal on the rotating wheel was measured using a rotary encoder (E6B2-CWZ3E, YUMO) attached to its center axis.

Step values of the encoder (4096 per full revolution, ~130 µm resolution) were digitized by a microcontroller (Teensy 3.5, PJRC) and recorded using custom python scripts at 40-50 Hz. Wheel tracking was triggered at the start of imaging and synchronized to the ongoing image acquisition through a digital input from the 2-photon microscope. In a subset of mice (3 out of 12; 2 implanted in MEC, 1 implanted in PaS), the precise synchronization was not available to us and these data were hence not used for comparison of movement and imaging data. A T-slot photo interrupter (EE-SX672, Omron) served as a lap (full-revolution) counter. Design and code of the wheel are publicly available under https://github.com/kavli-ntnu/wheel_tracker.

The self-paced task was performed under conditions of minimal sensory stimulation, in darkness, and with no rewards to signal elapsed time or distance run^36,37^. Prior to the imaging sessions, mice were accustomed to the setup through daily exposures over the course of two weeks (i.e., 15 sessions over 15 days, one session per day). In each session, after the mice were positioned on the wheel, they were gently head-restrained and free to run or rest for 30 or 60 min.

### 2-photon imaging in head-fixed animals

A custom-built 2-photon benchtop microscope (Femtonics, Hungary) was used for 2-photon imaging of the target areas (i.e., superficial layers of MEC, PaS, and VIS). A Ti:Sapphire laser (MaiTai Deepsee eHP DS, Spectra-Physics) tuned to a wavelength of 920 nm was used as the excitation source. Average laser power at the sample (after the objective) was 50–120 mW. Emitted GCaMP6m fluorescence was routed to a GaAsP detector through a 600 nm dichroic beamsplitter plate and 490-550 nm band-pass filter. Light was transmitted through a 16x/0.8NA water-immersion objective (Cat#MRP07220, Nikon) carefully lowered in close contact to the coverslip glued to the microprism (for MEC-PaS imaging) or above the coverslip in contact with the brain surface (for VIS imaging). For the microprism-implanted animals, the objective lens was aligned to the ventrolateral corner of the prism, to consistently identify the position of MEC and PaS across animals. Ultrasound gel (Aquasonic 100, Parker) or water was used to fill the gap between the objective lens and the glass coverslips. The software MESc (v 3.3 and 3.5, Femtonics, Hungary) was used for microscope control and data acquisition. Imaging time series of either ~30 min or ~60 min were acquired at 512×512 pixels (sampling frequency: 30.95 Hz, frame duration: ~32 ms; pixel size: either 1.78×1.78 µm^2^ or 1.18×1.18 µm^2^). Time series acquisition was initiated arbitrarily after the animal was head-restrained on the setup.

### Histology and reconstruction of field-of-view location

On the last day of imaging, after the imaging session, the mice were anesthetized with isoflurane (IsoFlo, Zoetis) and then received an overdose of sodium pentobarbital before transcardial perfusion with freshly prepared PFA (4% in PBS). After perfusion, the brain was extracted from the skull and kept in 4% PFA overnight for post-fixation. The PFA was exchanged with 30% sucrose to cryoprotect the tissue.

To verify the anatomical location of the imaged field of views (FOVs) in the microprism implanted animals, we used small, custom-made pins, derived from a thin piano wire coated with a solution of 1,1’-Dioctadecyl-3,3,3’,3’-Tetramethylindocarbocyanine Perchlorate (’DiI’; DiIC18(3)) (commercial name: Dil, ThermoFischer), to mark the location of the imaged tissue in relation to the prism footprint. A Dil-coated pin was inserted into the brain tissue at the location left empty by the prism footprint, and specifically targeted to the ventro-lateral corner of the footprint (see “Surgeries”). The pin was left in place to favor transfer of DiI from the metal pin to the brain tissue, and to leave a fluorescent mark on the location of the imaged FOV. After 30 to 60 seconds, the pin was removed and the brain was sliced on a cryostat in 30-50 µm thick sagittal sections. All slices were collected sequentially in a 24-well plate filled with PBS, before being mounted in their appropriate anatomical order on a glass slide in custom-made mounting medium. For confocal imaging, a Zeiss LSM 880 microscope (Carl Zeiss, Germany) was used to scan through the whole series of slices and locate the position of the DiI fluorescent mark. Images were then acquired using an EC Plan-Neofluar 20×/0.8 NA air immersion, 40×/1.3 oil immersion, or 63×/1.4 oil immersion objective (Zeiss, laser power: 2-15%; optical slice: 1.28–1.35 airy units, step size: 2 µm). Before acquisition, gain and digital offset were established to optimize the dynamic range of acquisition to the dynamic range of the GCaMP6m and DiI signals. Settings were kept constant during acquisition across brains. Based on the location of the red fluorescent mark, we could infer where, on the medio-lateral and dorso-ventral extent of the brain, the ventro-lateral corner of the microprism (and hence the 2-photon FOV aligned to it) was located.

We used the Paxinos mouse brain atlas^101^ to produce a reference flat map representing the medio-lateral and dorso-ventral extent of the MEC and PaS. Flat maps helped delineate the extent of the FOV that fell within the anatomical boundaries of either the MEC and adjacent PaS, and allowed for a standardized comparison across animals. For each imaged animal, we mapped the dorsoventral and mediolateral location of the DiI mark on the refence flat map (Extended data Fig. 1c). Animals were assigned to “MEC Imaging” or “PaS imaging” groups depending on the location of the FOV: a mouse would be further analysed as being part of the “MEC imaging” group if more than 50% of the area of the FOV occupied by GCaMP6m+ cells could be located in the MEC.

To verify the anatomical location of the FOVs in VIS in the glass-window implanted mice, we sliced the brain until we reached the anatomical coordinates at which the virus was infused (see “Surgeries”). Coronally cut slices of 50 µm thickness were collected sequentially in a 24 well plate, and immediately mounted in their appropriate anatomical order on a glass slide in custom-made mounting medium. For confocal imaging, a Zeiss LSM 880 microscope (Carl Zeiss, Germany) was used according to the same specification as described above for MEC/PaS.

### Analysis of imaging timeseries

Imaging timeseries data was analyzed using the Suite2p^39^ python library (https://github.com/MouseLand/suite2p). We used its built-in routines for motion correction, region of interests (ROI) extraction, neuropil signal estimation, and spike deconvolution. Non-rigid motion correction was chosen to align each frame iteratively to a template. Quality was assessed by visual inspection of the corrected stacks and built-in motion correction metrics. The Suite2p GUI was used to manually sub-select putative neurons based on anatomical and signal characteristics and to discard obvious artefacts that accumulated during the analysis, e.g., ROIs with footprints spanning large areas of the FOV, ROIs that did not have clearly delineated circumferences in the generated maximum intensity projection, or ROIs that were extracted automatically but showed no visible calcium transients.

Raw fluorescence calcium traces of each ROI were neuropil-corrected to create a fluorescence calcium signal “F_corr_” by subtracting 0.7 times the neuropil signal from the raw fluorescence traces. We used the Suite2p integrated version of non-negative deconvolution^38^ with tau=1 s to deconvolve F_corr_, yielding the basis for the binarized sequences that we refer to as the calcium activity (see section below “Binary deconvolved calcium activity and matrix of calcium activity”). To estimate the signal-to-noise-ratio (SNR) of each cell, we further thresholded the calcium activity (without binarization) at 1 standard deviation over the mean, yielding filtered calcium activity, and classified the remaining activity as noise. We additionally ensured that noise was temporally well segregated from filtered calcium activity by requiring datapoints classified as noise to be separated by at least one second before and ten seconds after filtered calcium activity. The SNR of the cell was then estimated as the ratio of the mean amplitude of F_corr_ during episodes of filtered calcium activity over the standard deviation of F_corr_ during episodes of noise. If no datapoints remained after the filtering of calcium activity, the cell was assigned a SNR of zero.

### Binary deconvolved calcium activity and matrix of calcium activity

In order to denoise the recorded fluorescence calcium signals and have good temporal resolution all analyses in the study were performed using the deconvolved calcium activity of the recorded cells. For each cell whose SNR was larger than 4, the deconvolved calcium activity (see “Analysis of imaging timeseries”) was downsampled by a factor of 4 by calculating the mean over time windows of ~129 ms (original sampling frequency = 30.95 Hz, sampling frequency used in the analyses = 7.73 Hz). Next, this signal was averaged over time and its standard deviation was calculated. A threshold equal to this average plus 1.5 times the standard deviation was used to convert the deconvolved calcium activity into a binary deconvolved calcium activity, such that all values above the threshold were set to 1 (“calcium events”), and all values below or equal to that threshold were set to 0. Unless stated otherwise, for all analyses throughout the study we used the deconvolved and binary calcium activity, to which for simplicity we refer to as “deconvolved calcium activity” or simply “calcium activity”. The calcium activity of all cells in a session with SNR > 4 was stacked to construct a binary “matrix of calcium activity” which had as many rows as neurons, and as many columns as time bins sampled at 7.73 Hz. The *population vectors* are the columns of the matrix of calcium activity.

### Autocorrelations and spectral analysis of single cell calcium activity

To determine if the calcium activity of single cells displays ultraslow oscillations, for each neuron the power spectral density (PSD) was calculated on the autocorrelation of its calcium activity. The PSD was computed using Welch’s method (“pwelch”, built-in Matlab function), with Hamming windows of 17.6 min (8192 bins of 129 ms in each window) and 50% of overlap between consecutive windows. Note that when calculating the PSD a large window was needed to identify oscillation frequencies << 0.1 Hz.

To visualize whether specific oscillatory patterns at fixed frequencies were present in the neural population, all autocorrelations from one session were sorted and stacked into a matrix, where rows are cells and columns are time lags. The sorting of autocorrelations was performed either (1) according to the maximum power of each PSD in a descending manner, or (2) according to the frequency at which each PSD peaked, in a descending manner. The frequency at which the PSD peaked was used as an estimate of the oscillatory frequency of the cell’s calcium activity.

### Correlation and PCA sorting methods

To determine whether neural population activity exhibits temporal structure we visualized the population activity by means of raster plots in which we sorted all cells according to different methods.

#### Correlation method

This method sorts cells such that those that are nearby in the sorting are more synchronized than those that are further away. First, each calcium activity was downsampled by a factor 4 by calculating the mean over counts of calcium events in bins of 0.52 s. The obtained calcium activity was then smoothed by convolving it with a gaussian kernel of width equal to four times the oscillation bin size, a bin size that was representative of the temporal scale of the population dynamics (see “Oscillation bin size”). The cross correlations between all pairs of cells were calculated using time bins as data points, and a maximum time lag of 10 time points, equivalent to ~5 s. This small time lag allowed us to identify near instantaneous correlation while keeping information about the temporal order of activity between cell pairs. The maximum value of the cross correlation between cell *i* and cell *j* was stored in the entry (*i*, *j*) of the correlation matrix *C*, which was a square matrix of N rows and N columns, where N was the total number of recorded neurons in the session with SNR > 4. If the cross correlation peaked at a negative time lag the value in the entry *(i*, *j*) was multiplied by −1. The entry with the highest cross correlation value was identified and its row, denoted by *i_max_*, was used as the lead cell for the sorting procedure and chosen to be the first cell in the sorting. Cells were then sorted according to the values in the entries *(i_max_*, *j*), *j* = 1,2, …, *N*, *j ≠ i_max_*, i.e. their correlations with the lead cell, in a descending manner.

#### PCA method

Computing correlations from the calcium activity or the calcium signals can be noisy due to variability in the frequency of the calcium events and fine tuning of hyperparameters (e.g. the size of the kernel used to smooth the calcium activity of all cells). To avoid this, we leveraged the fact that the periodic sequences of neural activity in the population oscillation constitute low dimensional dynamics with intrinsic dimensionality equal to 1, and sorted the cells based on an unsupervised dimensionality reduction approach (a similar approach was used in ref. 102). For each recording session, principal component analysis (PCA) was applied to the matrix of calcium activity (bin size = 129 ms; using Matlab’s built-in function “pca”), including all epochs of movement and immobility and using the rows (neurons) as variables and the columns (population vectors) as observations. The first two principal components (PCs) were kept, since two is the minimum number of components needed to embed non-linear 1-dimensional dynamics. Cells were sorted according to their loadings in PC1 and PC2, expecting that the relationship between these loadings would express the ordering in cell activation during the sequences.

The plane spanned by PC1 and PC2 was named the PC1-PC2 plane. In the PC1-PC2 plane, the loadings of each neuron (the components of the eigenvectors without being multiplied by the eigenvalues) defined a vector, for which we computed its angle 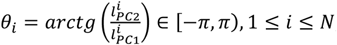, with respect to the axis of PC1, where 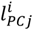 is the loading of cell *i* on *PCj*. Cells were sorted according to their angle *θ* in a descending manner.

Note that while we keep the first 2 PCs to sort the neurons, all PCs and the full matrices of calcium activity were used in the analyses (except for visualization purposes, e.g. see “Manifold visualization for MEC sessions”). Finally, note that because in PCA a PC is equivalent to −1 times the PC, the sorting and an inversion of the sorting are equivalent. The sorting was chosen so that sequences would progress from the bottom to the top in the raster plot.

The PCA method was used throughout the paper for sorting the recorded cells unless otherwise stated.

#### Random sorting of cell identities

A random ordinal integer ∈ [1, *N*], where *N* is the total number of recorded cells with SNR > 4, was assigned to each neuron without repetition across cells. Neurons were sorted according to those assigned numbers.

#### Sorting of temporally shuffled data

A shuffled matrix of calcium activity was built by temporally shuffling the calcium activity of each cell separately. For each cell, each time bin of the calcium activity was assigned a random ordinal integer ∈ [1, *T*] without repetition across time bins, where *T* is the total number of time bins (bin size = 129 ms), and time bins were ordered according to their assigned number.

The assignment of random ordinal integers was made separately for each cell, so that the obtained random orderings were not shared across cells. The PCA method was then applied to the shuffled matrix of calcium activity.

### Sorting methods based on non-linear dimensionality reduction techniques

The PCA method for sorting cells relies on a two-dimensional linear embedding. This linear embedding might not be optimal if the population vectors describe temporal trajectories that, despite being low-dimensional, lie on a curved surface. To take into account potential non-linearities, four additional sorting methods were implemented, based on the following non-linear dimensionality reduction techniques^103^: t-SNE, Laplacian Eigenmaps (LEM), Isomap, and UMAP^104^ (see parameters below). First, to express in the sortings the ordering of the cells during the slow temporal progression of the sequences, the four methods used a resampled matrix of calcium activity as input. To compute this matrix, for each session, we downsampled each calcium activity by a factor 4 by calculating its mean in bins of 0.52 s. The calcium activity of all cells was then smoothed by convolving them with a gaussian kernel whose width was given by the oscillation bin size (see “Oscillation bin size”). After applying t-SNE, LEM, Isomap or UMAP to the resampled matrix of calcium activity, we kept the first two dimensions obtained with each method, for the same reasons as presented for the PCA sorting method. To obtain the sorting, the following procedure was applied: We let Dim1 and Dim2 be the first two dimensions obtained with the chosen dimensionality reduction technique that we had applied to the resampled matrix. In analogy with the PCA method, the Dim1-Dim2 plane was spanned by Dim1 and Dim2 and for each cell the components on those dimensions defined a vector in this plane for which the angle *8* ∈ [*−π*, *π*) with respect to the axis of Dim1 was computed. Cells were then sorted according to their angles in a descending manner.

To apply t-SNE to the population activity we used a perplexity value of 50. First, we applied PCA to the resampled matrix of calcium activity, and then we used the projection of the neural activity onto the first 50 principal components as input to t-SNE. To apply LEM to the population activity, we used as hyperparameters k=15 and σ=2. Similarly, we used k=15 for running isomap. Finally, we used n_neighbors=30, min_dist=0.3 and correlation as metric for running UMAP.

We used the MATLAB implementation of UMAP^105^ and the Matlab Toolbox for Dimensionality Reduction (https://lvdmaaten.github.io/drtoolbox/). Finally, when displaying the raster plots that resulted from the different sortings, the first cell (located at the bottom of the raster plot) was always the same. This was accomplished by circularly shifting the cells in the different sortings such that the initial cell in all sortings coincided with the initial cell of the sorting obtained with the PCA method.

### Manifold visualization for MEC sessions

Sorting the cells and visualizing their combined neural activity through raster plots revealed the presence of oscillatory sequences of neural activity in the recorded data. To visualize the topology of the manifold underlying the oscillatory sequences of activity, both PCA and LEM were used.

PCA was applied to the matrix of calcium activity, which first had each row convolved with a gaussian kernel of width equal to 4 times the oscillation bin size (see “Oscillation bin size”). The manifold was visualized by plotting the neural activity projected onto the embedding defined by PC1 and PC2. In Fig. 2d (left) the neural activity of the entire session was projected onto the low-dimensional embedding. In Extended data Fig. 3d the neural activity corresponding to the concatenated epochs of uninterrupted population oscillation was projected onto the embedding.

For the LEM approach, first PCA was applied to the matrix of calcium activity, which was previously resampled to bins of 0.52 s as in “Sorting methods based on non-linear dimensionality reduction techniques”, and the first 5 principal components were kept. Next LEM was applied to the matrix composed of the 5 PCs, using as parameters k=15 and σ=2. We decided to keep 5 PCs prior to applying LEM to denoise the data, for which we leveraged the fact that sequences of activity constitute low-dimensional dynamics with intrinsic dimensionality equal to 1, and therefore truncating the data to the first 5 PCs should preserve the sequential activity. The manifold was visualized by plotting the neural activity projected onto the embedding defined by the first two LEM dimensions. In Fig. 2d (right) the neural activity of the entire session was projected onto the embedding.

Both approaches revealed a ring-shaped manifold along which the population activity propagated repeatedly with periodic boundary conditions. One “cycle” of the population oscillation was defined as one full turn of the population activity along the ring-shaped manifold.

### Phase of the oscillation

To track the progression of the population activity over time, we leveraged the low dimensionality of the ring-shaped manifold and the circular nature of the population activity, and parametrized the population activity with a single time-dependent parameter, which we called the “phase of the oscillation”. Hence, the phase of the oscillation varied as a function of time (bin size = 129 ms) and tracked the progression of the neural activity during the population oscillation. The neural activity was projected onto a two-dimensional plane using PCA. The use of PCA avoided the selection of hyperparameters, which is required in all non-linear dimensionality reduction techniques including LEM.

Let *PCi_t_ (t*) be the projection of the neural population activity onto Principal Component *i* (PC*i*). The neural population activity at time point *t* projected onto the plane defined by PC1 and PC2 is then given by (*PC*1*_t_ (t*), *PC*2*_t_ (t*)), which defines a vector in this plane. The phase of the oscillation is defined as the angle of this vector with respect to the PC1 axis and is given by

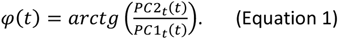

During one cycle of the population oscillation, the phase of the oscillation continuously traversed the range [-π,π), which was consistent with the population activity propagating through the network and describing one turn along the ring-shaped manifold.

### Joint distribution of cross correlation time lag and angular distance in the PCA sorting

To further characterize the sequential activation in the MEC neural population and to introduce a score that would determine the extent to which a session exhibited population oscillations (see “Oscillation score”), we determined the relationship between the time lags that maximized the cross correlation between the calcium activity of two cells (*τ*) and their angular distances in the PCA sorting (*d*). In the plane generated by PC1 and PC2, the loadings of each neuron defined a vector, for which we computed the angle 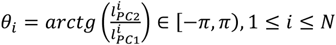, with respect to the axis of PC1, where 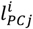 is the loading of cell *i* on *PCj* and *N* is the total number of recorded neurons (see “Correlation and PCA sorting methods”). The angular distance *d* between any two cells in the PCA sorting was calculated as the difference between their angles wrapped in the interval [*−π*, *π*) (see Fig. 2f left),

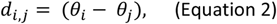

where 1 *≤ i ≤ N*,1 *≤ j ≤ N*. The Matlab function “angdiff” was used for computing this distance. Note that the angular distance maps how far apart two cells are in the raster plot when cells are sorted according to the PCA method.

To estimate the joint distribution of cross correlation time lags and angular distances in the PCA sorting, the cross correlations between all pairs of cells were calculated using a maximum time lag of 248 s. For each cell pair the time lag at which the cross correlation peaked (*τ*) and the angular distance in the PCA sorting (*d*) were calculated. A discrete representation was used for these two variables: in all analyses, and unless stated otherwise, the range of possible *τ* values, i.e. [-248,248] s, was discretized into 96 bins of size 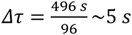 and the range of possible *d* values, i.e. [-*π*, *π*) rad, was discretized into 11 bins of size 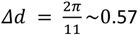 rad. Using those bins, the joint distribution of *τ* and *d* was expressed as a 2D histogram that counted the number of cell pairs observed for every combination of *τ* bins and *d* bins, normalized by the total number of cell pairs.

### Oscillation score

While striking population oscillations were observed in multiple sessions and animals, the population activity exhibited considerable variability, ranging from non-patterned activity to highly stereotypic and periodic sequences (Extended data Fig. 4a). This variability prompted us to quantify, for each session, the extent to which the population activity was oscillatory, which we did by computing an oscillation score. For each session, we first calculated the phase of the oscillation *<p(t*) (bin size = 129 ms, Equation 1), which tracks the progression of the population activity in the presence of population oscillations (see “Phase of the oscillation” and Fig. 2e). Next the PSD of *si n(<p(t*)) was calculated using Welch’s method with Hamming windows of 17.6 min (8192 bins of 129 ms in each window) and 50% of overlap between consecutive windows (“pwelch” Matlab function, see “Autocorrelations and spectral analysis of single cell calcium activity”). If the PSD peaked at 0 Hz and the PSD was strictly decreasing, the phase of the oscillation was not oscillatory and hence the population activity was not periodic in the analysed session. In this case the oscillation score was set to zero. Otherwise, prominent peaks in the PSD at a frequency larger than 0 Hz were identified. In order to disentangle large-amplitude peaks from small fluctuations in the PSD, a peak at frequency *f_max_* was considered prominent and indicative of periodic activity if its amplitude was larger than (i) 9 times the mean of the tail of the PSD (i.e. <PSD(*f*> *f_max_*)>, < > indicates the average over frequencies) and (ii) 9 times the minimum of the PSD between 0 Hz and *f_max_* (i.e. min(PSD(*f* < *f_max_*))). If no peak in the PSD met these criteria the oscillation score was set to zero.

Otherwise, the presence of a prominent peak in the PSD calculated on *si n(<p(t*)) was considered indicative of periodic activity at the population level. Yet a crucial component for observing oscillatory sequences is that cells fire periodically and that the time lag that maximizes the cross correlations between the calcium activity of pairs of cells that are located at a fixed distance in the sequence comes in integer multiples of a minimum time lag, which ensures that cells oscillate at a fixed frequency and that the calcium activity of one cell is temporally shifted with respect to the other. To quantify the extent to which these features were present in the data, we computed the joint distribution of time lags and angular distance in the PCA sorting (*τ* was discretized into 240 bins and *d* was discretized into 11 bins, see “Joint distribution of cross correlation time lag and angular distance in the PCA sorting”). Next for each bin *i* of *d*,1 *≤ i ≤* 11, we calculated the PSD of the distribution of *τ* conditioned on the distance bin *i* (Welch’s methods, Hamming windows of 128 *τ* bins with 50% overlap between consecutive windows, “pwelch” Matlab function). The presence of a peak in this signal indicated that for bin *i* of *d*, the time lag that maximizes the cross correlations between cells was oscillatory (i.e. it peaked at multiples of one specific time lag), as expected when cells are active periodically with an approximately fixed frequency and also with harmonics of the primary frequency. The presence (or absence) of a peak that satisfied the condition of being larger than (i) 10 times the mean of the tail of the PSD (same definition as above), and (ii) 4.5 times larger than the minimum between 0 Hz and the frequency at which the PSD peaked, was identified (same definition as above, the parameters are different from the ones used above because the signals are very different). The oscillation score was then calculated as the fraction of angular distance bins for which a peak was identified.

Based on the bimodal distribution of oscillation scores obtained in the MEC data (Extended data Fig. 4c), a session was considered to express population oscillations if the oscillation score was ≥ 0.72, which was equivalent to asking that at least 8 out of the 11 distributions of *τ* conditioned on bin *i* of *d*,1 *≤ i ≤* 11, had a significant peak in their PSD. This choice of cut-off also accounted for the fact that for distances in the PCA sorting that are close to zero, cells exhibit instantaneous coactivity rather than coactivity shifted by some specific time lag, which makes the conditional probability not oscillatory. After applying the cut-off, 15 of 27 MEC sessions in 5 animals were classified as oscillatory (Extended data Fig. 4c), and among those 15 sessions, 10 were recorded with synchronized behavioural tracking (see “Self-paced running behavior under sensory-minimized conditions”). The number of recorded cells in the oscillatory sessions ranged from 207 to 520. In the rest of the data set, 0 of 25 PaS sessions in 4 animals were classified as oscillatory, 0 of 19 VIS sessions in 3 animals were classified as oscillatory.

### Oscillation bin size

The population oscillation progressed at frequencies < 0.1 Hz that varied from session to session. The “oscillation bin size” was a temporal bin size representative of the time scale of the population oscillation in each session. It was used to quantify single cell and neural population dynamics, for which describing the neural activity at the right time scale was fundamental (e.g. see “Transition probabilities and graph representation”). For each oscillatory session the period of the population oscillation, denoted by *P_osc_*, was calculated as the inverse of the frequency *f_max_* at which the PSD of the signal *sin(<p(t*)) peaked (see Equation 1 and “Oscillation score”), i.e. *P_osc_* = *f_max_^−1^*. Note that this estimate of the period was reliable when during most of the session the network engaged in the population oscillation, in which case the estimate was equivalent to the length of the session divided by the total number of oscillation cycles. However, it became less reliable the more interrupted the population oscillation was.

The oscillation bin size *T_osc_* was computed as the period of the population oscillation divided by 10,

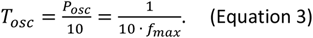

This choice of bin size was made so that each cycle of the population oscillation would progress across ~10 time points. Across 15 oscillatory sessions, the oscillation bin size ranged from 3 to 17 s (see Extended data Fig. 7l).

In sessions without population oscillations, there was not a well-defined peak in the PSD of *sin(<p(t*)), and therefore the oscillation bin size was not possible or meaningful to calculate. Yet, to perform the quantifications of network dynamics at temporal scales similar to the ones investigated in oscillatory sessions, the mean oscillation bin size computed across all oscillatory sessions was used (mean oscillation bin size = 8.5 s).

Unless otherwise indicated, the utilized bin size was 129 ms.

### Identification of individual oscillation cycles

The characterization of the population oscillation required multiple analyses that relied on identifying individual cycles, for example to quantify the length of the cycles and their variability. The procedure for identifying individual cycles was based on finding the time points at which each cycle began (visualized typically at the bottom of the raster plot) and ended (visualized typically at the top of the raster plot, see Extended data Fig. 5a). Note that the beginning and the end of the cycle are arbitrary because of the periodic boundary conditions in the cycle progression, and therefore a different pair of phases that are 2π apart could have been used for defining the beginning and the end of the cycle.

One cycle was defined as one full turn of the phase of the oscillation (see “Phase of the oscillation”), i.e. during one cycle the phase of the oscillation traversed 2*π*. To calculate the phase of the oscillation and determine the time epochs during which it traversed 2*π*, we smoothed the calcium activity of all cells (bin size = 129 ms) using a gaussian kernel of width equal to the oscillation bin size. Next, the phase of the oscillation was calculated and discretized into 10 bins (i.e. the range [*−π*, *π*) was discretized into 10 bins). Time points at which the phase of the oscillation belonged to a bin that was 3 or more bins away from the bin in the previous time point were considered as discontinuity points and were used to define the beginning and the end of putative cycles. Putative cycles were classified as cycles if the phase of the oscillation smoothly traversed the range [*−π*, *π*) rad in an ascending manner. To account for variability, decrements of up to 1 bin of the phase of the oscillation were allowed. Points of sustained activity were disregarded. Segments of cycles in which the phase of the oscillation covered at least 5 bins (i.e. 50% or more of the range [*−π*, *π*) rad) were also identified.

### Cycle length, population oscillation frequency and inter-cycle interval

The length of individual cycles (cycle length) was defined as the amount of time that it takes the phase of the oscillation to cover the range [*−π*, *π*) in a smooth and increasing manner, which is consistent with the population activity completing one full turn along the ring-shaped manifold. To calculate the cycle length, the time interval between the beginning and the end of the cycle was determined (see “Identification of individual oscillation cycles”).

To quantify the variability in cycle length within and between sessions, two approaches were adopted. In approach 1 (Extended data Fig. 5e left), the standard deviation of cycle lengths was computed for each oscillatory session. To estimate significance, in each of 500 iterations all cycles across 15 oscillatory sessions were pooled (421 cycles in total) and randomly assigned to each session while keeping the original number of cycles per session unchanged. For each iteration the standard deviation of the cycle lengths randomly assigned to each session was calculated. In approach 2 (Extended data Fig. 5e right), for each session *i*,1 *≤ i ≤* 1*5*, where 15 is the total number of oscillatory sessions, we considered all pairs of cycles within session *i* (“within session” group) or alternatively all pairs of cycles such that one cycle belongs to session *i* and the other cycle to session *j*, *j ≠ i* (“between session” group). For each cycle pair in each group, the ratio between the shortest cycle length and the longest cycle length was calculated. The mean was computed over pairs of cycles in each group for each session separately. Notice that the larger this ratio the more similar are the cycle lengths.

The frequency of the population oscillation was calculated as the total number of identified individual cycles in a session, divided by the total amount of time the network engaged in the population oscillation during the session, which was computed as the length of the temporal window of concatenated cycles.

The inter-cycle interval was defined as the length of the epoch from the termination of one cycle and the beginning of the next one.

### Mean event rate during segments of the oscillation cycle

To determine how population activity varied during individual cycles of the population oscillation (Extended data Fig. 5c), the following approach was adopted. For each oscillatory session (see “Oscillation score”) all individual cycles were identified (see “Identification of individual oscillation cycles”). Each cycle was divided into 10 segments of equal length. For each cycle segment, the mean event rate was calculated as the total number of calcium events across cells divided by cycle segment duration and number of cells. For each session the mean event rate per segment was calculated over cycles.

### Locking to the phase of the oscillation

To calculate the extent to which individual cells were tuned to the population oscillation, two quantities were used: the locking degree and the mutual information between the calcium event counts and the phase of the oscillation. For each oscillatory session, the phase of the oscillation *<p(t*) was computed (see Equation 1) and individual cycles were identified (see “Identification of individual oscillation cycles”). Next, the time points that corresponded to all individual cycles in one session were concatenated, which generated a new matrix of calcium activity in which the network engaged in the population oscillation uninterruptedly.

The locking degree was computed for each cell as the mean resultant vector length over the phases of the population oscillation at which the calcium events occurred (bin size = 129 ms, function “circ_r” from the Circular Statistics Toolbox for Matlab^106^). The locking degree has a lower bound of 0 and upper bound of 1. It is equal to 1 if all oscillation phases at which the calcium events occurred are the same, i.e. perfect locking, and equal to zero if all phases at which the calcium events occurred are evenly distributed (total absence of locking). To estimate significance, for each cell a null distribution of locking degrees was built by temporally shuffling the calcium activity of that cell 1000 times while the phase of the oscillation remained unchanged, and by computing, for each shuffle realization, the locking degree (shuffling was performed as in “Sorting of temporally shuffled data”). The 99^th^ percentile of the estimated null distribution was used as a threshold for significance.

In order to assess the robustness of the locking degree, the obtained results were compared with a second measure based on information theory^107^: the mutual information between the counts of calcium events (“event counts”) and the phase of the oscillation (bin size = 0.52 s). To estimate the reduction in uncertainty about the phase of the oscillation (P) given the event counts of the calcium activity (S), Shannon’s mutual information was computed as follows^108^:

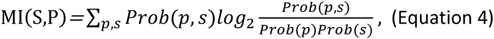

where *Prob(p*, *s*) is the joint probability of observing a phase of the oscillation *p* and an event count *s*, *Prob(s*) is the marginal probability of event counts and *Prob(p*) is the marginal probability of the phase of the oscillation. All probability distributions were estimated from the data using discrete representations of the phase of the oscillation and the event counts. The event counts were partitioned into s_max_+1 bins to account for the absence of event counts as well as all possible event counts, where s_max_ is the maximum number of event counts per cell in a 0.52 s bin, and the phase of the oscillation was discretized into 10 bins of size 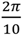.

The mutual information is a non-negative quantity that is equal to zero only when the two variables are independent, i.e. when the joint probability is equal to the product of the marginals *Prob(p*, *s*) = *Prob(p*)*Prob(s*). However, limited sampling can lead to an overestimation in the mutual information in the form of a bias^109^. In order to correct for this bias, the calcium activity was temporally shuffled (as in “Sorting of temporally shuffled data”) and the mutual information between the event counts of the shuffled calcium activity and the phase of the oscillation, which remained unchanged, was calculated. This procedure, which destroyed the pairing between event counts and phase of the oscillation, was repeated 1000 times and the average mutual information across the 1000 iterations was computed and used as an estimation of the bias in the mutual information calculation. In the right panel of Fig. 3a, we report both the mutual information and the bias. In Extended data Fig. 6a, the corrected mutual information was reported (MI_c_), where the bias (〈MI*_sh_*〉*_it_*_e_*_r_*_a*t*_*_ions_*) was subtracted out from the Shannon’s mutual information (MI): MI_c_=MI-〈MI*_sh_*〉*_it_*_e_*_r_*_a*t*_*_ions_*.

Note that the locking degree and the mutual information between the event counts and the phase of the oscillation yielded consistent results (see Fig. 3a and Extended data Fig. 6a).

### Tuning of single cells to the phase of the oscillation

The selectivity of each cell to the phase of the oscillation was visualized through tuning curves and quantified through their preferred phase.

#### Tuning curves

The phase of the oscillation *φ(t*) was estimated (Equation 1) and the range of phases [*−π*, *π*) was partitioned into 40 bins of size 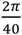 rad. For each cell the tuning curve in the phase bin *j*, *j* = 0, …,39, was calculated as the total number of event counts that occurred at phases within the range 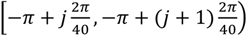 divided by the total number of event counts during the population oscillation.

#### Preferred phases

The preferred phase of each cell was calculated as the circular mean over the oscillation phases at which the calcium events occurred (function “circ_mean” from the Circular Statistics Toolbox for Matlab^106^).

To determine the extent to which the preferred phases across locked cells were uniformly distributed in one recorded session, the distribution of the cells’ preferred phases, that we shall denote *Q*, was estimated by discretizing the preferred phases into 10 bins of size 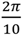 rad. The entropy of this distribution 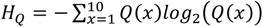 was calculated and used to compute the entropy ratio *H_ratio_*, which quantifies how much *Q* departs from a flat distribution:

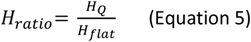

where *H_flat_* is the entropy of a flat distribution using 10 bins, i.e. *H_flat_* = 3.32 bits. The closer *H_ratio_* is to 1 the flatter *Q* is, and therefore all preferred phases tend to be equally represented. The smaller *H_ratio_* is, the more uneven *Q* is and some preferred phases tend to be more represented than others.

To estimate significance, for each session the procedure for calculating *H_ratio_* was repeated for 1000 iterations of a shuffling procedure where the preferred phase of the cells was calculated after the values of the phase of the oscillation were temporally shuffled. In Extended data Fig. 6f, both panels, for each session the 1000 shuffle realizations were averaged.

### Participation index

The Participation Index (PI) quantifies the extent to which a cell’s calcium events were distributed across all cycles of the population oscillation, or rather concentrated in a few cycles. The participation index was calculated for each cell separately as the fraction of cycles needed to account for 90% of the total number of calcium events. To compute the participation, individual cycles were identified (see “Identification of individual oscillation cycles”), and for each cell the number calcium events per cycle was calculated and normalized by the total number of calcium events across all concatenated cycles, which yields the fraction of calcium events per cycle. This quantity was sorted in an ascending manner and its cumulative sum was calculated. The participation index is the minimum fraction of the total number of the cycles for which the cumulative sum of the fraction of calcium events per cycle *≥* 0.9 (results remain unchanged when the cumulative sum is required to be *≥* 0.95).

### Relationship between tuning to the phase of the oscillation and single-cell oscillatory frequency

To determine whether the frequency of oscillation of single cell calcium activity was correlated with the extent to which the cell was locked and participated in the population oscillation, for each cell the ratio between its oscillatory frequency (see “Autocorrelations and spectral analysis of single cell calcium activity”) and the frequency of the population oscillation (see “Cycle length, population oscillation frequency and inter-cycle interval”) was calculated and denoted “relative frequency”. Next, for each session cells were divided into two groups: one group had cells with relative frequency ~1 (cells whose oscillatory frequencies were most similar to the population oscillation frequency), and the other group had cells with relative frequency *≠*1 (cells whose oscillatory frequencies were most different from the population oscillation frequency). The size of each group was the same and was given by a percentage α of the total number of recorded cells in a session. For each group the locking degree (see “Locking to the phase of the oscillation”) and the participation index (see “Participation index”) were compared. For the quantification across all 15 oscillatory sessions, the mean locking degree and participation index were calculated for each group separately and for each session separately, and all 15 sessions were pooled. α was varied from 5% to 50%.

### Anatomical distribution of preferred phases and participation indexes

To determine whether the entorhinal population oscillation resembled travelling waves, during which neural population activity moves progressively across anatomical space^43–48^, the distributions of anatomical pairwise distances for cells with similar and different (1) preferred phase and (2) participation index were computed. To perform these quantifications the first step was to calculate, for each session, the anatomical distance between all pairs of cells. To calculate those pairwise distances we used the centroid of each cell in the FOV (Suite2P^39^).

#### Preferred phase

Because the progression of the neural population activity during the population oscillation can be tracked by the phase of the oscillation (Fig. 2e), we determined whether there is topography in the cells’ preferred phases. The preferred phase of all cells in one session were computed (see “Tuning of single cells to the phase of the oscillation”) and cells were divided into two groups, one of preferred phases ~0 rad, and one of preferred phases ~π rad (Extended data Fig. 6i). The size of each group was the same and was given by a percentage α of the total number of locked cells in a session (see “Locking to the phase of the oscillation”). All cells in each group were locked to the phase of the oscillation. α was varied from 5% to 50%. Pairwise anatomical distances between the cells with preferred phase ~0 rad were calculated and assigned to the group “similar”. Pairwise anatomical distances in the “different” group were determined such that one cell of each pair had a preferred phase ~0 rad and the other cell a preferred phase ~*π* rad. A comparison of the two groups of pairwise distances is shown for one example session in Fig. 3f left. For quantification across all 15 oscillatory sessions, in Fig. 3f right, the means for the two groups, similar and different, were computed for each session separately. Notice that there were no significant differences in the pairwise anatomical distances between cells with similar and different preferred phases regardless of the value of α (Extended data Fig. 6j).

#### Participation index

Given that several properties of MEC cells follow a dorsoventral or mediolateral organization^2,3,42^ we determined whether there is topography in the neurons’ participation in the oscillation cycles (see “Participation index”). The same procedure as described for the preferred phases was followed. Cells were divided into two groups. The size of each group was the same and was given by a percentage α of the total number of locked cells in a session. One group comprises the cells with the lowest participation indexes, and the other group the cells with the highest participation indexes (Extended data Fig. 6k). Pairwise anatomical distance between all cell pairs in the low participation index group were calculated and assigned to the group “similar”. Pairwise anatomical distances for the “different” group were determined for all pairs of cells such that one cell of the pair belonged to the low participation index group, and the other cell to the high participation index group. Notice that there were no significant differences in pairwise anatomical distances between cells with similar and different PIs regardless of the value of α (Extended data Fig. 6l).

### Procedure for merging steps

In order to average out the variability observed in single cells at the level of oscillatory frequency, locking degree and participation index while preserving the temporal properties of the population oscillation, an iterative process that defines new variables from combining the calcium activity of cells in small neighborhoods was implemented for each session separately (Extended data Fig. 7a). This process is similar to a coarse-graining approach^110^.

First, the *N* recorded cells in one session were sorted according to the PCA method. In the first iteration of the procedure, named merging step one, the calcium activity (see “Binary deconvolved calcium activity and matrix of calcium activity”) of pairs of cells that were positioned next to each other in the PCA sorting were added up (merging step 1 in Extended data Fig. 7a). This resulted in 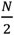 new variables, which in merging step 2 were grouped together in pairs of adjacent variables by adding up their activity, which yielded 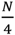 new variables. Note that because in the PCA sorting cells whose activity is synchronous are positioned adjacent to each other, the new variables consist of groups of co-active cells.

In general, merging step *j* generates 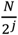 variables by adding up the activity of pairs of 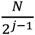 variables from merging step *j −* 1, *j* > 1, with each new variable defined as:

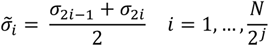

where 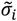 is the *i^th^* new variable that results from adding up *C_2i-1_* and *C_2i_*, which were computed in the previous merging step, *j −* 1. In merging step 1, *C_2i-1_* and *C_2i_* are the calcium activity of cells in the position 2*i −* 1 and 2*i*, 1 *≤ i ≤ N*, in the sorting obtained with the PCA method.

This procedure was repeated 6 times until *~*10 variables were obtained in each session (the exact number of variables depended on the number of recorded cells, *N*, in each session). If *N* was an odd number, the last cell in the sorting obtained with the PCA method was discarded and the procedure was applied to the first *N −* 1 cells in the sorting. In every merging step the participation index (see “Participation index”) of each new variable was calculated (see Extended data Fig. 7b).

### Division of cells into ensembles

After 5 merging steps (and for approximately 10 variables), the participation index reached a plateau (Extended data Fig. 7b). This motivated the decision to split the recorded cells into 10 variables, which we later used to quantify the network dynamics (see “Analysis of network dynamics using ensembles of co-active cells”). From now on we will refer to those variables as “ensembles”, to highlight the fact that cells in each ensemble are co-active. The same number of ensembles was used in sessions that did not exhibit population oscillations.

To distribute cells into 10 ensembles, cells were sorted according to the PCA method. If 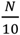 is an integer, where *N* is the total number of cells in one session, then each ensemble contains 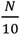 cells and the set of cells that belong to ensemble *i*,1 *≤ i ≤* 10, is 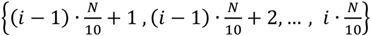. If 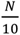 is *not* an integer then ensembles 1 to 9 contain 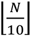 cells and ensemble 10 contains 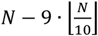 cells, where 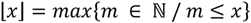 and 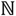 is the set of natural numbers. In this case the set of cells that belongs to each ensemble is:

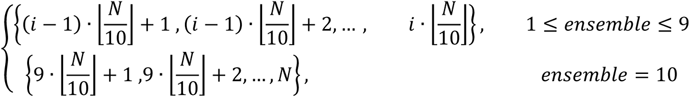

Note that each cell was assigned to only one ensemble.

After each cell was assigned to one of the 10 ensembles, the activity of each ensemble as a function of time was calculated as the mean calcium activity across cells in that ensemble.

Finally, to calculate the oscillation frequency of ensemble activity, the PSD was calculated (Welch’s methods, 8.8 min Hamming window with 50% overlap between consecutive windows, “pwelch” Matlab function). The oscillation frequency was estimated as the frequency at which the PSD peaked. For each session, the oscillation frequency of the activity of the ensembles was compared to the frequency of the population oscillation, which was computed as the total number of cycles in the session divided by the amount of time the network engaged in the population oscillation. The latter was calculated as the length of the temporal window of concatenated cycles (see “Identification of individual oscillation cycles”).

### Tuning of single-cell activity to ensemble activity

To quantify the degree of tuning of a cell’s calcium activity to the ensemble activity, and hence determine whether ensemble activity was representative of single cell calcium activity, we calculated, for all cells in a recorded session, the Pearson correlation between the calcium activity and the ensemble activity. Cells were divided into 10 ensembles (see “Division of cells into ensembles”) and the activity of each ensemble as a function of time was calculated as the mean calcium activity across cells in the ensemble (bin size = 129 ms). For each neuron *i*, 1 *≤ i ≤ N*, where *N* is number of recorded cells in the session, the Pearson correlation *P_i_*_,*j*_ between the neuron’s calcium activity and the activity of ensemble *j*, 1 *≤ j ≤* 10, was calculated for each ensemble separately. When calculating this set of 10 correlations, the activity of the cell for which the tuning is being computed was excluded in the calculation of the ensemble activity (note that by construction each cell is assigned to only one ensemble). Next, for each cell *i*, the most representative ensemble was calculated as the one for which the Pearson correlation was maximal, i.e.,

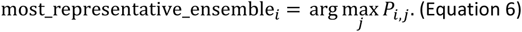

In order to determine whether the activity of the ensemble a cell was assigned to (see “Division of cells into ensembles”) was the most representative of the single cell calcium activity, we quantified how similar the most representative ensemble and the ensemble assigned based on the PCA sorting were, expecting that the most representative ensemble and the assigned ensemble would coincide. For each cell the distance between these was computed subject to periodic boundary conditions in the ensembles (for example, the distance between ensemble one and ten was one and not nine).

For each session the fraction of cells that displayed specific distances between their assigned ensemble based on the PCA sorting and their most representative ensemble was calculated for the entire range of distances and presented as a probability. Probabilities were next averaged across sessions (Extended data Fig. 7f). To estimate significance, for each cell in a session the procedure for identifying the most representative ensemble was repeated in 500 iterations of a shuffle realization where the ensemble activity remained fixed but the calcium activity was temporally shuffled (as in “Sorting of temporally shuffled data”). For each of the 500 shuffle realizations per session the probabilities of observing specific distances between the PCA-assigned and the most representative ensemble were calculated and averaged, yielding the mean shuffled probability per session. These probabilities (15 in total for the 15 oscillatory sessions) were then pooled and compared to the recorded data.

The probability that the assigned ensemble based on the PCA method and the most representative ensemble coincide was large for MEC (Extended data Fig. 7f), intermediate for VIS and low for PaS (Extended data Fig. 10c).

### Anatomical distribution of ensembles

Analyses performed on the preferred phases and participation indexes of single cells indicated that the population oscillation is not topographically organized, and hence it is not a travelling wave (see “Anatomical distribution of preferred phases and participation indexes” and Fig. 3e-h). To determine whether this result was upheld when cells were sorted in ensembles of co-active neurons, the centroid of each cell in the FOV (provided by Suite2P^39^) was used to calculate the anatomical distance between all pairs of cells in a session. Next, for each session the pairwise anatomical distances were divided into two groups: the “within ensemble” group and the “across ensemble” group. In the former, only pairwise anatomical distances between cells that were assigned to the same ensemble were considered (see “Division of cells into ensembles”). In the latter, we considered pairwise anatomical distances between cells of different ensembles, such that one cell of the pair was assigned to ensemble *i*, *i* = 1, …, 10, and the other to ensemble *j*, *j* = 1, …, 10, *i ≠ j*. This was done for each session and each ensemble separately. In Extended data Fig. 7j, for each ensemble of the example session shown in Fig. 2a, the “within ensemble” group was compared to the “across ensemble” group. Next, for the example session, the data in both groups was pooled and the two groups were compared in the left panel of Extended data Fig. 7k. For the quantification across all 15 oscillatory sessions in the right panel of Extended data Fig. 7k (including the session in Extended data Fig. 7j), the means in both groups were calculated for each session separately.

### Analysis of network dynamics using ensembles of co-active cells

We adopted an ensemble approach to quantify the network dynamics (see “Procedure for merging steps” and “Division of cells into ensembles”). With a total of 10 ensembles this approach averaged out the variability observed in single-cell oscillation frequency, locking degree and participation index while keeping the temporal progression of the oscillatory sequences (Extended data Fig. 7n). In sessions with population oscillations, all individual cycles were identified (see “Identification of individual oscillation cycles”) and the corresponding time bins were concatenated, which yielded a new matrix of calcium activity in which the population oscillation was uninterrupted. Next, cells were divided into ensembles (see “Division of cells into ensembles”) and ensemble activity was downsampled using as bin size the oscillation bin size of the session (see “Oscillation bin size”). This procedure yielded a matrix, the “ensemble matrix”, with the activity of each ensemble corresponding to a single row (10 rows in total), and as many columns as time points when sampled at the oscillation bin size. In non-oscillatory sessions, the full matrix of calcium activity was used and the temporal downsampling was conducted at the mean oscillation bin size computed across all 15 oscillatory sessions; i.e. bin size = 8.5 s (see “Oscillation bin size calculation” for a description of the bin size used in non-oscillatory sessions). For both types of sessions (with and without oscillations), the activity of the 10 ensembles was described through a vector expressing, at each time point, the ensemble number with the highest activity at that time point (see Extended data Fig. 7m,n). This vector was used to perform the following analyses (i-iii).

#### (i)#Transition probabilities and graph representation

The transition probability from ensemble *i* to ensemble *j* was quantified as the number of times the transition *i → j* was observed in the data of one session, normalized by the total number of transitions in one session. Transitions were identified from the vector that contained the ensemble number with maximum activity at each time point (transitions to the same ensemble between consecutive time points were disregarded). Transitions were allocated in a matrix of transition probabilities *T* of size 10×10, since 10 ensembles were used. In this matrix, the component *(i*, *j*) expressed the transition probability from ensemble *i* to ensemble *j*.

To establish statistical significance of the transition probabilities, the data was shuffled 500 times. In each shuffle realization, each row of the matrix of calcium activity (with concatenated cycles in the case of oscillatory sessions) was temporally shuffled (as in “Sorting of temporally shuffled data”), and the procedure for calculating the ensemble matrix and transition probabilities was applied to the shuffled data. For each transition *i → j* the 95^th^ percentile of the null distribution was used to define a cut-off.

The matrices of transition probabilities obtained from the recorded data and from the shuffle realizations were used as adjacency matrices to create graphs. In the graph representation each node represents one ensemble, each edge indicates the transition probability between two nodes, the thickness of the edge is proportional to the transition probability, and the arrow indicates the transition direction. In Extended data Fig. 7p and Extended data Fig. 10d, the edges in red indicate that the corresponding transition probabilities were larger than the cut-off for significance.

#### (ii)#Probability of sequential activation of ensembles

To determine whether preferences in ensemble transitions gave rise to sequences of ensemble activity, we calculated the probability of sequential ensemble activation according to the following procedure. From the vector expressing the ensemble number with the highest activity at each time point (sampled at the oscillation bin size), strictly increasing sequences of all possible lengths (from 2 to 10 ensembles) were identified. The number of ensembles in each sequence was the number of ensembles that were active in consecutive time points (epochs of sustained activity were disregarded). While the sequences had to be strictly increasing, they did not have to be continuous. Sequences could skip ensembles, in which case the maximum number of ensembles in one sequence was less than 10. The probability of the sequential activation of *k* ensembles, *k* = 2, …,10, was next estimated as the number of times a sequence of *k* ensembles was found, normalized by the total number of identified sequences. Note that all subsequences were also included in this estimation. For example, if the ensembles 1, 2 and 3 were active in consecutive time points, a sequence of three ensembles was identified, as well as three subsequences of two ensembles each: 1,2, as well as 2,3 and 1,3.

In order to test for significance, the shuffled data from “Transition probabilities and graph representation” was used. The procedure to compute the probability of sequential activation of ensembles was applied to each of the 500 shuffle realizations performed per session. Shuffled data was compared with recorded data.

#### (iii)#Sequence score

The *sequence score* measures how sequential the ensemble activity is. It is calculated from the probability of sequential activation of ensembles as the probability of observing sequences of 3 or more ensembles. The sequence score was calculated for each session of the dataset separately. To determine if the obtained scores were significant, for each session the 500 shuffle realizations used in “Probability of sequential activation of ensembles” for assessing significance of the probability of sequential activation of ensembles were used to calculate the sequence score on shuffled data. Those values were used to build a null distribution, and the 99^th^ percentile of this distribution was chosen as the threshold for significance.

### Estimation of number of completed laps on the wheel, speed and acceleration

Features of the animal’s behaviour were used to determine whether the MEC population oscillation was modulated by movement.

The wheel had a radius of 8.54 cm (see “Self-paced running behavior under sensory-minimized conditions”) and a perimeter of 53.66 cm. Therefore animals had to run for ~53.7 cm to complete one lap on the wheel. For each session, we estimated the number of completed laps on the wheel from the position on the wheel recorded as a function of time. The number of completed laps during one cycle of the oscillation (see “Identification of individual oscillation cycles”) was calculated as the total distance run during the cycle divided by 53.7 cm.

The speed of the animal was numerically calculated as the first derivative of the position on the wheel as a function of time (the sampling frequency of the position was 40 Hz for mice 60355 (MEC), 60353, 60354 and 60356 (PaS). The sampling frequency was 50 Hz for mice 60584 and 60585 (MEC), 60961, 92227 and 92229 (VIS). For mice 59911, 59914 (MEC) and 59912 (PaS), the wheel tracking was not synchronized to the ongoing image acquisition; see “Self-paced running behavior under sensory-minimized conditions”. The obtained speed signal from the former group of mice was interpolated so that the speed values matched the downsampled imaging time points (sampling frequency = 7.73 Hz), and smoothed using a square kernel of 2 s width. A threshold was applied such that all speed values that were less than 2 cm/s were set to zero and all speed values larger than 2 cm/s remained unchanged. The obtained speed signal was used to define immobility (running) bouts as the set of consecutive time points (bin size=129 ms) for which the speed was equal to (larger than) zero (a similar approach was used in ref. 36). The acceleration was numerically calculated as the first derivative of the speed signal. Notice that in this case no interpolation was needed.

Because the available data did not have enough statistical power, it was not possible to compare the animals’ behaviour, for example in terms of its running speed and acceleration, between periods with and without ongoing population oscillations.

### Estimation of the probability of observing population oscillations

To determine whether the MEC population oscillation was observed during different behavioural states, the probability of observing the population oscillation was calculated conditioned on whether the animal was running or immobile. For each oscillatory session with behavioural tracking synchronized to the imaging data (10 sessions over 3 animals, see “Self-paced running behavior under sensory-minimized conditions” and “Oscillation score”), all individual cycles were identified (see “Identification of individual oscillation cycles”). The subset of time bins that belonged to individual cycles of the oscillation were extracted and labeled as “oscillation”. Next, a second label was assigned to the time bins depending on whether they occurred during running or immobility bouts (bin labelled as “running” and “immobility” respectively, see “Estimation of number of completed laps on the wheel, speed and acceleration”). After applying this procedure, each time bin had two labels, one indicating the running behavior, and one indicating the presence (or absence) of population oscillation. To estimate the probability of observing the population oscillation conditioned on the animal’s running behavior, all bins labelled as running or immobility were identified and from each subset, the fraction of bins labelled as oscillation was calculated. These probabilities were computed for each session separately.

### Sequences during immobility bouts of different lengths

The population oscillation occurred both during running and immobility bouts. To quantify the extent to which individual cycles progressed during different lengths of immobility bouts, the following procedure was adopted. First, for each session, all immobility bouts were identified and assigned to bins of different lengths (see “Estimation of number of completed laps on the wheel, speed and acceleration”; length bins = 0-3s, 3-5s, 5-10s, 10-15s, 15-20s, >25 s). Second, all individual oscillation cycles were identified (see “Identification of individual oscillation cycles”). Third, for each session and each length bin, the fraction of immobility bouts that were fully occupied by continued cycles was calculated. To estimate significance, for each session the time bins that belonged to all individual cycles were temporally shuffled. The third step of the procedure described above was performed for 500 shuffle iterations per session. In Fig. 5c, the recorded data has 10 data points per length bin, and the shuffled data has 5000 data points per length bin, since 500 shuffled realizations per session were pooled.

### Analysis of speed and cycle onset

To determine whether the onset of the MEC population oscillation cycles was modulated by the animaĺs running speed, changes in speed before and after cycle onset were investigated. For each session all individual cycles were identified (see “Identification of individual oscillation cycles”) and for each cycle the mean speed over windows of 10 s before and after cycle onset was calculated. Because no differences in the mean speed were observed before and after onset (Extended data Fig. 2d left panel), we next determined whether changes in speed were correlated with the onset of oscillation epochs, which were defined as epochs with uninterrupted oscillations, i.e. epochs with successive cycles. The same analysis described above was repeated but only for the subset of cycles that were 10 s or more apart, i.e. for cycles that belonged to different oscillation epochs.

### Manifold visualization for example session in VIS and PaS

To visualize whether the topology of the manifold underlying the population activity in example sessions recorded in VIS and PaS was also a ring, PCA was used and a similar procedure to the one described in “Manifold visualization for MEC sessions” was adopted.

For each example session, one corresponding to VIS and one corresponding to PaS (Fig. 6a-d), PCA was applied to the matrix of calcium activity, which first had each row convolved with a gaussian kernel of width equal to four times 8.5 s, which is the mean oscillation bin size computed across oscillatory sessions (see “Oscillation bin size”). Neural activity was projected onto the embedding generated by PC1 and PC2. Extended data Fig. 9b shows the absence of a ring-shaped manifold in VIS and PaS example sessions.

### Coactivity and synchronization in PaS and VIS sessions

Sessions recorded in PaS and VIS did not exhibit population oscillations. To further characterize their population activity, synchronization and neural co-activity were calculated.

#### Synchronization

Neural synchronization was calculated as the absolute value of the Pearson correlation between the calcium activity of pairs of cells (bin size = 129 ms). For each session, the Pearson correlation was calculated for all pairs of calcium activity (correlations with the same calcium activity were not considered) and used to build a distribution of synchronization values. In Extended data Fig. 10j, these distributions were averaged across sessions for each brain area separately.

#### Co-activity

For each time bin in a session (bin size = 129 ms) the co-activity was calculated as the number of cells that had simultaneous calcium events divided by the total number of recorded cells in the session. This number represented the fraction of cells that was active in individual time bins. Using all time bins of the session, a distribution of co-activity values was calculated. In Extended data Fig. 10k, the distributions were averaged across sessions for each brain area separately.

### Data analysis and statistical analysis

Data analyses were performed with custom-written scripts in Python and Matlab (R2021b). Results were expressed as the mean ± SEM unless indicated otherwise. Statistical analysis was performed using MATLAB and p-values are indicated in the figure legends and figures (n.s.: *p*>0.05, \**p* <0.05, ** *p* <0.01, *** *p* <0.001). Student t-tests were used for paired and unpaired data. For data that displayed no Gaussian distribution and that was unpaired, the Wilcoxon rank-sum test was used. For paired data or one-sampled data, the Wilcoxon signed-rank test was used. Two-tailed tests were used unless otherwise indicated. Correlations were determined using Pearson or Spearman correlations. Friedman tests were used for analyses between groups. No statistical methods were used to predetermine sample sizes but our sample sizes were similar to those reported in previous publications from the lab and in other publications in the field.

### Code availability

Code for reproducing the analyses in this article will be available after publication at figEDhare and/or GitHub.

### Data availability

The datasets generated during the current study will be available after publication at figEDhare.

## References

1. Buzsáki, G., & Moser, E. I. (2013). Memory, navigation and theta rhythm in the hippocampal-entorhinal system. Nature Neuroscience, 16(2), 130–138.

2. Hafting, T., Fyhn, M., Molden, S., Moser, M. B., & Moser, E. I. (2005). Microstructure of a spatial map in the entorhinal cortex. Nature, 436(7052), 801–806.

3. Sargolini, F., Fyhn, M., Hafting, T., McNaughton, B. L., Witter, M. P., Moser, M. B., & Moser, E. I. (2006). Conjunctive representation of position, direction, and velocity in entorhinal cortex. Science, 312(5774), 758–762.

4. Savelli, F., Yoganarasimha, D., & Knierim, J. J. (2008). Influence of boundary removal on the spatial representations of the medial entorhinal cortex. Hippocampus, 18(12), 1270–1282.

5. Solstad, T., Boccara, C. N., Kropff, E., Moser, M. B., & Moser, E. I. (2008). Representation of geometric borders in the entorhinal cortex. Science, 322(5909), 1865–1868.

6. Høydal, Ø. A., Skytøen, E. R., Andersson, S. O., Moser, M. B., & Moser, E. I. (2019). Object-vector coding in the medial entorhinal cortex. Nature, 568(7752), 400–404.

7. Singer, W. (1993). Synchronization of cortical activity and its putative role in information processing and learning. Annual Review of Physiology, 55(1), 349–374.

8. Laurent, G. (1996). Dynamical representation of odors by oscillating and evolving neural assemblies. Trends in Neurosciences, 19(11), 489–496.

9. Buzsaki, G. (2006). Rhythms of the Brain. Oxford University Press.

10. von der Malsburg, C. E., Phillps, W. A., & Singer, W. E. (2010). Dynamic Coordination in the Brain: From Neurons to Mind. MIT Press.

11. Mitchell, S. J., & Ranck Jr, J. B. (1980). Generation of theta rhythm in medial entorhinal cortex of freely moving rats. Brain Research, 189(1), 49–66.

12. Chrobak, J. J., & Buzsáki, G. (1998). Gamma oscillations in the entorhinal cortex of the freely behaving rat. Journal of Neuroscience, 18(1), 388–398.

13. Colgin, L. L. (2016). Rhythms of the hippocampal network. Nature Reviews Neuroscience, 17(4), 239–249.

14. Rabinovich, M., Huerta, R., & Laurent, G. (2008). Transient dynamics for neural processing. Science, 321(5885), 48–50.

15. Vyas, S., Golub, M. D., Sussillo, D., & Shenoy, K. V. (2020). Computation through neural population dynamics. Annual Review of Neuroscience, 43, 249–275.

16. György Buzsáki, M. D. (2019). The Brain from Inside Out. Oxford University Press.

17. Petsche, H., Stumpf, C., & Gogolak, G. (1962). The significance of the rabbit’s septum as a relay station between the midbrain and the hippocampus I. The control of hippocampus arousal activity by the septum cells. Electroencephalography and Clinical Neurophysiology, 14(2), 202–211.

18. Gray, C. M., König, P., Engel, A. K., & Singer, W. (1989). Oscillatory responses in cat visual cortex exhibit inter-columnar synchronization which reflects global stimulus properties. Nature, 338(6213), 334–337.

19. Laurent, G., Wehr, M., & Davidowitz, H. (1996). Temporal representations of odors in an olfactory network. Journal of Neuroscience, 16(12), 3837–3847.

20. Harris, K. D., Csicsvari, J., Hirase, H., Dragoi, G., & Buzsáki, G. (2003). Organization of cell assemblies in the hippocampus. Nature, 424(6948), 552–556.

21. Albrecht, D., Royl, G., & Kaneoke, Y. (1998). Very slow oscillatory activities in lateral geniculate neurons of freely moving and anesthetized rats. Neuroscience Research, 32(3), 209–220.

22. Penttonen, M., Nurminen, N., Miettinen, R., Sirviö, J., Henze, D. A., Csicsvári, J., & Buzsáki, G. (1999). Ultra-slow oscillation (0.025 Hz) triggers hippocampal afterdischarges in Wistar rats. Neuroscience, 94(3), 735–743.

23. Ruskin, D. N., Bergstrom, D. A., Kaneoke, Y., Patel, B. N., Twery, M. J., & Walters, J. R. (1999). Multisecond oscillations in firing rate in the basal ganglia: robust modulation by dopamine receptor activation and anesthesia. Journal of Neurophysiology, 81(5), 2046–2055.

24. Allers, K. A., Kreiss, D. S., & Walters, J. R. (2000). Multisecond oscillations in the subthalamic nucleus: effects of apomorphine and dopamine cell lesion. Synapse, 38(1), 38–50.

25. Allers, K. A., Ruskin, D. N., Bergstrom, D. A., Freeman, L. E., Ghazi, L. J., Tierney, P. L., & Walters, J. R. (2002). Multisecond periodicities in basal ganglia firing rates correlate with theta bursts in transcortical and hippocampal EEG. Journal of Neurophysiology, 87(2), 1118–1122.

26. Wichmann, T., Kliem, M. A., & Soares, J. (2002). Slow oscillatory discharge in the primate basal ganglia. Journal of Neurophysiology, 87(2), 1145–1148.

27. Aladjalova, N. A. (1957). Infra-slow rhythmic oscillations of the steady potential of the cerebral cortex. Nature, 179(4567), 957–959.

28. Leopold, D. A., Murayama, Y., & Logothetis, N. K. (2003). Very slow activity fluctuations in monkey visual cortex: implications for functional brain imaging. Cerebral Cortex, 13(4), 422–433.

29. Lecci, S., Fernandez, L. M., Weber, F. D., Cardis, R., Chatton, J. Y., Born, J., & Lüthi, A. (2017). Coordinated infraslow neural and cardiac oscillations mark fragility and offline periods in mammalian sleep. Science Advances, 3(2), e1602026.

30. Squire, L. R., Stark, C. E., & Clark, R. E. (2004). The medial temporal lobe. Annual Review of Neuroscience, 27, 279–306.

31. Hasselmo, M. E. (2011). How we remember: Brain Mechanisms of Episodic Memory. MIT press.

32. Alonso, A., & Garcia-Austt, E. (1987). Neuronal sources of theta rhythm in the entorhinal cortex of the rat. Experimental Brain Research, 67(3), 493–501.

33. Dombeck, D. A., Khabbaz, A. N., Collman, F., Adelman, T. L., & Tank, D. W. (2007). Imaging large-scale neural activity with cellular resolution in awake, mobile mice. Neuron, 56(1), 43–57.

34. Heys, J. G., Rangarajan, K. V., & Dombeck, D. A. (2014). The functional micro-organization of grid cells revealed by cellular-resolution imaging. Neuron, 84(5), 1079–1090.

35. Gu Y, Lewallen S, Kinkhabwala AA, Domnisoru C, Yoon K, Gauthier JL, Fiete IR, Tank DW (2018). A map-like micro-organization of grid cells in the medial entorhinal cortex. Cell, 175(3), 736–750.

36. Villette, V., Malvache, A., Tressard, T., Dupuy, N., & Cossart, R. (2015). Internally recurring hippocampal sequences as a population template of spatiotemporal information. Neuron, 88(2), 357–366.

37. Carrillo-Reid, L., Miller, J. E. K., Hamm, J. P., Jackson, J., & Yuste, R. (2015). Endogenous sequential cortical activity evoked by visual stimuli. Journal of Neuroscience, 35(23), 8813–8828.

38. Friedrich, J., Zhou, P., & Paninski, L. (2017). Fast online deconvolution of calcium imaging data. PLoS Computational Biology, 13(3), e1005423.

39. Pachitariu, M., Stringer, C., Dipoppa, M., Schröder, S., Rossi, L. F., Dalgleish, H., … & Harris, K. D. (2017). Suite2p: beyond 10,000 neurons with standard two-photon microscopy. BioRxiv doi: https://doi.org/10.1101/061507.

40. Cunningham, J. P., & Byron, M. Y. (2014). Dimensionality reduction for large-scale neural recordings. Nature Neuroscience, 17(11), 1500–1509.

41. Rowland, D. C., Obenhaus, H. A., Skytøen, E. R., Zhang, Q., Kentros, C. G., Moser, E. I., & Moser, M. B. (2018). Functional properties of stellate cells in medial entorhinal cortex layer II. Elife, 7, e36664.

42. Obenhaus, H. A., Zong, W., Jacobsen, R. I., Rose, T., Donato, F., Chen, L., … & Moser, E. I. (2022). Functional network topography of the medial entorhinal cortex. Proceedings of the National Academy of Sciences, 119(7), e2121655119.

43. Meister, M., Wong, R. O., Baylor, D. A., & Shatz, C. J. (1991). Synchronous bursts of action potentials in ganglion cells of the developing mammalian retina. Science, 252(5008), 939–943.

44. Wong, R. O., Meister, M., & Shatz, C. J. (1993). Transient period of correlated bursting activity during development of the mammalian retina. Neuron, 11(5), 923–938.

45. Garaschuk, O., Linn, J., Eilers, J., & Konnerth, A. (2000). Large-scale oscillatory calcium waves in the immature cortex. Nature Neuroscience, 3(5), 452–459.

46. Adelsberger, H., Garaschuk, O., & Konnerth, A. (2005). Cortical calcium waves in resting newborn mice. Nature Neuroscience, 8(8), 988–990.

47. Ackman, J. B., Burbridge, T. J., & Crair, M. C. (2012). Retinal waves coordinate patterned activity throughout the developing visual system. Nature, 490(7419), 219–225.

48. Muller, L., Chavane, F., Reynolds, J., & Sejnowski, T. J. (2018). Cortical travelling waves: mechanisms and computational principles. Nature Reviews Neuroscience, 19(5), 255–268.

49. Ahmed, O. J., & Mehta, M. R. (2012). Running speed alters the frequency of hippocampal gamma oscillations. Journal of Neuroscience, 32(21), 7373–7383.

50. Kropff, E., Carmichael, J. E., Moser, E. I., & Moser, M. B. (2021). Frequency of theta rhythm is controlled by acceleration, but not speed, in running rats. Neuron, 109(6), 1029–1039.

51. Boccara, C. N., Sargolini, F., Thoresen, V. H., Solstad, T., Witter, M. P., Moser, E. I., & Moser, M. B. (2010). Grid cells in pre-and parasubiculum. Nature Neuroscience, 13(8), 987–994.

52. Stringer, C., Pachitariu, M., Steinmetz, N., Carandini, M., & Harris, K. D. (2019). High-dimensional geometry of population responses in visual cortex. Nature, 571(7765), 361–365.

53. Yoon, K., Buice, M. A., Barry, C., Hayman, R., Burgess, N., & Fiete, I. R. (2013). Specific evidence of low-dimensional continuous attractor dynamics in grid cells. Nature Neuroscience, 16(8), 1077–1084.

54. Gardner, R. J., Hermansen, E., Pachitariu, M., Burak, Y., Baas, N. A., Dunn, B. A., Moser, M.-B. & Moser, E. I. (2022). Toroidal topology of population activity in grid cells. Nature, 602(7895), 123–128.

55. Miller, J. E. K., Ayzenshtat, I., Carrillo-Reid, L., & Yuste, R. (2014). Visual stimuli recruit intrinsically generated cortical ensembles. Proceedings of the National Academy of Sciences, 111(38), E4053–E4061.

56. Buzsáki, G., & Vanderwolf, C. H. (1983). Cellular bases of hippocampal EEG in the behaving rat. Brain Research Reviews, 6(2), 139–171.

57. Csicsvari, J., Hirase, H., Czurkó, A., Mamiya, A., & Buzsáki, G. (1999). Oscillatory coupling of hippocampal pyramidal cells and interneurons in the behaving rat. Journal of Neuroscience, 19(1), 274–287.

58. Csicsvari, J., Jamieson, B., Wise, K. D., & Buzsáki, G. (2003). Mechanisms of gamma oscillations in the hippocampus of the behaving rat. Neuron, 37(2), 311–322.

59. Novak, P., Lepicovska, V., & Dostalek, C. (1992). Periodic amplitude modulation of EEG. Neuroscience Letters, 136(2), 213–215.

60. Vanhatalo, S., Palva, J. M., Holmes, M. D., Miller, J. W., Voipio, J., & Kaila, K. (2004). Infraslow oscillations modulate excitability and interictal epileptic activity in the human cortex during sleep. Proceedings of the National Academy of Sciences, 101(14), 5053–5057.

61. Nir, Y., Mukamel, R., Dinstein, I., Privman, E., Harel, M., Fisch, L., … & Malach, R. (2008). Interhemispheric correlations of slow spontaneous neuronal fluctuations revealed in human sensory cortex. Nature Neuroscience, 11(9), 1100–1108.

62. Watson, B. O. (2018). Cognitive and physiologic impacts of the infraslow oscillation. Frontiers in Systems Neuroscience, 44.

63. Pastalkova, E., Itskov, V., Amarasingham, A., & Buzsaki, G. (2008). Internally generated cell assembly sequences in the rat hippocampus. Science, 321(5894), 1322–1327.

64. Malvache, A., Reichinnek, S., Villette, V., Haimerl, C., & Cossart, R. (2016). Awake hippocampal reactivations project onto orthogonal neuronal assemblies. Science, 353(6305), 1280–1283.

65. Fyhn, M., Hafting, T., Treves, A., Moser, M. B., & Moser, E. I. (2007). Hippocampal remapping and grid realignment in entorhinal cortex. Nature, 446(7132), 190–194.

66. McNaughton, B. L., & Morris, R. G. (1987). Hippocampal synaptic enhancement and information storage within a distributed memory system. Trends in Neurosciences, 10(10), 408–415.

67. Leutgeb, S., Leutgeb, J. K., Treves, A., Moser, M. B., & Moser, E. I. (2004). Distinct ensemble codes in hippocampal areas CA3 and CA1. Science, 305(5688), 1295–1298.

68. Alme, C. B., Miao, C., Jezek, K., Treves, A., Moser, E. I., & Moser, M. B. (2014). Place cells in the hippocampus: eleven maps for eleven rooms. Proceedings of the National Academy of Sciences, 111(52), 18428–18435.

69. Buzsáki, G., & Tingley, D. (2018). Space and time: the hippocampus as a sequence generator. Trends in Cognitive Sciences, 22(10), 853–869.

70. Lubenov, E. V., & Siapas, A. G. (2009). Hippocampal theta oscillations are travelling waves. Nature, 459(7246), 534–539.

71. Patel, J., Fujisawa, S., Berényi, A., Royer, S., & Buzsáki, G. (2012). Traveling theta waves along the entire septotemporal axis of the hippocampus. Neuron, 75(3), 410–417.

72. Dragoi, G., & Tonegawa, S. (2011). Preplay of future place cell sequences by hippocampal cellular assemblies. Nature, 469(7330), 397–401.

73. Bittner, K. C., Milstein, A. D., Grienberger, C., Romani, S., & Magee, J. C. (2017). Behavioral time scale synaptic plasticity underlies CA1 place fields. Science, 357(6355), 1033–1036.

74. Ben-Yishai, R., Bar-Or, R. L., & Sompolinsky, H. (1995). Theory of orientation tuning in visual cortex. Proceedings of the National Academy of Sciences, 92(9), 3844–3848.

75. Skaggs, W. E., Knierim, J. J., Kudrimoti, H. S. & McNaughton, B. L. A model of the neural basis of the rat’s sense of direction. Adv. Neural Inf. Process. Syst. 7, 173–180 (1995).

76. Zhang, K. (1996). Representation of spatial orientation by the intrinsic dynamics of the head-direction cell ensemble: a theory. Journal of Neuroscience, 16(6), 2112–2126.

77. Fiete, I. R., Senn, W., Wang, C. Z., & Hahnloser, R. H. (2010). Spike-time-dependent plasticity and heterosynaptic competition organize networks to produce long scale-free sequences of neural activity. Neuron, 65(4), 563–576.

78. Rajan, K., Harvey, C. D., & Tank, D. W. (2016). Recurrent network models of sequence generation and memory. Neuron, 90(1), 128–142.

79. Hebb DO (1949) The organization of behaviour. New York: Wiley.

80. Abeles, M. (1991). Corticonics: Neural Circuits of the Cerebral Cortex. Cambridge University Press.

81. Diesmann, M., Gewaltig, M. O., & Aertsen, A. (1999). Stable propagation of synchronous spiking in cortical neural networks. Nature, 402(6761), 529–533.

82. Kumar, A., Rotter, S., & Aertsen, A. (2008). Conditions for propagating synchronous spiking and asynchronous firing rates in a cortical network model. Journal of Neuroscience, 28(20), 5268–5280.

83. Kumar, A., Rotter, S., & Aertsen, A. (2010). Spiking activity propagation in neuronal networks: reconciling different perspectives on neural coding. Nature Reviews Neuroscience, 11(9), 615–627.

84. Litwin-Kumar, A., & Doiron, B. (2012). Slow dynamics and high variability in balanced cortical networks with clustered connections. Nature Neuroscience, 15(11), 1498–1505.

85. Schaub, M. T., Billeh, Y. N., Anastassiou, C. A., Koch, C., & Barahona, M. (2015). Emergence of slow-switching assemblies in structured neuronal networks. PLoS Computational Biology, 11(7), e1004196.

86. Steriade, M. (1997). Synchronized activities of coupled oscillators in the cerebral cortex and thalamus at different levels of vigilance. Cerebral Cortex 7(6), 583–604.

87. Buzsáki, G. (2002). Theta oscillations in the hippocampus. Neuron, 33(3), 325–340.

88. Taube, J. S., Muller, R. U., & Ranck, J. B. (1990). Head-direction cells recorded from the postsubiculum in freely moving rats. I. Description and quantitative analysis. Journal of Neuroscience, 10(2), 420–435.

89. Mosheiff, N., & Burak, Y. (2019). Velocity coupling of grid cell modules enables stable embedding of a low dimensional variable in a high dimensional neural attractor. Elife, 8, e48494.

90. Waaga, T., Agmon, H., Normand, V. A., Nagelhus, A., Gardner, R. J., Moser, M. B., … & Burak, Y. (2022). Grid-cell modules remain coordinated when neural activity is dissociated from external sensory cues. Neuron, S0896-6273(22)00247-1. doi: 10.1016/j.neuron.2022.03.011. Epub ahead of print. PMID: 35385698.

91. Nicola, W., & Clopath, C. (2019). A diversity of interneurons and Hebbian plasticity facilitate rapid compressible learning in the hippocampus. Nature Neuroscience, 22(7), 1168–1181.

92. Luczak, A., Bartho, P., & Harris, K. D. (2013). Gating of sensory input by spontaneous cortical activity. Journal of Neuroscience, 33(4), 1684–1695.

93. Zhou, S., Masmanidis, S. C., & Buonomano, D. V. (2020). Neural sequences as an optimal dynamical regime for the readout of time. Neuron, 108(4), 651–658.

94. Kraus, B. J., Brandon, M. P., Robinson II, R. J., Connerney, M. A., Hasselmo, M. E., & Eichenbaum, H. (2015). During running in place, grid cells integrate elapsed time and distance run. Neuron, 88(3), 578–589.

95. Heys, J. G., & Dombeck, D. A. (2018). Evidence for a subcircuit in medial entorhinal cortex representing elapsed time during immobility. Nature Neuroscience, 21(11), 1574–1582.

96. Tsao, A., Sugar, J., Lu, L., Wang, C., Knierim, J. J., Moser, M. B., & Moser, E. I. (2018). Integrating time from experience in the lateral entorhinal cortex. Nature, 561(7721), 57–62.

97. Rybakken, E., Baas, N., & Dunn, B. (2019). Decoding of neural data using cohomological feature extraction. Neural Computation, 31(1), 68–93.

98. Chaudhuri, R., Gerçek, B., Pandey, B., Peyrache, A., & Fiete, I. (2019). The intrinsic attractor manifold and population dynamics of a canonical cognitive circuit across waking and sleep. Nature Neuroscience, 22(9), 1512–1520.

## Methods references

99. Donato, F., Jacobsen, R. I., Moser, M. B., & Moser, E. I. (2017). Stellate cells drive maturation of the entorhinal-hippocampal circuit. Science, 355(6330), eaai8178.

100. Low, R. J., Gu, Y., & Tank, D. W. (2014). Cellular resolution optical access to brain regions in fissures: imaging medial prefrontal cortex and grid cells in entorhinal cortex. Proceedings of the National Academy of Sciences, 111(52), 18739–18744.

101. Paxinos, G., & Franklin, K. B. (2019). The Mouse brain in Stereotaxic Coordinates. Academic press.

102. Stringer, C., Pachitariu, M., Steinmetz, N., Reddy, C. B., Carandini, M., & Harris, K. D. (2019). Spontaneous behaviors drive multidimensional, brainwide activity. Science, 364(6437), eaav7893.

103. Van Der Maaten, L., Postma, E., & Van den Herik, J. (2009). Dimensionality reduction: a comparative review. J Mach Learn Res, 10(66-71), 13.

104. McInnes, L., Healy, J. & Melville, J. UMAP: Uniform manifold approximation and projection for dimension reduction Preprint at https://arxiv.org/abs/1802.03426 (2018).

105. Connor Meehan, Jonathan Ebrahimian, Wayne Moore, and Stephen Meehan (2022). Uniform Manifold Approximation and Projection (UMAP) (https://www.mathworks.com/matlabcentral/fileexchange/71902), MATLAB Central File Exchange.

106. Berens, P. (2009). CircStat: a MATLAB toolbox for circular statistics. Journal of Statistical Software, 31, 1–21.

107. Shannon, C. E. (1948). A mathematical theory of communication. The Bell System Technical Journal, 27(3), 379–423.

108. Cover, T.M., and Thomas, J. A. (2006). Elements of Information Theory. New Jersey, NJ: Wiley.

109. Panzeri, S., Senatore, R., Montemurro, M. A., & Petersen, R. S. (2007). Correcting for the sampling bias problem in spike train information measures. Journal of Neurophysiology, 98(3), 1064–1072.

110. Meshulam, L., Gauthier, J. L., Brody, C. D., Tank, D. W., & Bialek, W. (2019). Coarse graining, fixed points, and scaling in a large population of neurons. Physical Review Letters, 123(17), 178103.

